# A regulatory region that controls Wnt gene expression following tissue injury is required for proper muscle regeneration

**DOI:** 10.1101/2025.10.01.679323

**Authors:** Catriona Y. Logan, Xinhong Lim, Matt Fish, Makiko Mizutani, Brooke Swain, Roel Nusse

## Abstract

The capacity to detect and respond to injury is critical for the recovery and long-term survival of many organisms. Wnts are commonly induced by tissue damage but how they become activated transcriptionally is not well understood. Here, we report that mouse *Wnt1* and *Wnt10b* are induced following injury in both lung and muscle. These Wnts occupy the same chromosome and are transcribed in opposite directions with 12kb between them. We identified a highly conserved cis-acting regulatory region (enhancer) residing between *Wnt1* and *Wnt10b* that, when fused to a LacZ reporter, is activated post-injury. This enhancer harbors putative AP-1 binding sites that are required for reporter activity, a feature observed in other injury-responsive enhancers. Injured muscles in mice carrying a germ-line deletion of the enhancer region display reduced *Wnt1* and *Wnt10b* expression and show elevated intramuscular adipogenesis--a hallmark of impaired regenerative capacity—revealing a requirement of this enhancer for proper regeneration. Enhancer redundancy is common in development, but our in vivo analysis shows that loss of a single injury-responsive regulatory region in adult tissues can produce a detectable regenerative phenotype.

**Summary:** A new, previously unknown shared regulatory region residing between two Wnts, *Wnt1* and *Wnt10b*, is induced by tissue damage and required for muscle regeneration.

## Introduction

The ability of an organism to regenerate or repair tissues following injury requires that a tissue can detect damage and deploy mechanisms that facilitate the restoration of tissue architecture. Central to this process is the transcriptional activation of genes, mediated by regulatory sequences termed ‘enhancers.’ Many cell signals that regulate embryonic development are induced following tissue injury and re-used in adults to drive regeneration or repair (Maddaluno et al., 2017; Aros et al., 2021; Fazilaty and Basler, 2023). Regulatory sequences that promote developmental expression of some of these signals have been characterized, but how tissue damage is sensed and integrated at the level of enhancers following injury, particularly for cell-to-cell signaling molecules, is not well understood.

Wnts are a family of developmental signals that also function in both normal adult tissue homeostasis and disease (Nusse and Clevers, 2017). Additionally, Wnt genes are commonly induced following injury in a wide variety of organisms to facilitate repair, regeneration, or re-patterning (Whyte et al., 2012) and may even confer regenerative capacity (Campos, 2025). The induction of Wnts post-injury is often rapid (McClure et al., 2008; Petersen and Reddien, 2009; Fernandez-Martos et al., 2011; Vizcaya-Molina et al., 2018; Cazet et al., 2021) suggesting that Wnts are well-positioned as cell-cell signaling factors to integrate injury signals and transition cells from a homeostatic to regenerative state. Only a few studies have explored the transcriptional regulation of Wnts, with most focused on developing embryos (Echelard et al., 1994; Danielian et al., 1997; Rowitch and and McMahon, 1998; Park et al., 2012; O’Brien et al., 2018; Lekven et al., 2019; van de Grift et al., 2025). In *Drosophila*, an injury-responsive Wnt enhancer that drives *Wingless* and *Wnt6* in early larval wing discs has been described (Harris et al., 2016), but how Wnts are activated following injury in adult tissues and particularly in vertebrates is unknown.

Multiple types of injury responsive enhancers have now been identified. Some display a dual function, utilized during both development and regeneration (Huang et al., 2012), while others are largely dedicated to damage responses (Guenther et al., 2015; Heller et al., 2022). Other enhancers can be separated into different tissue-specific domains or modules (Kang et al., 2016), display temporal regulation, becoming silenced as tissues mature (Harris et al., 2016; Harris et al., 2020), or possess distinct initial injury-sensing and regeneration-deploying activities, with important functional consequences that may predict whether a species can regenerate or not (Wang et al., 2020). Recently, an enhancer that drives one gene under normal conditions that can be re-purposed to drive another gene upon injury has been described (Rao et al., 2025). These examples reflect the diversity and complexity of regulatory sequences that control injury-responsive gene activation and highlight the need for deeper characterization of their dynamics, function, and logic.

Additionally, to what degree injury-responsive enhancers are unique vs. redundant, and to what extent mutations in injury responsive enhancers negatively affect tissue restoration are not well understood. A requirement for injury-responsive enhancers have been demonstrated in several examples (Hewitt et al., 2017; Soukup et al., 2019; Wang et al., 2020; Sun et al., 2022; Gracia-Latorre et al., 2022; Zlatanova et al., 2023), but defects can be subtle or hard to detect (Sun et al., 2022). Understanding how impaired enhancer function or variations in enhancer sequences might predispose one towards poor tissue healing and function over time (Zaugg et al., 2022), has important implications for human health.

In this study, we investigated how Wnts are transcriptionally activated in adult tissues following damage. We report the identification of a regulatory region residing between *Wnt1* and *Wnt10b* that in vivo responds to tissue injury and is required for proper regeneration of damaged muscle.

## Results

### *Wnt1* and *Wnt10b* are up regulated in response to tissue injury

To ask how injury is sensed by Wnt genes and to identify those genes induced by injury, we performed an mRNA in situ hybridization screen in four different injury models for all 19 Wnt genes (Fig. S1A-D, Table S1). Treatments included Naphthalene administration (lung) (Hong et al., 2001), Barium Chloride (BaCl2) injection (muscle) (Wosczyna et al., 2019), Carbon Tetrachloride (CCl4) exposure (liver) and Streptozotocin injury (pancreas) (Furman, 2021).

All tissue injuries triggered expression of multiple Wnts (Figure 1A-D, Fig. S1A-D, Table S1). Among those induced, *Wnt1* and *Wnt10b* were observed in two specific contexts--following BaCl2 injection in the Tibialis Anterior (TA) muscle (Figure 1A-G, Fig. S1B), and Naphthalene injection in the lung (Fig. S1A, S2A,B). BaCl2 injection is an acute chemical injury model in which muscle fibers degenerate and then regenerate over 2 weeks (Hardy et al., 2016). Naphthalene injection is an acute airway injury model in which damaged conducting airway rapidly regenerates over a similar time frame (Mahvi et al., 1977; Stripp et al., 1995; Van Winkle et al., 1995). Both in lung and muscle, uninjured tissue initially displayed low *Wnt1* and *Wnt10b* transcripts, but injury induced *Wnt1* and *Wnt10b* mRNA signals, with expression continuing for several days (Figure 1A,B, Fig. S2A,B). Quantifying *Wnt1* and *Wnt10b* positive nuclei in muscle confirmed elevation of mRNA over 1-3 days post-injury, although the temporal dynamics differed between the Wnts (Figure 1C,D). These data demonstrate that Wnt genes are induced by injury in a variety of mouse tissues (Figure 1, Fig. S1A-D, Fig. S2, Fig. S3), and *Wnt1* and *Wnt10b* are two specific Wnts that become expressed in more than one injury context.

**Figure 1:**
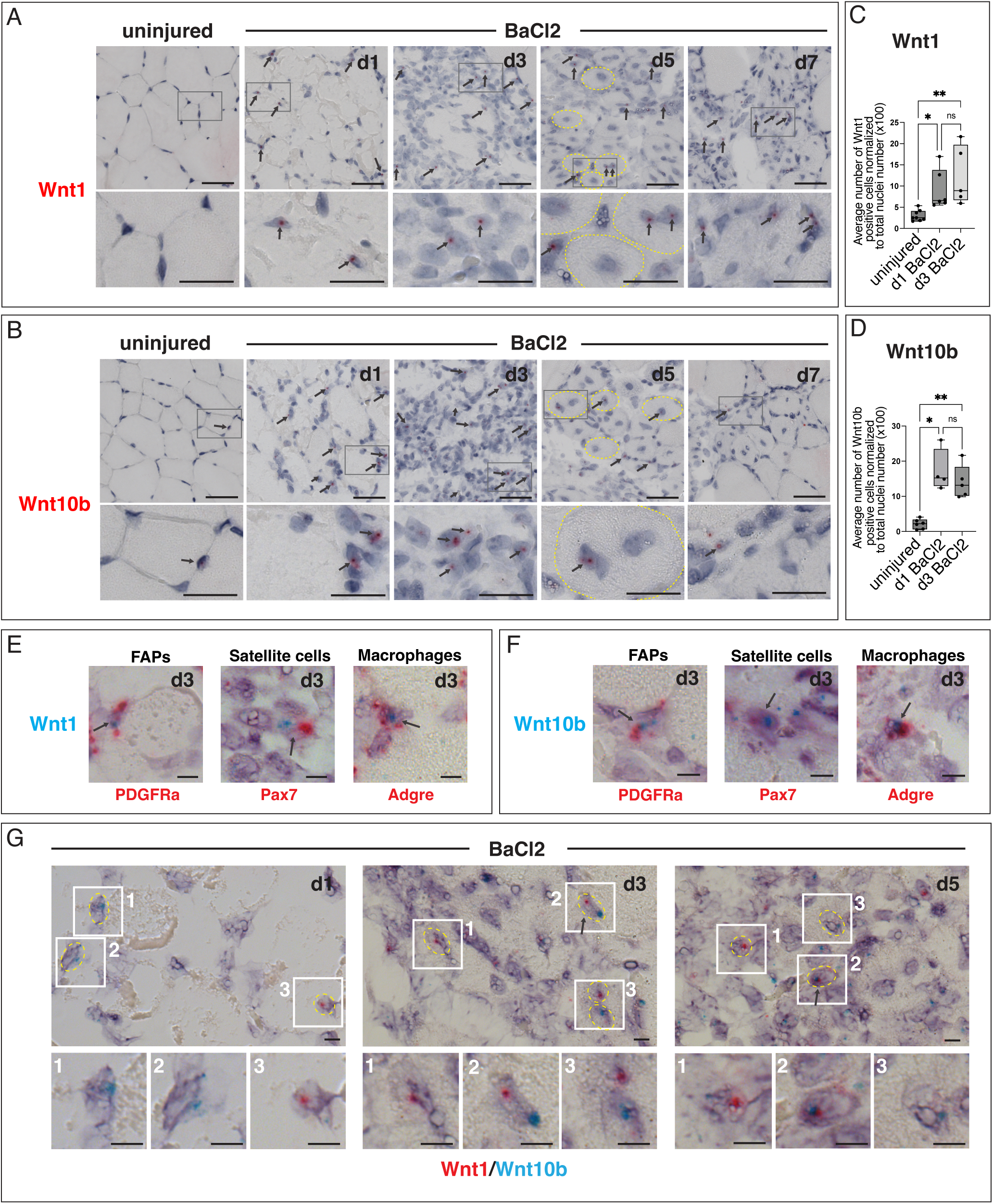
*Wnt1* and *Wnt10b* are induced by tissue injury following BaCl2 injection. (A, B) Wnt expression post-BaCl2 induced injury in skeletal muscle as assessed by RNAscope mRNA in situ hybridization. (A) *Wnt1* and (B) *Wnt10b* expression in TA muscle following BaCl2 injection at 1, 3, 5 and 7 days post-injury. Uninjured contralateral muscle at 1 day is shown for comparison. *Wnt1* and *Wnt10b* positive cells are infrequently observed in uninjured muscle. Transcripts appear in single cells post-injury and then in nascent myofibers which display centrally located nuclei. Boxes mark regions that are enlarged under each panel. Arrows point to red mRNA signals. Yellow dotted lines outline myofibers. (C, D) Quantification of *Wnt1* and *Wnt10b* expression at days 1 and 3 post-injury. (C) Number of *Wnt1* positive nuclei increases significantly at 1 and 3 days post-injury compared to uninjured contralateral muscles. (*P<0.05, **P<0.005, One-way ANOVA, Kruskal-Wallis test) n=8 uninjured consists of n=4 d1 uninjured, n=4 d3 uninjured. n=6 d1 BaCl2, n= 5 d3 BaCl2. (D) Number of *Wnt10b* positive nuclei increases significantly at 1 and 3 days post-injury compared to uninjured contralateral muscles. (*P<0.05, **P<0.005, Brown-Forsythe and Welch one-way ANOVA tests) n=6 uninjured consists of n=3 d1 uninjured, n=3 d3 uninjured, n=4 d1 BaCl2, n=5 d3 BaCl2. (E, F) Duplex mRNA in situ hybridization showing *Wnt1*(E) and *Wnt10b*(F) signals in *PDGFRa*, *Pax7*, and *Adgre* positive cells that mark the FAPs, satellite cells, and macrophages respectively at 3 days post-inury. Arrows point to examples of double-positive nuclei. n=2 animals (G) Duplex mRNA in situ hybridization showing that *Wnt1* and *Wnt10b* can be colocalized. Representative examples from 1-, 3-, and 5-days post-injury are shown. White numbered boxes mark regions magnified in the row below each image. Yellow dotted lines mark individual nuclei. Arrows mark cells scored as *Wnt* double-positive. Graphs show mean ± s.d. (Scale bar: (A, B) 20 um; (E-G) 5 um)

For comparison, we examined *Wnt5a* post-BaCl2 injury, because it is found in muscle and induced by tissue damage (Polesskaya et al., 2003; Reggio et al., 2020). *Wnt5a* displayed transcripts at similar abundance and intensity to *Wnt1* and *Wnt10b*, with *Wnt5a* expression gradually increasing between 1-3 days post-injury (Fig, S3A,B). These data show that *Wnt1* and *Wnt10b* staining levels overall, are typical for expression of Wnts detected in injured muscle tissue (Figure 1A,B, Fig. S3A).

We asked which cells express *Wnt1* and *Wnt10b* in muscle. Previous studies show that Wnts are predominantly found in Fibro-adipogenic Progenitors (FAPs) (Joe et al., 2010; McKellar et al., 2021) scmuscle.bme.cornell.edu/), one of the stromal cell populations that regulates muscle regeneration (Joe et al., 2010; Wosczyna et al., 2019), with low-level expression in other cell types (Reggio et al., 2020). Using 2-plex in situ hybridization, we observed that a subset of *Wnt1* (Figure 1E) and *Wnt10b* (Figure 1F) positive cells are FAPs, marked by *PDGFRa*. A subset of Wnt-positive cells also expresses the satellite cell marker *Pax7*, and *Adgre*, a marker predominantly expressed in macrophages (Figure 1E,F). Double-positive nuclei expressing *Wnt/PDGFRa* or *Wnt/Pax7* are rare, comprising less than 15% of the *Wnt1* and *Wnt10b*-expressing cells (Fig. S4A,B). Similarly, *Wnt/Adgre* double-positive cells never constituted more than 25% of Wnt-positive cells. (Fig. S4A,B). Because not all FAPs express *PDGFRa*, and FAPs can shuttle between dynamic cell states (Oprescu et al., 2020), we also examined co-expression of *Wnt1* and *Wnt10b* with other FAP markers (*Osr1, Wisp1, Dpp4, Dlk1* and *CxCl14*). *Wnt1* and *Wnt10b* co-staining with these FAP markers was also rare but detectable, showing that both Wnts are found in multiple FAP sub-populations (Fig. S4A,B,D). We may have under-counted double-positive cells because a strongly expressed transcript masks a more weakly expressed one when stained by non-fluorescent dual in situ hybridization. We also counted cells as double-positive only if their nuclei could clearly be distinguished from neighboring cells. We do not know whether *Wnt1/Wnt10b* expression post-injury in multiple cell types reflects a general role for Wnt signals in response to tissue damage, or if each Wnt plays distinct functions within each cell population.

We also asked if *Wnt1* and *Wnt10b* were co-expressed in the same cells. *Wnt1* and *Wnt10b* double-positive nuclei are observed but their numbers are low at 1, 3, and 5 days post-injury (Fig. S4C). Most nuclei display one or the other signal, but not both (Figure 1G). Since the first few days post-injury are highly dynamic, non-overlapping Wnt staining could be due to differences in the dynamics of promoter engagement with enhancers, mRNA perdurance, and expression of these genes in distinct cell sub-populations. Despite low abundance, we estimated that at least ∼10% of cells were *Wnt1* and *Wnt10b* double-positive at any given timepoint (Fig. S4C) suggesting that a common regulatory element might drive expression of these two Wnts.

### Injury-responsive regulatory elements reside between Wnt1 and Wnt10b

In the Zebrafish, *Wnt1* and *Wnt10b* are coordinately expressed through shared developmental enhancers (Lekven et al., 2019). Given the expression of *Wnt1* and *Wnt10b* in damaged adult lung and muscle, we asked if these genes might share regulatory elements post-injury in the mouse. *Wnt1* and *Wnt10b* reside on chromosome 15, are transcribed in opposite directions, and are separated by 12kb which provided a small, tractable interval to search for putative injury-responsive regulatory sequences. We identified 3 peaks of high conservation in the intergenic region between *Wnt1* and *Wnt10b* (Figure 2A) and hypothesized that they might regulate the injury-responsive expression of these Wnts.

**Figure 2:**
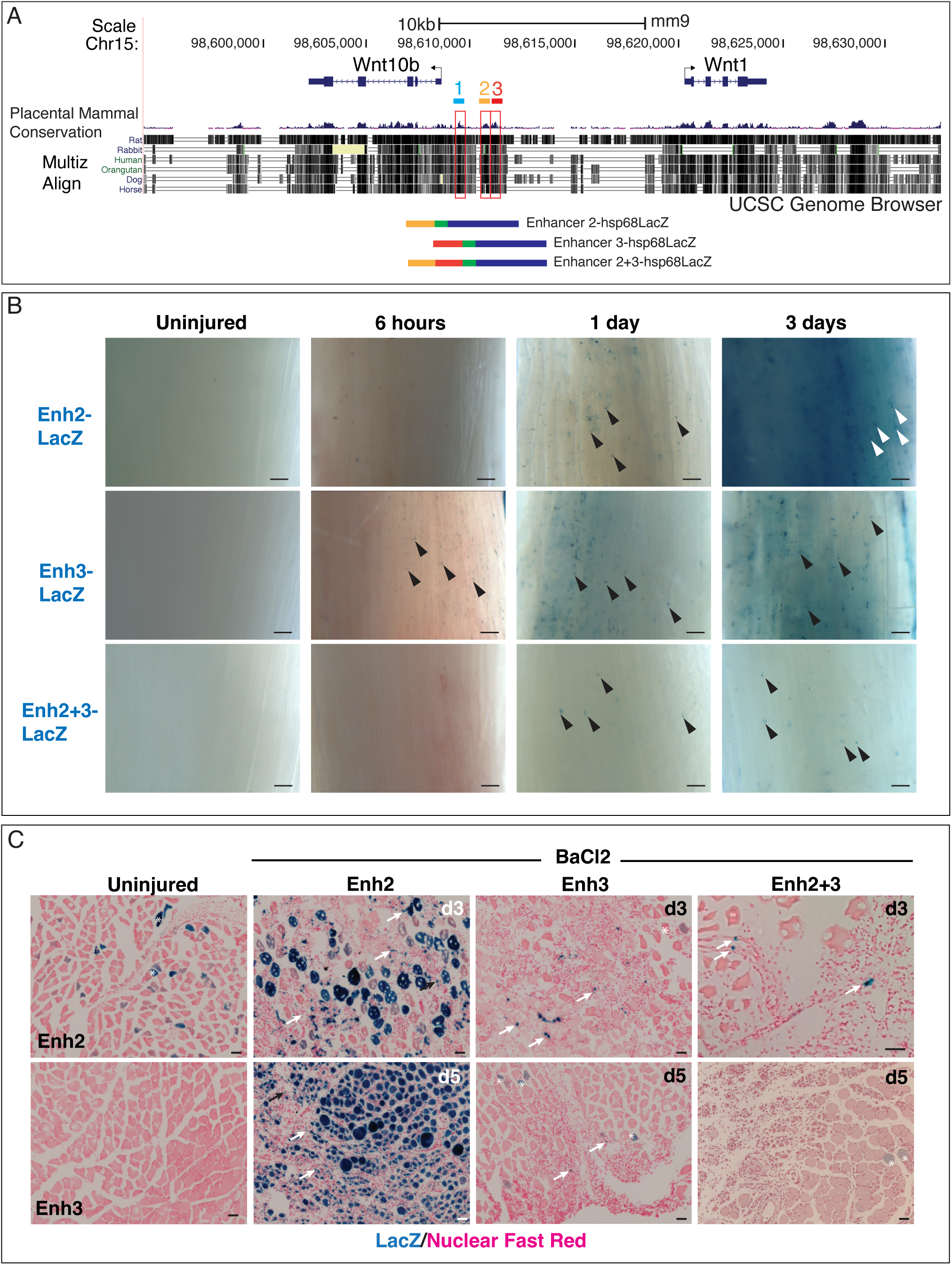
Regulatory sequences residing between *Wnt1* and *Wnt10b* are injury responsive. (A) Data from the UCSC Genome Browser shows that *Wnt1* and *Wnt10b* reside on the same chromosome and are transcribed in opposite directions (arrows). Three highly conserved sequences between *Wnt1* and *Wnt10b* are labeled 1 (light blue), 2 (yellow), 3 (red). Multiple genome alignment tracks (Multiz Align) and conserved peaks from placental mammal alignments are shown. Below, schematics of constructs containing putative Enhancers 2, 3, and 2+3 fused to the *LacZ* gene are provided. (B) BaCl2 injury of adult reporter mice containing Enh2, Enh3, and Enh2+3-LacZ reveal that all three sequences drive reporter expression following muscle injury. Whole mount muscles are shown. Uninjured contralateral muscles at 6 hours are provided for comparison. Arrowheads mark examples of single cells that express the reporter. (C) Sectioned tissues from LacZ stained BaCl2 injured TA muscles from Enhancer 2, 3, and 2+3 reporter mice at 3-5 days. Uninjured muscles from Enhancer 2 and 3 reporters are shown. Arrows mark examples of single cells that express the reporter. Some background staining is observed (asterisks), but staining is variable and sporadic. (Scale bar: 20 um)

We selected a 6kb region that encompassed the three most prominent conserved peaks for further analysis. This 6kb sequence and sequences representing each individual peak were fused to a LacZ reporter construct containing a basal *Hsp68* promoter (Kothary, 1989) (Figure 2A, see Fig. S5A for all constructs). Multiple adult transgenic lines carrying independent insertions for each construct were produced.

Initially all lines were tested in the lung. Reporter activity was assessed at 5 days post-Naphthalene injection, as restoration of the damaged airway is not fully complete at this timepoint and we could verify that injury had occurred (Zemke et al., 2009; Volckaert et al., 2011). The 6kb region (n=3/3 independent lines), as well as Enhancers 2 (Enh2, n=2/2 independent lines), 3 (Enh3, n=5/5 independent lines) and 2+3 (Enh2+3, n=3/4 independent lines) induced LacZ activity following injury in damaged airways. We could not ascertain the injury-responsiveness of Enhancer 1, as only two adult lines were obtained, one which responded to injury and one that did not (n=1/2 independent lines) (Fig. S5B, Table 1). Vehicle-injected animals did not display reporter expression in airways (Fig. S5B). LacZ staining was occasionally observed in non-airway cell types, but patterns were inconsistent between independently generated lines, suggesting that expression was due to reporter insertion into a transcriptionally active genomic locus but not reflective of enhancer-specific activity. Non-transgenic animals lacked staining (not shown) and the basal promoter alone fused to *LacZ* (‘No Enhancer’) produced sparse reporter activity (Fig. S5B, Table 1).

**Table 1:**
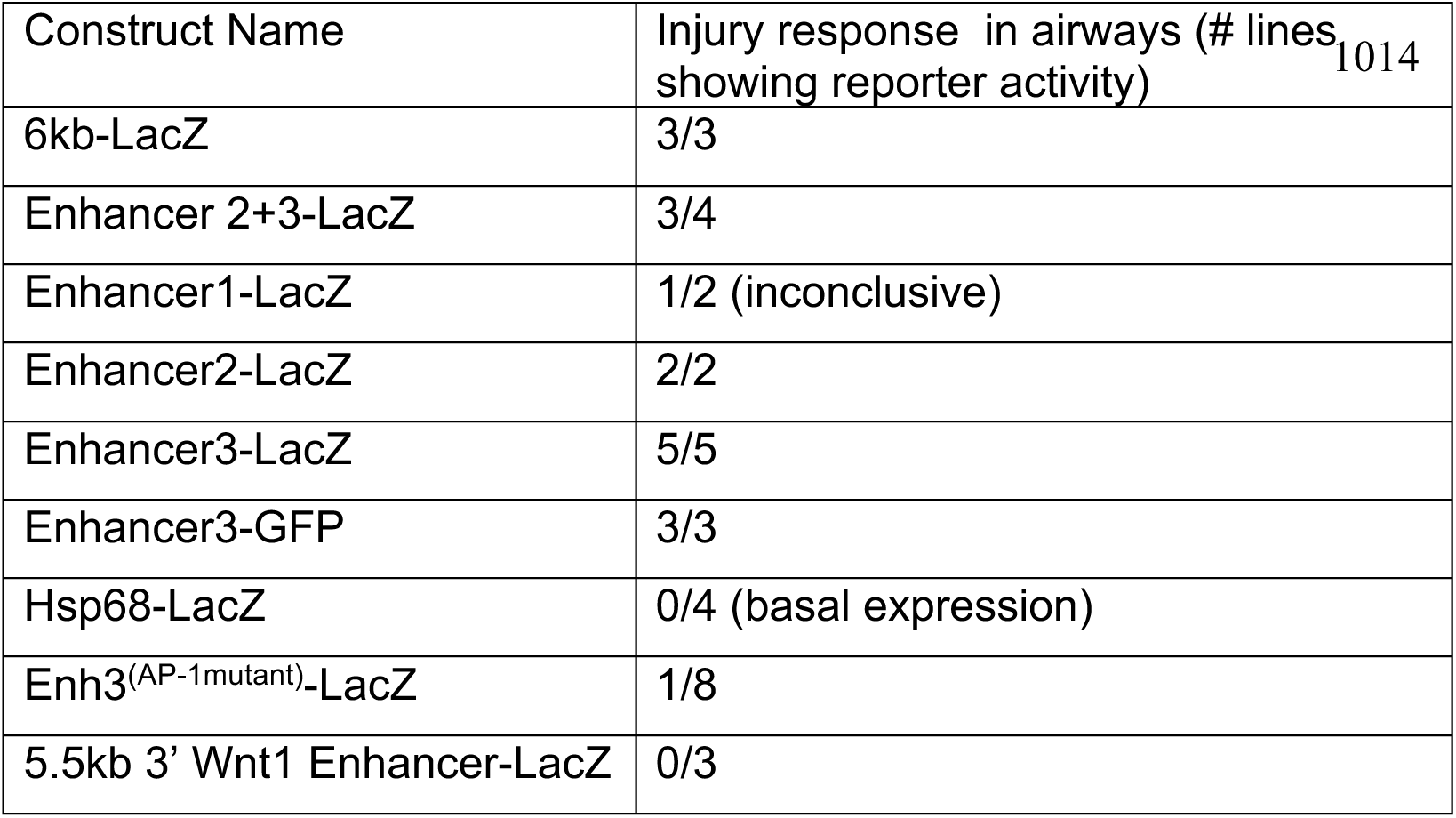
Reporter activity in transgenic mouse lungs at 5 days post-Naphthalene injury.

Finally, a Wnt1-LacZ mouse carrying a 5.5kb 3’ regulatory region that drives embryonic *Wnt1* in the central nervous system (CNS) expression (Echelard et al., 1994) was examined in injured lung (Fig. S5B, Table 1). This reporter failed to display LacZ activity (n=0/3 embryos), showing that this well-known embryonic *Wnt1* regulatory sequence is not activated following injury.

From these data, we conclude that the 6kb region residing between *Wnt1* and *Wnt10b* harbors injury-responsive elements, with Enhancers 2 and 3 showing injury-responsive activity.

### Injury-responsive enhancers 2 and 3 become expressed in damaged muscle

Wnts and Wnt signaling are well-studied in developing muscle (von Maltzahn et al., 2012; Girardi and Le Grand, 2018) and play roles in muscle regeneration (Polesskaya et al., 2003; Brack et al., 2008; Murphy et al., 2014; Huraskin et al., 2016; Reggio et al., 2020; Gurriaran-Rodriguez et al., 2024; Kamizaki et al., 2024). Because *Wnt1* and *Wnt10b* transcripts are induced by muscle injury (Figure 1) and *Wnt10b* mutants display a regeneration phenotype in injured muscles (Vertino et al., 2005), we reasoned that Enhancers 2 and/or 3 might regulate *Wnt1/Wnt10b* in muscle tissue. One representative mouse line of Enh2-LacZ (Enh2-LacZ) and Enhancer 3-LacZ (Enh3-LacZ) and Enhancer 2+3-LacZ (Enh2+3-LacZ) was selected for all subsequent experiments.

Following BaCl2 injury, Enh2-LacZ and Enh3-LacZ are activated. As early as 6 hours, Enh3-LacZ displays staining in small, scattered cells that are visible as speckles in whole mount muscle tissue, followed by speckled Enh2-LacZ staining at 24 hours post-injury (Figure 2B). Enh3-LacZ is similarly rapidly induced before Enh2-LacZ in injured lung (Fig. S6). By 3 days, BaCl2-injured Enh2 reporter tissues display very dark *LacZ* staining in whole-mounts (Figure 2B) which in section appear as blue regenerating myofibers as well as small individual cells (Figure 2C, arrows). Enh3 reporter mice exhibit only small cells near injured myofibers at days 3-5 (Figure 2C, arrows) and staining in nascent myofibers is not observed.

We examined the Enh2+3-LacZ reporter to assess the combined activity of Enhancers 2 and 3 in injured muscle. Speckled staining begins to appear in Enh2+3-LacZ mice at 1 day post-injury in whole-mounts and continues through 3 days (Figure 2B, arrowheads), although expression appears weaker compared to Enhancers 2 or 3 reporters alone (Fig. 2B). In sections by day 3, single cells next to myofibers are observed, but no expression in regenerating myofibers is seen (Figure 2C). By day 5, virtually no reporter expression is detected (Figure 2C).

Uninjured muscles display no reporter activity (Figure 2B,C), although sporadic myofiber staining can sometimes be observed (Figure 2C, asterisks). This staining has been seen in other contexts (Guenther et al., 2015), possibly resulting from basal activity of the Hsp68-LacZ construct and is considered background. Non-transgenic muscles display no LacZ staining (data not shown). Table S2 summarizes reporter expression observed in Enh2-LacZ, Enh3-LacZ, and Enh2+3-LacZ muscles. Together, these data suggest that Enhancers 2 and 3 likely integrate different sets of transcription factor inputs as cells detect and respond to injury over time. However, the main function of the Enh2+3 region may be to promote up-regulation of Wnt gene expression in single cells that reside between damaged myofibers.

We asked if our reporters recapitulate *Wnt1* and *Wnt10b* gene expression. Double in situ hybridization for *LacZ* and Wnt resulted in all cells displaying *LacZ* expression (data not shown), likely due to the multimerized nature of transgenic insertions (Bishop, 1996). As an alternative approach, because the Enh2 and 3-LacZ reporters display two phases of expression in which b-gal staining is first found in single cells that reside between myofibers and then in regenerating myofibers from day 3 onwards (Figure 2B,C), we asked whether *Wnt1* and *Wnt10b* also display similar expression patterns. *Wnt1* and *Wnt10b* transcripts are visible in cells residing between injured myofibers at d1, with no expression found within the myofibers themselves (Figure 1A,B). By day 5, Wnt expression is observed in regenerating myofiber nuclei (Figure 1A,B), reminiscent of the strong Enh2-LacZ staining observed in nascent myofibers (Figure 2B,C). Given that Enh2+3 does not express in myofibers during regeneration (Fig. 2B,C), we do not know if Enh2 regulates Wnt expression in myofibers with perhaps other enhancers, or if myofiber expression is controlled independently by other regulatory elements. Nevertheless, the appearance of *Wnt1* and *Wnt10b* in isolated single cells post-injury and the similar appearance of single LacZ-positive cells in injured muscles, suggests that these enhancers could drive the initial phase of injury-responsive Wnt expression.

We asked if publicly available data supports the idea that Enh2 and Enh3 function as enhancers. UCSC Genome Browser data shows that both sequences display distal enhancer-like signatures (DNAse1-hypersensitivity (Fig. S7A) in uninjured mouse lung and muscle, and H3K27 acetylation marks (Fig. S7B)). A GEO dataset in which ATAC-seq was performed on muscle stem cells following BaCl injury at 1, 16, 32 and 60 hours (GSE189044, (Dong et al., 2022)), also shows peaks at Enh2 and Enh3 (data not shown). *Wnt1* and *Wnt10b* may also reside within a Topologically Associated Domain (TAD), with contact points possible between their promoters and Enh2 and Enh3 (Fig. S7C). None of the publicly available data match our experimental conditions exactly, but these findings suggest that Enhancers 2 and 3 could function as regulatory sequences.

### Putative AP-1 binding sites are required for the injury-response by Enhancer 3

The rapid induction of Enh3-LacZ in injured muscle (Figure 2B) and lung (Fig. S6) suggests that this regulatory sequence might be particularly responsive to the initial stress of tissue damage. To ask which transcription factors activate the Enh3 reporter, we performed a bioinformatic analysis which identified two putative AP-1 sites (mm9 mouse genome alignment, sequence TGAGTCA; AP-1 consensus sequence is TGAG/CTCA). One site was identified using GREAT (McLean et al., 2010) and the other was identified by TFSEARCH (http://diyhpl.us/∼bryan/irc/protocol-online/protocol-cache/TFSEARCH.html).

If AP-1 binding regulates Enh3, we hypothesized that it might be highly responsive to a wide variety of injuries. We observed reporter activation in multiple injury contexts, including airway and tracheal damage induced by Naphthalene (Hsu et al., 2014), liver injury by CCl4 (Weber et al., 2003), skin injury by biopsy punch (Ansell et al., 2014), and heart tissue subjected to myocardial infarction (Kolk et al., 2009)(Figure 3B). Uninjured tissues (Figure 3B) and injured non-transgenic tissues (Fig. S8) display no b-gal activity. In the muscle, in situ hybridization revealed several AP-1 family members, *FosL1*, *FosL2*, *c-fos* and *fos-b*, that were dynamically expressed between 4-24 hours post-injury (Fig. S9). These data show that Enh3 is highly injury-responsive and suggest that immediate-early genes such as AP-1 may induce early and rapid activation of Enh3 post-injury.

**Figure 3:**
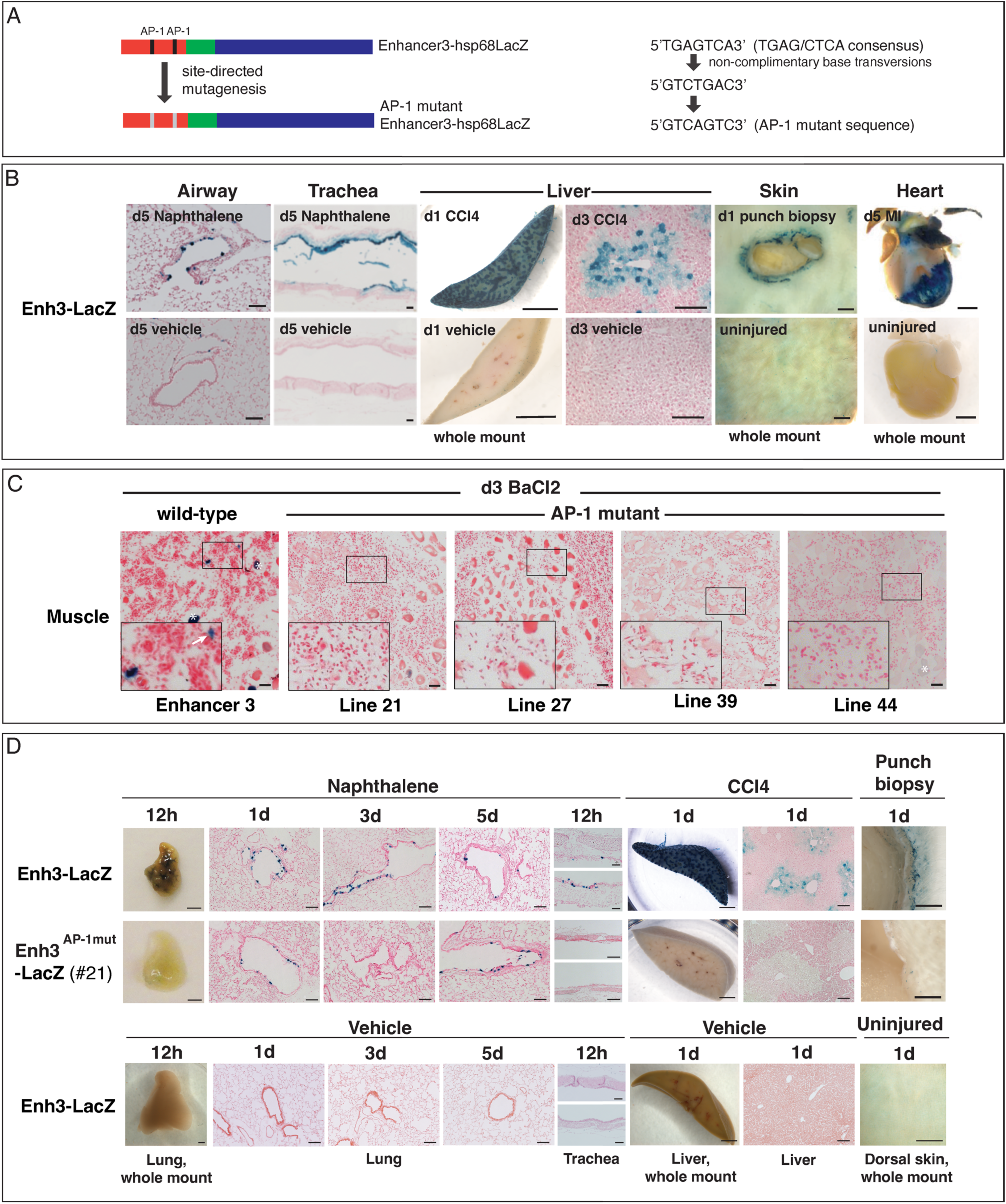
AP-1 binding sites are required for Enhancer 3 reporter activity in response to injury. (A) Schematic of Enh3-LacZ showing two putative AP-1 binding sites and a mutated version in which the AP-1 binding sites are altered. Predicted AP-1 binding sequences found in Enhancer 3 (TGAGTCA), the predicted sequence from non-complimentary base transversions (GTCTGAC), and the final mutant sequence (GTCAGTC) are shown. (B) X-gal staining in multiple injured tissues from Enhancer 3 reporter mice. Uninjured tissues are shown below. Naphthalene injury in airways and trachea at 5 days, CCl4 injection in the liver at 1 day (as whole mount) and 3 days (in tissue section), punch biopsy of the skin at 1 day post-injury, and myocardial infarction (MI) of the heart at 5 days all produce robust Enh3-LacZ activation. Uninjured Enh3-LacZ tissues do not stain with X-gal. (C) Injured muscle from wt Enh3-LacZ compared to AP-1 mutant Enh3-LacZ lines. 4 different mutant lines are shown. Boxes outline areas magnified in the lower left of the image. Arrow marks positive Enh3-LacZ signal. Asterisks mark background myofiber staining. (D) LacZ staining in lung, liver and skin to compare wt Enhancer 3 reporter activity vs. a representative AP-1 mutant line (line #21 is shown). The wild-type Enhancer 3 reporter displays X-gal staining in all tissue injury contexts. The mutant reporter displays low expression that more closely resembles uninjured wt Enh3 tissue (see Fig. S11 for more AP-1 mutant lines). (Scale bar: (B, D) 100 um lung, trachea, liver sections; 2 mm lung, liver, skin whole mounts; 1 mm heart; (C) 20 um)

To directly test whether predicted AP-1 binding sites are required for Enh3 injury-responsiveness, we performed site-directed mutagenesis to alter the AP-1 consensus TGAG/CTCA sequences to GTCAGTC (Figure 3A). Interestingly, in multiple genome alignments, only the house mouse carries two putative AP-1 binding sequences (Fig. S10A-C) and only one of the two predicted sites is highly conserved. We proceeded to mutate both sites to assess a requirement for AP-1 sequences in mouse Enh3, but only the conserved site could be involved in injury sensing.

Multiple, independently generated stable adult Enh3 reporter lines carrying the mutant AP-1 sites fused to Hsp68-LacZ were generated. In the muscle, injured tissues from wild-type (wt) Enh3 reporter mice displayed strong LacZ staining in single cells residing between damaged myofibers, but AP-1 mutant reporters failed to show LacZ staining (Figure 3C). Occasionally, non-specifically stained single myofibers were observed (Figure 3C, line 44, asterisk).

Similarly, following injury in the lung, liver, and skin, wt Enh3-LacZ displayed strong staining in damaged tissues (Figure 3D), but mutant AP-1 reporter lines failed to show robust injury-responsive reporter activity, resembling staining more reminiscent of uninjured wt Enh3-LacZ (Figure 3D). Figure 3D shows one representative example out of n=8/8 AP-1 mutant lines displaying reduced staining (see Fig. S11 for more lines, Table S3). Uninjured mutant AP-1 lines display no reporter activity (Fig. S11 for representative example). Together, these data show that the putative AP-1 site(s) in Enhancer 3 are required for injury-responsive reporter activation.

### Deletion of Enhancers 2 and 3 results in loss of *Wnt1* and *Wnt10b* expression

To ask if Enhancers 2 and 3 are required in vivo, we deleted the Enh2+3 regulatory region using CRISPR-Cas9. Two separate mouse lines, named Δ1 and Δ2, were generated, in which the Δ1 deletion (1209bp) was 206bp larger than Δ2 (1003bp) (Figure 4A,B). Δ1/+ and Δ2/+ mice were crossed to each other to produce Δ1/Δ2 mice (Figure 4B). We reasoned that this crossing scheme would minimize the chance of generating loci homozygous for deleterious alterations to the genome due to non-specific effects of CRISPR-Cas9-mediated genome editing.

**Figure 4:**
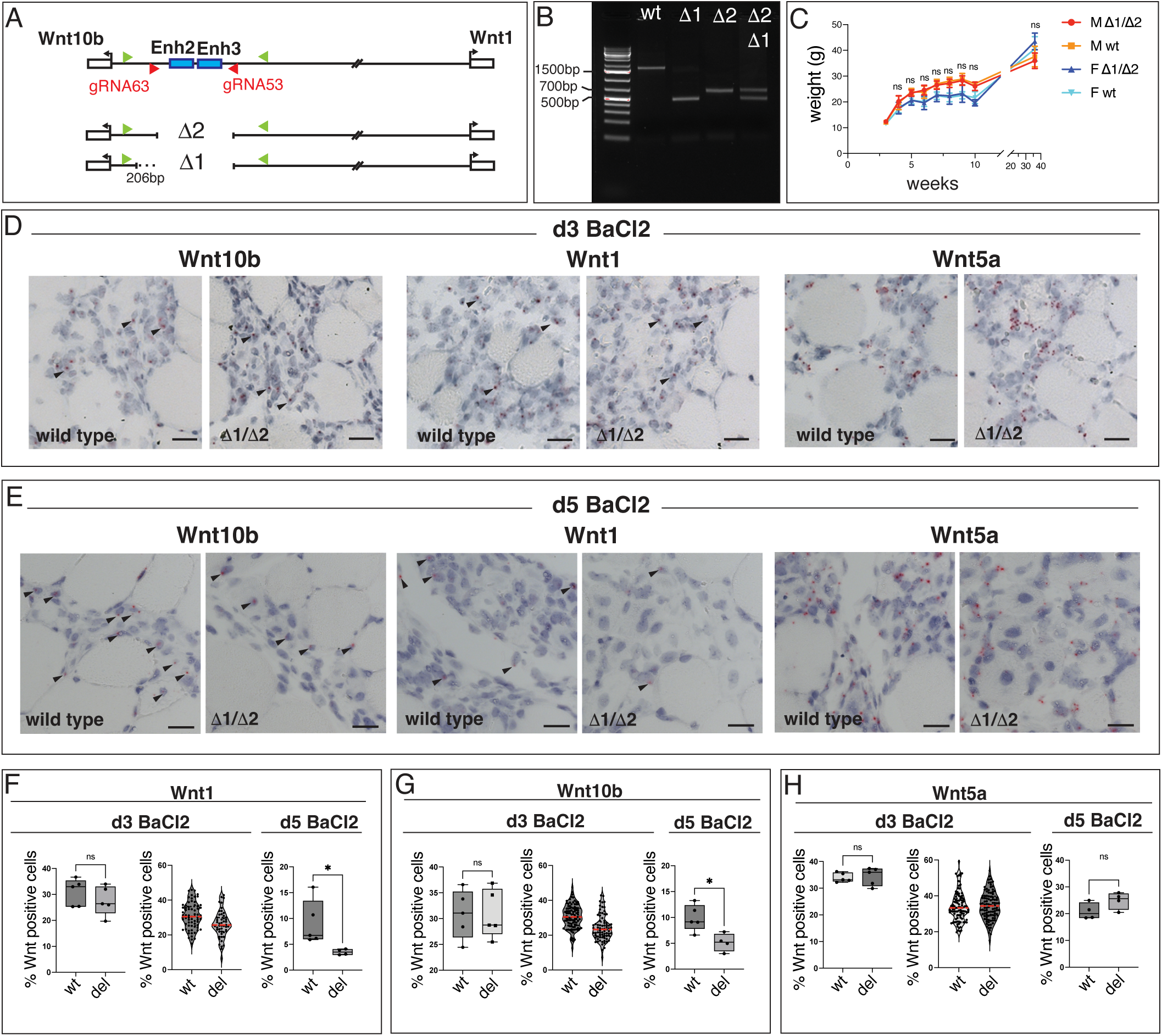
Loss of Enhancers 2 and 3 produces viable mice, but expression of Wnt1 and Wnt10b is reduced post-injury. (A) Schematic showing CRISPR-Cas9-mediated deletion of Enhancers 2 and 3. Guide RNAs (red arrowheads) and PCR genotyping primers (green arrowheads) are shown. Two independent mouse lines carrying Enhancer 2+3 deletions were generated, labeled Δ1 and Δ2. Δ1 is 206bp larger than Δ2. (B) PCR reactions from genotyping primers that flank the Enh2/3 region. Products sizes are 1.6kb(wt), 500bp(Δ1) and 700bp(Δ2). Δ1/Δ2 mice produce PCR bands that run as a doublet. (C) Δ1/Δ2 and wt mice display similar overall weights. Both males (M) and females (F) were followed between 2.5-35 weeks. (n= 9 wt M, n=9 Δ1/Δ2 M, n=11 wt F, n=10 Δ1/Δ2 F) (D-E) *Wnt1*, *Wnt10b*, and *Wnt5a* expression assessed by mRNA in situ hybridization in adult TA muscles. Arrowheads mark examples of transcripts. (D) Staining at 3 days post-BaCl2 injury. (n>3 mice). (E) Staining at 5 days post-BaCl2 injury. (n>3 mice) (F-H) Quantification of Wnt expressing nuclei at 3 and 5 days post-injury. Each dot on the box plots represents mean percents of Wnt-positive nuclei normalized to total number of nuclei counted per field for each mouse. Violin plots display all quantified fields. (F) A slight decrease in *Wnt1* positive cells is detected at 3 days followed by significant reduction at d5. (G) A slight decrease in *Wnt10b* positive cells is detected at 3 days, followed by significant reduction at d5. (H) Wnt5a expression shows no decline over time in Δ1/Δ2 muscles compared to wt. Whiskers in box plots represent the Min to Max.; red lines mark the median in violin plots (n=5 wt, n=5 del for *Wnt1*, *-10b* and *-5a* at d3; n=5 wt, n=4 del for *Wnt1*, *-10b* at d5; n=4 wt, n=4 del for *Wnt5a* at d5) (*Wnt1* d3 Mann-Whitney test; *Wnt1* d5 *p<0.05 Mann-Whitney test; Wnt10b d3 Unpaired t-test; *Wnt10b* d5 *p<0.05 Unpaired t-test; *Wnt5a* d3 Welch’s t-test; *Wnt5a* d5 Unpaired t-test; ns, not significant) (Scale bar: 20 um)

We obtained viable Δ1/Δ2 animals, but Mendelian ratios were slightly lower for this class than predicted when assessed at weaning age (Table 2; 25% expected; 19.44% observed). The reduced percentage of Δ1/Δ2 animals also predicted that the remaining classes, if all equally healthy, each should produce ratios of ∼26.85, but Δ1/+ and Δ2/+ animals were obtained at slightly lower frequencies than expected (Δ1/+ 25.66%; Δ2/+ 24.34%; Table 2). Loss of the enhancer region may be mildly detrimental in both homozygotes and heterozygotes. We did not assess why Δ1/Δ2 animals are lost during gestation or soon after birth, but Enh2-LacZ, Enh3-LacZ, and Enh2+3-LacZ reporters are all expressed during development in various patterns (Fig. S12). We did not test whether sites of LacZ staining overlap with *Wnt1* or *Wnt10b* expression. *LacZ* expression in the dorsal CNS reminiscent of the 3’ 5.5kb regulatory element that drives *Wnt1* expression during embryogenesis (Echelard et al., 1994; Danielian et al., 1997; Reddy et al., 2001; Veltmaat et al., 2004) was not observed. Further studies are needed to understand how low levels of lethality are caused by Δ1/Δ2 deletion, and the possible role of Enh2 and Enh3 in embryonic *Wnt1* and *Wnt10b* expression.

**Table 2:**
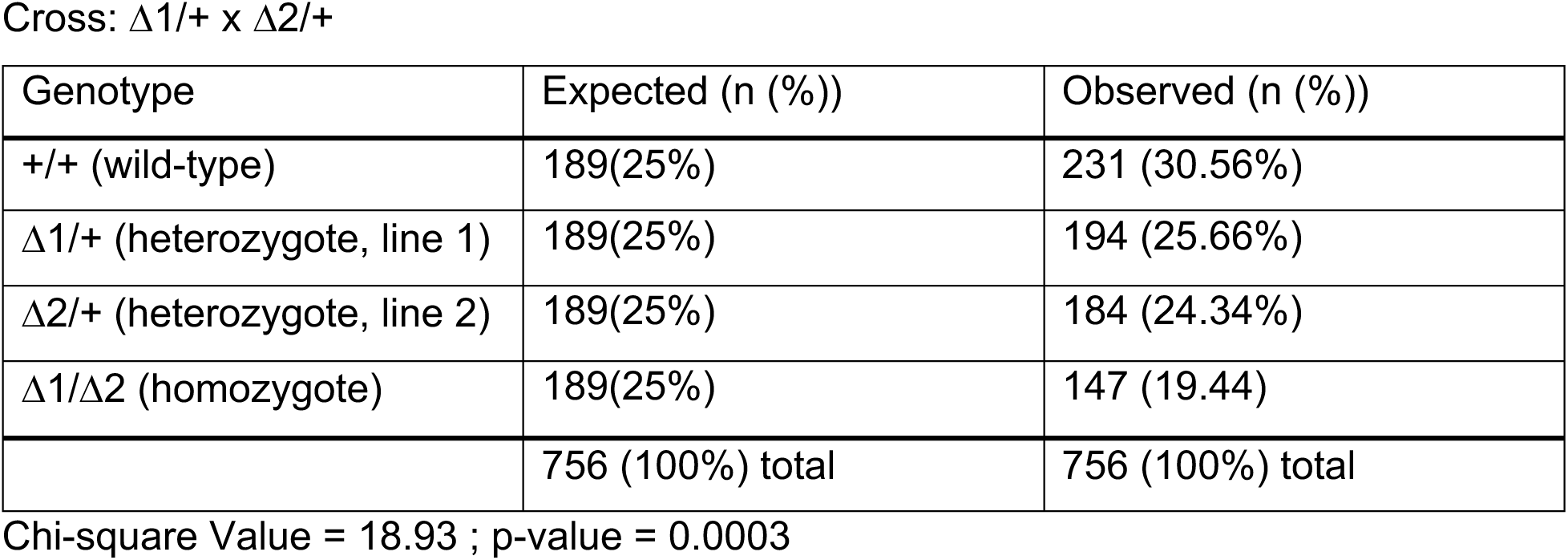
Mice carrying deletions of Enhancers 2 and 3 are generated at sub-Mendelian ratios.

Cross: $\Delta 1/+$ x $\Delta 2/+$
| Genotype | Expected (n (%)) | Observed (n (%)) |
| --- | --- | --- |
| +/+ (wild-type) | 189(25%) | 231 (30.56%) |
| $\Delta 1/+$ (heterozygote, line 1) | 189(25%) | 194 (25.66%) |
| $\Delta 2/+$ (heterozygote, line 2) | 189(25%) | 184 (24.34%) |
| $\Delta 1/\Delta 2$ (homozygote) | 189(25%) | 147 (19.44) |
|  | 756 (100%) total | 756 (100%) total |

Chi-square Value = 18.93 ; p-value = 0.0003

During the post-natal period, we observed no obvious lethality among pups, and of the Δ1/Δ2 mice that were born, body weights appeared normal (Figure 4C). We detected no obvious behavioral or morphological defects, suggesting that mice lacking Enhancers 2 and 3, once born, are mostly unaffected in the absence of injury.

We performed mRNA in situ hybridization for both *Wnt1* and *Wnt10b* transcripts to look for loss of *Wnt1* and *Wnt10b* expression following deletion of Enhancers 2 and 3 at 3 days (Figure 4D, F, G) and 5 days (Figure 4E-G) post-injury. Visual examination of stained tissue did not reveal a dramatic decrease in transcripts, but quantification of *Wnt1* and *Wnt10b* positive nuclei showed a gradual decline in gene expression over 3-5 days (Figure 4F, G). Comparison of *Wnt1* and *Wnt10b* levels in wild-type (wt) vs. Δ1/Δ2 (del) tissue showed a slight decrease at day 3, most obvious in violin plots when all quantified fields were plotted individually. By day 5, there was a ∼50% reduction in the number of *Wnt* positive nuclei. Given that Enhancer 2+3-LacZ promotes transcriptional activity in single cells during earlier timepoints in response to BaCl2-induced tissue damage, the gradual reduction in Wnt expression may reflect an inability of the Wnts to maintain sustained expression when Enhancers 2 and 3 are lost.

In contrast to *Wnt1* and *Wnt10b*, *Wnt5a* expression appeared unchanged in injured wt vs. Δ1/Δ2 tissues, as assessed by visual examination of transcript staining in sections (Figure 4D, E), and when the number of *Wnt5a* positive nuclei was quantified (Figure 4H). These data are consistent with the hypothesis that Enh2 and Enh3 specifically drive *Wnt1* and *Wnt10b* expression, and *Wnt5a* expression in the muscle is regulated independently of *Wnt1* and *Wnt10b*.

To ask if we could show by a different method that Enh2 and Enh3 control *Wnt1* and *Wnt10b* expression levels, we performed Pyrosequencing at 3 days-post injury. By crossing two different mouse strains (FVB and CAST/EIJ) carrying different SNPs that allowed us to distinguish *Wnt1* and *Wnt10b* gene expression driven by the wt vs. Δ1 or Δ2 alleles, we could measure how well Wnts were expressed when Enh2 and Enh3 were lost in the same muscle, under identical injury conditions. *Wnt1* and *Wnt10b* genes were similarly expressed by both alleles in FVB^wt^/CAST^wt^ mice, but expression was reduced from the Δ1 or /Δ2 alleles in FVB^λ−.1^/CAST^wt^ or FVB^λ−.2^/CAST^wt^ mice (Fig. S13). Together, these data show that Enhancers 2 and 3 are injury responsive regulatory sequences that indeed likely regulate *Wnt1* and *Wnt10b* in injured muscle; not only are the enhancers sufficient to drive injury-responsive reporter activity, but they are also necessary for *Wnt1* and *Wnt10b* expression in damaged tissues.

### Deletion of Enhancers 2 and 3 produces adipogenesis in injured muscle

When we asked whether Enh2+3 deletion might generate a phenotype, we noticed that Δ1/Δ2 TA muscles contained elevated fat at 14 days post-BaCl2 injection. Fat was detected by staining with an antibody to Perilipin 1, which marks adipocytes (Blanchette-Mackie, 1995) (Figure 5A-C, Fig. S14G-I). Uninjured muscles showed low Perilipin staining (Figure 5A, Fig. S14G) but both wt and Δ1/Δ2 muscles displayed a range of Perilipin signals at 14 days post-injury (Figure 5B,C, Fig. S14H,I). Highest levels of Perilipin staining observed in wt muscles was always reduced compared to that of Δ1/Δ2 muscles (Figure 5D,E, Fig. S14C,H,I). Representative muscles spanning the range of fatty infiltration observed for each genotype are shown in Fig. S14H, I).

**Figure 5:**
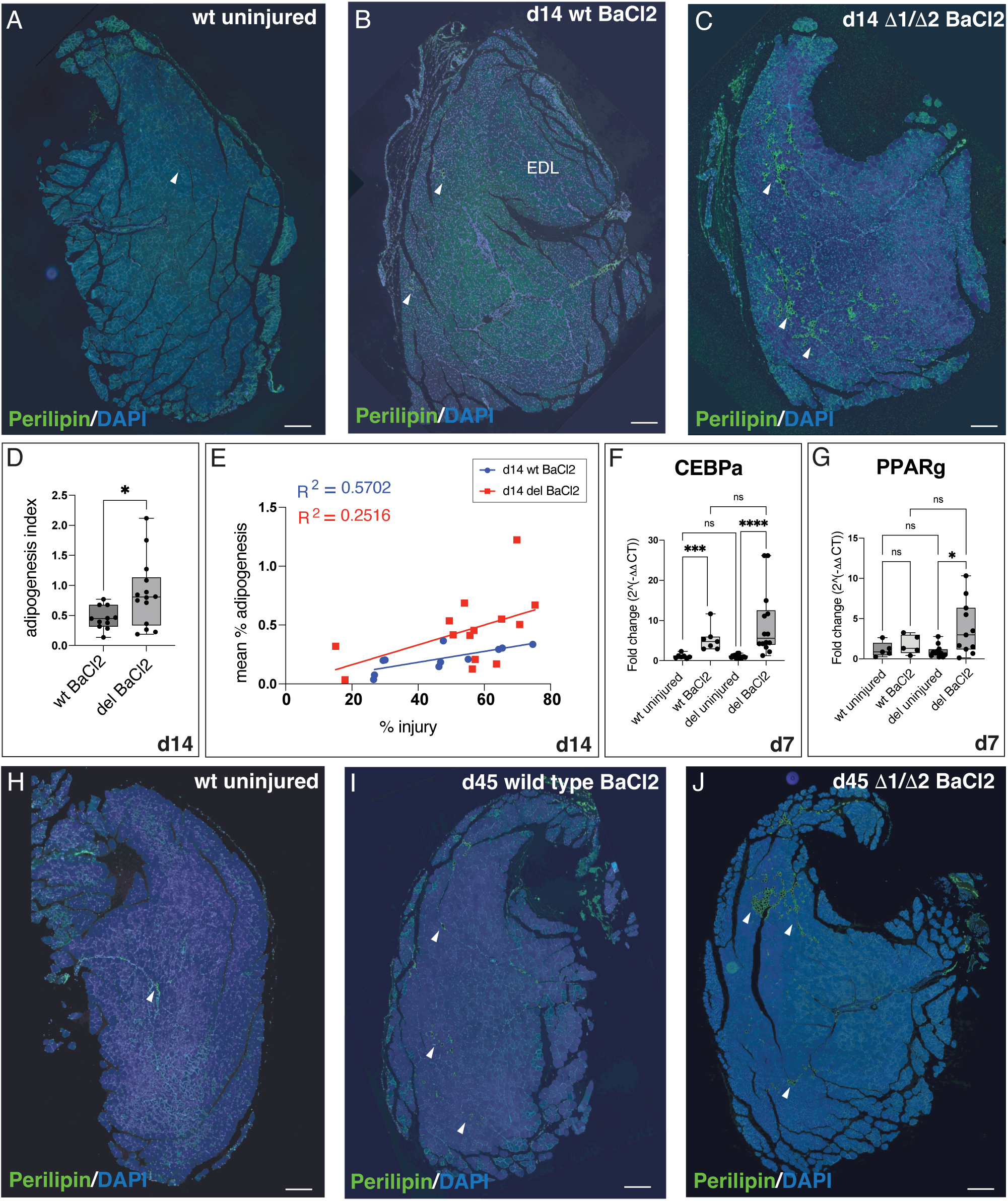
Mice lacking Enhancers 2 and 3 display elevated intramuscular adipogenesis. (A-C) Perilipin stained TA muscle sections. Arrowheads mark Perilipin positive areas. (A) Uninjured wt muscle shows low adipogenesis. (B) Wild-type muscles show small patches of adipocytes. (C) Δ1/Δ2 muscles show larger areas of adipocytes. (B) and (C) represent similarly injured muscles (∼75% injury). (D) Box and whisker plot showing the ‘adipogenesis index ‘ (mean % adipogenesis values normalized to % injury values). (n=11 wt, n=14 Δ1/ Δ2) (*p=0.0213, Welch’s t-test) (E) Scatter plot of mean % adipogenesis for wt and Δ1/Δ2 muscles plotted against levels of injury exhibited by BaCl2 injected muscles at 14 days. For wt muscles, a moderate positive correlation between % injury and % adipogenesis is seen (R^2^=0.5702, F(1,9) = 11.94, p=0.0072). For Δ1/Δ2 data, simple linear regression analysis shows a poorer fit (R^2^=0.2516, F(1,12)=4.04, p=0.0676). Slopes of the regression lines are not significantly different between the two genotypes (p=0.4596) but the difference in intercept is significant, showing that adipogenesis is independent of injury level, but dependent on genotype (F(1, 21)=4.727, p=0.0407). (n = 11 wt, n=14 Δ1/Δ2 muscles) (F) RT-qPCR of *CEBPa* expression following injury at 7 days. Uninjured: n=7 wt, n=14 Δ1/Δ2, BaCl2 injured: n=7 wt, n=14 Δ1/Δ2. (One-way ANOVA with Kruskal-Wallis test; wt uninjured vs. wt BaCl2: ***p=0.0003; Δ1/Δ2 uninjured vs. Δ1/Δ2 BaCl2: ****p<0.0001; ns, not significant) (G) RT-qPCR of *PPARg* expression post-injury at 7 days. Uninjured: n=5 wt, n=11 Δ1/Δ2, BaCl2 injured: n=5 wt, n=11 Δ1/Δ2. (Ordinary one-way ANOVA; Δ1/Δ2 uninjured vs. Δ1/Δ2 BaCl2, *p=0.0221; ns, not significant). (H-J) Perilipin stained muscle 45 days post-injury (arrowheads). (H) Representative example of Perilipin staining in uninjured wt muscle. (I) Wild-type and (J) Δ1/Δ2 muscle stained for Perilipin at 45 days post-injury. Images representative of average Perilipin staining levels at this timepoint are shown (calculated as 0.17% (wt) 0.67% (Δ1/Δ2); Overall average is 0.26% (wt), 0.74% (Δ1/Δ2)). EDL=Extensor Digitalis Longus. Graphs show mean ± s.d. (Scale Bar: 200 um)

We quantified adipogenesis in injured muscle of wt and Δ1/Δ2 mice at 14 days post-injury. We measured levels of adipogenesis over total section area and generated ‘% adipogenesis’ values over 30-50 sections per muscle, because fatty infiltration levels can fluctuate over the length of injured muscle (Fig. S14A). The extent of injury can also vary between BaCl2 injected muscles, so we also calculated a ‘% injury’ value for each sample by quantifying areas displaying centrally located nuclei (Folker and Baylies, 2013; Hardy et al., 2016; Meyer, 2018) and normalizing the values to the total section area. We further generated an ‘adipogenesis index’ by normalizing the % adipogenesis value to % injury value. The adipogenesis index value was significantly higher in Δ1/Δ2 muscles compared to wt muscles (Figure 5D).

When we plotted mean % adipogenesis values against % injury values, it was evident that although increased injury positively correlated with increased fat accumulation, Δ1/Δ2 muscles displayed much higher and more variable levels of fat (Figure 5E). Simple linear regression analysis of wt vs. Δ1/Δ2 samples did not show significantly different slopes between the two genotypes, but a significant difference in elevation (intercept) of the lines was detected. This supports the observation that loss of Enhancers 2 and 3 and the accompanying decrease in Wnt expression results in more fatty infiltration in Δ1/Δ2 muscle compared to wt. Additional visualization of the data by individual mouse or by mean % adipogenesis values support the observation that there is higher adipogenesis in Δ1/Δ2 muscles compared to wt (Fig. S14B-D).

We also quantified adipogenesis using H&E stained sections in which adipocytes appear as round ‘holes’ wherever fat is present (Biltz and Meyer, 2017). This method, in which adipocyte area in wt vs. Δ1/Δ2 muscles was measured and compared shows elevated fat accumulation in injured Δ1/Δ2 muscles compared to wt (Fig. S14E). We additionally measured expression levels of well-known adipogenic regulators, *CEPBa* and *PPARg*, at 7 days post-injury by RT-qPCR. Both genes were elevated in Δ1/Δ2 samples compared to wt (Figure 5F,G), although we were unable to observe statistical significance.

To ask if increased adipogenesis in Δ1/Δ2 muscles stems from reduction in Wnt signaling, we measured *Axin2* expression, a downstream target of the Wnt/b-catenin pathway (Jho et al., 2002; Lustig et al., 2002). We observed reduced *Axin2* expression at 5 days post-injury in Δ1/Δ2 muscles compared to wt, but the results were not statistically significant (Fig. S14F). *Axin2* expression is still abundant within wt and Δ1/Δ2 muscles (Ct values = ∼26-27), suggesting that other Wnts likely maintain signaling levels even when *Wnt1* and *Wnt10b* are lost.

Adipogenesis in the muscle is thought to be a transient process that resolves quickly after injury within a few days (Wagatsuma, 2007; Lukjanenko et al., 2013). We asked if elevated adipogenesis is observed if muscles recover longer. We injured the TA muscle of wt and Δ1/Δ2 mice with BaCl2 and waited for 45 days. We observed more adipocytes in wt muscles post-injury compared to uninjured muscles (Figure 5 H,I) indicating that adipogenesis does not necessarily completely resolve by 45 days. Moreover, Δ1/Δ2 muscles continued to display elevated fatty infiltration compared to wt (Figure 5 J,K, Fig. S15 A-D), although this effect was most pronounced in 2 out of the 8 mice analyzed (Fig. S15 C,D). These data show that even at 45 days, there is lingering fatty infiltration that can be detected at higher levels in Δ1/Δ2 muscles compared to wt.

Since muscle regeneration involves reciprocal cell-cell interactions between satellite cells, FAPs and other cell types (Joe et al., 2010; Uezumi et al., 2010; Lemos et al., 2015; Wosczyna et al., 2019), we asked if changes in FAP functions due to enhancer deletion might manifest as additional defects. We examined expression of *PDGFRa*, *Pax7*, and *CEBPb* using RT-qPCR. Both *PDGFRa* and *Pax7* increase post-injury as satellite cells and FAPs proliferate (Joe et al., 2010) but no differences were observed between wt and Δ1/Δ2 tissues (Fig. S16 A,B). *CEBPb* expression normally decreases following injury and upon satellite cell exit from quiescence (Lala-Tabbert et al., 2021); wt and Δ1/Δ2 muscles displayed similar *CEBPb* expression (Fig. S16C). We examined myofiber cross sectional area (CSA) across whole muscle sections (Fig. S16D,E) and also quantified CSAs only in injured myofibers, defined as those exhibiting centrally located nuclei (Fig.S16F), but no differences were observed. Finally, Collagen deposition within injured muscles at 14 days post-injury was also similar between wt and Δ1/Δ2 muscles (Fig. S16G,N).

Since CRISPR-Cas9-mediated deletion of Enhancers 2 and 3 is not a conditional allele and produces animals constitutively lacking the enhancer region from birth, we also asked if muscle defects were detectable prior to injury. We surveyed basal levels of *PDGFRa* (Fig. S16H), *Pax7* (Fig. S16I), myofiber CSAs (Fig. S16J, K), myosin heavy chain expression *(Myh 1, 2, 4*) (Fig. S16L) and Collagen deposition (Fig. S16M,N), but we detected no significant differences between wt and Δ1/Δ2 muscles.

Finally, we asked if loss of Enhancers 2 and 3 only affects muscle, or whether defects are found in other tissues. We saw no obvious phenotype when airways were injured using Naphthalene (data not shown). Because airway cells were restored and regeneration appeared normal, we did not examine the airway further; it is possible that we missed subtle defects. In the skin, Enhancer 2 and 3 reporters were expressed following punch biopsy (Fig. S17A), but unlike the muscle and lung, *Wnt1* and *Wnt10b* were not detected in areas surrounding the injury (Fig. S17B). Moreover, Enhancer 2+3-LacZ reporter mice failed to stain with X-gal (Fig. S17A). Enhancers 2 and 3 may each bind transcription factors induced by injury, but together do not drive gene expression. Skin punch biopsy experiments also produced no detectable wound closure defects (Fig. S17C,D). These results demonstrate that activation of individual enhancers may not always necessarily translate into functional target gene expression, underlining the importance of tissue context.

Together, these data show that loss of Enhancers 2 and 3 produces a regenerative defect specifically in the muscle, in which elevated fatty infiltration is observed. We do not know if levels of adipogenesis seen here adversely affect muscle function. However, the presence of fat in muscle is often observed in aged, disused or diseased muscle, or in experimental contexts where muscle regeneration is perturbed (Sciorati et al., 2015; Pagano et al., 2018; Wang et al., 2024). Our results suggest that the Enh2/3 regulatory region, a ∼1kb sequence that regulates *Wnt1* and *Wnt10b* in response to injury, is required to produce the correct balance of cell types as muscles recover from injury.

## Discussion

We have identified an enhancer region residing between *Wnt1* and *Wnt10b* genes in the mouse that responds to tissue injury and is required for muscle regeneration. Using an in vivo approach to study a single injury-responsive enhancer locus, we describe its spatial and temporal activities, examine its requirement for Wnt expression and proper cell fate choices during muscle regeneration, and extend our understanding of how Wnts activated in response to injury facilitate tissue regeneration.

Evidence that *Wnt1* and *Wnt10b* share a single injury-responsive regulatory region is supported by the observation that expression of both Wnts is greatly reduced when Enhancers 2 and 3 are deleted (Figure 4D-G, Fig. S13). Co-expression of *Wnt1* and *Wnt10b* in the same cells is detectable but infrequent however (Figure 1G), perhaps resulting from differences in mRNA stability or the duration and frequency of enhancer contacts with each Wnt promoter. Understanding how Enhancers 2 and 3 engage with promoters and other regulatory elements to drive *Wnt1* and *Wnt10b* expression post-injury in its full genomic context will require further studies.

To assess the importance of Enhancers 2 and 3, we made germline deletions of this regulatory region. Except for mild detrimental effects during embryogenesis (Table 2), uninjured animals lacking the enhancers are viable, fertile, and display no obvious defects (Figure 4C, Fig. S16). Muscles post-injury, however, show higher levels of adipocytes in Δ1/Δ2 tissues compared to wt (Figure 5, Fig. S14, S15) indicating impaired muscle regeneration, as fatty infiltration is often observed in diseased, disused, or aging muscle (Marcus et al., 2010; Hamrick et al., 2016; Zhu et al., 2024). Penetrance of the phenotype was variable, but this is not surprising given the deletion of regulatory, rather than coding sequence. Only a few examples of injury-responsive enhancers that produce phenotypes when deleted or mutated have been reported (Hewitt et al., 2017; Soukup et al., 2019; Suzuki et al., 2019; Harris et al., 2020; Wang et al., 2020; Zlatanova et al., 2023), and redundancy, a common feature of developmental enhancers (Osterwalder et al., 2018), may make it difficult to detect enhancer functions in adult tissues. The phenotype we describe here, to our knowledge, is the first to show a requirement for a regulatory region that regulates Wnt expression during regeneration in higher vertebrates.

Muscle regeneration involves complex cell-to-cell signaling interactions between FAPs, satellite cells, and immune cells (Joe et al., 2010; Uezumi et al., 2010; Lemos et al., 2015; Wosczyna et al., 2019). Following muscle injury, we observe *Wnt1* and *Wnt10b* in small cells scattered between myofibers that likely includes FAPs and other cell types (Figure. 1E,F). In the visceral fat, *Wnt10b* inhibits adipogenesis (Longo et al., 2004; Christodoulides et al., 2006; Cawthorn et al., 2012) and *Wnt10b*-/-mutants display fat in injured muscles (Vertino et al., 2005). A simple model is to propose that *Wnt1/Wnt10b* prevents FAPs from making adipocytes. When *Wnt1/Wnt10b* expression is reduced, as observed in Δ1/Δ2 tissues, adipocytes may be generated due to decreased suppressive signals. Other Wnts are induced following muscle injury, however, and several play important roles in regeneration (Van Winkle et al., 1995; Le Grand et al., 2009; Reggio et al., 2020). Whether their presence in Δ1/Δ2 muscles compensates for *Wnt1* and *Wnt10b* loss and can explain the relatively mild adipogenic phenotypes we observe is not known. Further work will be required to understand the specificity vs. redundancy of different Wnt ligands, the specific cell-cell interactions between different muscle cell sub-populations post-injury, and how *Wnt1* and *Wnt10b* expression is regulated in different temporal and spatial domains within muscle, both by Enhancers 2 and 3, and likely by additional regulatory elements.

We acknowledge several limitations of our study. First, we did not rescue the adipogenic phenotype to demonstrate that it stems from loss of *Wnt1* and *Wnt10b*. Designing a robust rescue system for two Wnts in vivo is challenging, and Wnt over-expression by tools such as AAV injections incur additional tissue damage. *Wnt10b* plays an inhibitory role in visceral fat by preventing pre-adipocyte differentiation into adipocytes (Christodoulides et al., 2009; Perkins et al., 2023). Expressing *Wnt1* in vitro has a similar inhibitory activity (Ross et al., 2000), although an endogenous role for *Wnt1* in adipogenesis has not been reported. In muscle, in vitro experiments failed to show an effect of Wnt10b on isolated mouse FAPs (Reggio et al., 2020), but *Wnt10b*-/-mutants display excess adipocytes following muscle injury in vivo (Vertino et al., 2005), suggesting that *Wnt10b* (and possibly *Wnt1*) are inhibitory for adipocyte differentiation in muscle.

We were also unable to perform genetic sensitization experiments to assess the role of other genes that might interact with *Wnt1* and *Wnt10b* to regulate adipogenesis. We introduced one copy of a *b-catenin* Δ allele (Brault et al., 2001) and one copy of a *Wntless* Δ allele (Carpenter et al., 2010) (both in mixed backgrounds) into Δ1/Δ2 mice (FVB background) to ask if reducing b-catenin or reducing all Wnts might worsen fatty infiltration in muscle. The results were uninterpretable, with mice displaying variable adipogenesis in both wt and Δ1/Δ2 animals. We discovered, perhaps not surprisingly, that genetic background profoundly affects enhancer phenotypes, particularly when the defects observed are subtle.

Finally, we asked if older Δ1/Δ2 mice might display more severe defects, since aging increases fatty infiltration in muscle (Zhu et al., 2024). We performed experiments twice on 16 to 24 month old wt and del animals, but the mice did not survive. BaCl2 can induce arrhythmias (Mattila et al., 1986; Jung et al., 2019), which perhaps old mice are less able to tolerate. When we reduced BaCl2 dosage on a third cohort, the animals survived but displayed poor injury. Evaluating the effects of Enhancer 2 and 3 deletions in aged mice will require further experiments.

Based on work done in invertebrates, it has been proposed that an ancient and conserved injury-responsive gene regulatory network might exist for Wnt gene regulation (Srivastava, 2021). *Drosophila* harbors a cluster of Wnts that reside on the same chromosome consisting of *Wnt4/Wingless/Wnt6/Wnt10*. A damage-responsive element between *wingless* and *Wnt6* utilizes AP-1 binding to drive Wnt expression (Harris et al., 2020), perhaps reminiscent of the close gene arrangement of the putative AP-1 binding sites and Wnts we see in mouse. Given Wnt induction in virtually all injury contexts that have been described to date, could there be common mechanisms that activate different Wnts post-injury that could span across invertebrates to vertebrates? A definitive answer to these possibilities will require further functional studies in other non-mammalian species.

## Materials and Methods

### Mouse husbandry and Mouse lines

All animal experiments and methods were approved by the Institutional Animal Care and Use Committee (IACUC) at Stanford University. Experiments were performed on FVB mice (Charles River) at 12-18 weeks of age. Both males and females were used in this study. Data are combined, unless differences were noted and specified in the text. For timed matings, plugs were checked in the morning before 9am and noon was considered E0.5 d.p.c. Wnt1-LacZ mice (Echelard et al., 1994) and Castaneus mice (CAST/Eij strain) were obtained from Jackson Laboratories (Wnt1-LacZ, stock# 002865; CAST/Eij, stock #000928). Transgenic mice were generated by the Stanford Transgenic Knockout and Tumor Model Center and Cyagen Biosciences (Santa Clara, CA, USA) through pronuclear injection of transgene constructs into FVB embryos (Hogan, 1994). CRISPR-cas9 mediated deletion of Enhancers 2 and 3 was carried out by the Stanford Transgenic Knockout and Tumor Model Center.

### Injury models

Naphthalene injuries were performed before 9 am via a single intraperitoneal injection of 0.2 um sterile-filtered Naphthalene (Sigma, #147141) dissolved in corn oil (Sigma, #C8267) at a concentration of 275mg/kg as previously described (Reynolds et al., 2000; Hong et al., 2001). For muscle injuries, mice were anaesthetized with isofluorane and given either Buprenorphine SR (1mg/kg, subcutaneously) or Ethiqa XR (3.25 mg/kg subcutaneously) prior to injection of BaCl2. 50 ul of sterile filtered 1.2% BaCl2 (Sigma #342920) solution was injected into the left tibialis anterior (TA) muscle. The muscle was poked to distribute the BaCl2 as described in (Wosczyna et al., 2019). The right TA muscle served as the uninjured control. For injuries of pancreatic islets (Furman, 2021), animals were initially fasted for 6 hours, and their weights and blood glucose levels were measured. Then, a 1-time injection of 175mg/kg Streptozotocin (STZ) (Sigma, #0130) was administered intraperitoneally. After injection, animals were given sucrose water (15g/L) for 48 hours, and blood glucose levels were monitored at 2, 5 and 7 days. High blood glucose readings by 5-7 days indicate successful injury to the pancreatic islets, and pancreatic tissues were isolated at 7 days. For liver injury using CCl4, sterile filtered CCl4 (Sigma, #289116) dissolved in corn oil (Sigma, #C8267) at a 1:4 ratio was injected at a dose of 1ml/kg. Sterile corn oil served as the vehicle control (Zhao et al., 2019). Punch Biopsy of the skin was performed similarly to the procedure described in (Lindsey, 2020). Briefly, mice anaesthetized with isofluorane were shaved and the skin was cleaned with 10% Povidone Iodine (PDI, #B40600). Topical Lidocaine/Prilocaine Cream (Alembic) was administered where the biopsy site would be located. Mice were then subcutaneously given Carpofen (5mg/kg) or Ethiqa XR (3.25 mg/kg). A 5mm biopsy punch (Acuderm) was used to excise a full-thickness skin sample.

### mRNA in situ hybridization

Tissues were fixed in 4% Neutral-buffered formalin for 24 hours at room temperature (RT) and then processed for paraffin embedding. Sections were cut at 5 um thickness, and mRNA in situ hybridization using either the RNAscope 1-Plex, or 2.5 HD Duplex RNA Detection Kits was performed according to the manufacturer’s instructions (ACDBio). Wnt Probes for in situ hybridization were Mm-Wnt1 (#401091, NM_021279.4, region 1204 - 2325); Mm-Wnt2 (#313601, NM_023653, region 857– 2086); Mm-Wnt2b (#405031, NM_009520.3, region 1307 - 2441); Mm-Wnt3 (#312241, NM_009521.2, region 134 - 1577); Mm-Wnt3a (#405041, NM-009522.2, region 667 - 1634); Mm-Wnt4 (#401101, NM_ 009523.2, region 2147–3150); Mm-Wnt5a (#316791, NM_009524.3, region 200–1431); Mm-Mm-Wnt5b (#405051, NM_001271757.1, region 319 - 1807); Mm-Wnt6 (#401111, NM_009526.3, region 780 - 2026); Mm-Wnt7a (#401121, NM_009527.3, region 1811 - 3013); Mm-Wnt7b (#401131, NM_009528.3, region 1597 - 2839); Mm-Wnt8a (#405061, NM_009290.2, region 180 – 1458); Mm-Wnt8b (#405071, NM_011720.3, region 2279 - 3217); Mm-Wnt9a (#405081, NM_139298.2, region 1546 - 2495); Mm-Wnt9b (#405091, NM_011719, region 727–1616); Mm-Wnt10a (#401061, NM_009518.2, region 479 - 1948); Mm-Wnt10b (# 401071, NM_011718.2region 989 - 2133); Mm-Wnt11 (#405021, NM_009519.2, region 818 - 1643); Mm-Wnt16 (#401081, NM_053116.4, region 453 - 1635). Additional probes were: Mm-Pax7 (#314181-C2, NM_011039.2, region 602 – 1534); Mm-PDGFRa (#480661-C2, NM_011058.2, region 223-1161), Mm-Adgre (#460651-C2, NM_ 010130.4, region 85-1026), Mm-FosL1 (#421981, NM_010235.2, region 2-1386), Mm-FosL2-C2 (#421991-C2, NM_008037.4, region 1612 – 2669), Mm-Fosb (#539721, NM_008036.2, region 370 – 1302); Mm-Fos (#316921-C2, NM_010234.2, region 407 - 1427), MmWnt10b-C2 (# 401071-C2, NM_011718.2, region 989 - 2133), Mm-Wnt1-C2 (#401091-C2, NM_021279.4, region 1204 - 2325). Every in situ experiment was run with a negative and positive control: DapB (negative control probe, EF _191515, region 414–862), and Polr2a (positive control probe, NM_009089.2, region 2802–3678) or 2.5 Duplex Positive Control Probe (#321651), 2-Plex Negative Control Probe (#320751). Images were captured on a Zeiss Axio Imager.Z2 microscope. Brightness was adjusted globally on images using Photoshop to improve visualization in figures submitted for publication. For quantification of number of nuclei expressing different Wnts, Gill’s Hematoxylin-stained nuclei (StatLab HXGHE1LT) and nuclei stained with the different in situ probes were counted using the multi-point tool in ImageJ/Fiji. Values for % Wnt positive nuclei were calculated as the total number of Wnt-positive nuclei divided by the total number of nuclei counted in each field X 100. All data were analyzed using GraphPad (Prism) software.

### Construction of Transgenic mice

The plasmid used for construction of Transgenic mice was developed by Kothary et al., (1989) and was a gift of Dr. David Kingsley (Stanford University, Stanford, CA). For the generation of reporter mice, BAC RP2394M12 DNA (CHORI) and BAC RP23236-E21(CHORI, for the 6kb construct) were used as templates. Enhancer segments were amplified using PCR primers designed with Not1 restriction sites. Coordinates for the different enhancers (mm9 assembly) are: 6kb: 98608459-98614672, Enhancer 1: 98609340-98609567, Enhancer 2: 98610564-98610827, Enhancer 3: 98610878-98611376.

Primers for the enhancer sequences were:

6kbF Not1: 5’ ATA AGA ATG CGG CCG CAG ATG TAG AAC TCT CAG CTC CTC CTG AAC C 3’, 6kbR Not1: 5’ ATA AGA ATG CGG CCG CGT CTG TCT GTC TGT CTG TCT GTC TGT CTG T 3’; Enhancer1F Not1: 5’ ATA AGA ATG CGG CCG CCG GCT CTG GAT CTA TGT GAC TT 3’, Enhancer1R Not1: 5’ ATA AGA ATG CGG CCG CAG GCA AAG GCA CAA ACT ACG 3’; Enhancer2F Not1: 5’ ATA AGA ATG CGG CCG CCC TCT CTG AAG TCT TGC TCT TTG 3’, Enhancer2R Not1: 5’ ATA AGA ATG CGG CCG CTC CCC CTC TCA TCT AGG TCT C 3’; Enhancer3F Not1: 5’ATA AGA ATG CGG CCG CTA GTA GAC AAG GGG GTA ACA TTC TG 3’, Enhancer3R Not1: 5’ ATA AGA ATG CGG CCG CCA TCT TCC TTT TGA TGA GAA AAG TG 3’, Enhancer 2/3 was cloned using the Enhancer2F Not1 and Enhancer 3F Not1 primers. The Enhancer 2 element is 264bp. The Enhancer 3 reporter carries a 524bp element.

PCR fragments were cloned into the Not1 sites of hsp68-LacZ and sequenced. Plasmids for injection were isolated using the Qiagen Maxiprep kit (Qiagen, 12662) and submitted for linearization and injection by the Stanford Transgenic Knockout and Tumor Model Center and by Cyagen Biosciences. Enzymes for linearization were designed to eliminate as much plasmid sequence as possible and were as follows: 6kb and Enhancer 2: ApaI/NaeI, Enhancer1: DraI/XhoI, Enhancer3, Enhancer 2/3 and hsp68-LacZ: NaeI/XhoI. For Enhancer3-eGFP, the eGFP sequence from pEGFP-N1 (Clontech) and the Enhancer3-hsp68 promoter sequence were cloned into pBluescript. An EcoRI/EcoRI PCR fragment containing the enhancer and basal promoter obtained from Enhancer3-LacZ was generated and fused to an EcoRI/XhoI PCR fragment carrying eGFP and an SV40 polyA tail. The plasmid sequenced and then linearized with PvuI/XhoI prior to pronuclear injection.

Transgenic mice were identified by genotyping with primers to LacZ: Galp1: 5’-TTTACAACGTCGTGACTG-3’, Galp2:5’-TGATTTGTGTAGTCGGTT-3’ Galp1 and Galp 2 (Dr. W.T. O’Brien). Transgenic mice carrying eGFP were genotyped using primers GFP03: 5’-ACGGCAAGCTGACCCTGAAGT-3’ and GFP04: 5’-GCTTCTCGTTGGGGTCTTTGC-3’. To generate adult transgenic lines, LacZ or GFP positive pups were crossed and tested through at least the G2 generation to eliminate mosaicism, separate instances of multiple insertions on different chromosomes, and to confirm mendelian segregation of single transgenic loci.

Because we were searching for injury-responsive enhancers, it was important to distinguish between the inability to be activated by injury vs. lack of expression due to integration into a silenced locus. Because the mouse brain is thought to express over 80% of genes within the genome (Lein et al., 2007; Thompson et al., 2014), we reasoned that the brain would be one of the most transcriptionally active organs within the body, and that if our transgene failed to express in the brain, it could indicate that it was silenced. We therefore defined silencing of the transgene as those lines that failed to express in uninjured adult brain tissue and those that were not induced following injury. These animals were removed from the study and not analyzed further.

### X-Gal staining

For whole-mount staining, tissues were harvested and fixed at room temperature in 4% paraformaldehyde (PFA) for 10 minutes. Lungs were gently inflated with 4% PFA using a 5ml syringe attached to a 25Ga needle prior to rocking in PFA. A short fixation was critical to avoid losing activity of the b-galactosidase enzyme due to over-fixation. Tissues were washed in PBS containing 2 mM MgCl2, 0.01% deoxycholate and 0.02% Nonidet P-40, and then placed in X-gal staining buffer (1 mg/mL X-Gal (5-bromo-4-chloro-3-indolyl-b-D-galactopyranoside (Sigma-Aldrich), 4 mm K3 Fe(CN)6, 4 mm K4 Fe(CN)6 · 3 H2O, 2 mM MgCl, 0.01% deoxycholate and 0.02% Nonidet P-40 in PBS). After staining overnight, tissues were post-fixed in 4% PFA for 24 hours and then washed in PBS for examination as whole mount specimens on an M80 Leica dissecting scope, or for further processing into paraffin blocks which were cut at 5 um thickness. For X-gal staining on tissue sections, isolated tissues were fixed in 1% PFA for 1 hour on ice and then washed three times in PBS for 5 minutes at room temperature (RT) before being placed in OCT overnight with rocking at 4°C. Specimens were then embedded in OCT (Tissue-Tek) and frozen for cryosectioning. Tissues were sectioned at 8 um, dried briefly, fixed in 4% PFA at room temperature for 5 minutes, and then placed in X-gal staining solution. To compare relative activities of the Enhancer 2 and 3 reporters, slides were stained for two hours at room temperature. To obtain stained specimens that reach saturation, slides were incubated in X-gal staining buffer for 24 hours at room temperature. After staining, slides were post-fixed overnight at 4°C in 4% PFA. After sections were stained with X-gal, slides were counterstained with Nuclear Fast Red (Vector Labs) or EosinY (Sigma, #HT110116), dehydrated through Ethanol and Histoclear (National Diagnostics, #HS-200), and mounted with Cytoseal-60 (Epredia, #HT8310-4). Slides were imaged on a Zeiss Axio Imager.Z2 microscope. In compiling figures, global adjustments to brightness and contrast, color balance, and levels settings were made to provide uniformity across images.

### Identification of AP-1 binding sites and site-directed Mutagenesis

GREAT (McLean et al., 2010)was initially used to generate a list of potential candidates for putative transcription factor binding sites in Enhancers 2 and 3, in which a single AP-1 binding site was identified. A second AP-1 site was identified using TFSEARCH (http://diyhpl.us/∼bryan/irc/protocol-online/protocol-cache/TFSEARCH.html)

To generate mutant AP-1 sites, we identified two putative AP-1 binding sequences within Enhancer 3 with the sequence ‘TGAGTCA’ (mm9, chr15:98610877-98611400 (reverse complement)) and designed non-complementary base transversions to create a ‘GTCTGAC’ sequence. We noticed that the last four nucleotides of this altered sequence produced a new TGAC sequence (reminiscent of the AP-1 consensus sequence **TGA**G/**C**TCA) followed by a ‘TCT,’ in one of the two predicted AP-1 binding regions. Although this new sequence was unlikely to function as another AP-1 binding site, we further modified ‘GTCTGAC’ to ‘GTCAGTC’ to ensure that we were not creating a new site with AP-1 binding affinity. As of this writing, the sequence GTCAGTC resembles no known consensus binding sites.

Primers carrying the desired sequence GTCAGTC were generated. Primers were: NPP1F 5’-GGGTAACATTCTGTCTTGGT**<u>GTCAGTC</u>**GAAGACTCCTTGG-3’, NPP1R 5’-CCAAGGAGTCTTC**<u>GACTGAC</u>**ACCAAGACAGAATGTTACCC −3’, NPP2F: 5’ GCTTAGCAACAGA**<u>GTCAGTC</u>**CCCAAGACCC-3’, NPP2R: 5’ GGGTCTTGGG**<u>GACTGAC</u>**TCTGTTGCTAAGC −3’ The two sites were mutated sequentially using the Quick-Change Site Directed Mutagenesis Kit (Agilent #200523) following the manufacturer’s instructions. Wild-type Enhancer 3-LacZ was used as a template. Introduction of the first mutant site was verified by sequencing before introducing the second site, and the second mutant site was also confirmed by sequencing. Before injecting into mice, to verify that a functional promoter and LacZ gene were still present in the construct, wild-type Enhancer 3-LacZ and AP-1 mutant Enhancer 3-LacZ constructs were transfected into 293 cells along with a pCMV-GFP as a transfection control. Cells expressing the wild-type and AP-1 mutant constructs both showed b-gal activity in the transfected cells. Constructs were then linearized using Xho1 and Nae1 before pronuclear injection of the transgene (Cyagen Biosciences (Santa Clara, CA, USA)).

### Generation of mice carrying a deletion of regulatory sequences for Enhancers 2 and 3

To generate mice harboring a deletion of the Enh2/3 region, candidate guide RNAs were identified using CRISPRscan (Moreno-Mateos et al., 2015) (https://www.crisprscan.org/). Several candidate sgRNAs were screened for their ability to cleave a target template in vitro using a Guide-itT^M^ sgRNA In Vitro Transcription and Screening Systems Kit (Clontech), following the manufacturer’s instructions. A 2kb target template was generated by PCR using FVB genomic DNA amplified with the following primers: 5’-GAATGGTTGTGAGCCACCTT-3’ and 5’-GAGCTCCTTCCCATTTAGGG-3’. SgRNAs were synthesized using the HiScribe^TM^ Quick T7 High Yield RNA Synthesis Kit (NEB #E2050). sgRNAs with an in vitro cutting efficiency of over 50% were considered acceptable candidates. Two sgRNAs were selected: gG18NGG-53: 5’-TGGGCAAGCCTTTGAAGCCC-3’ and gG18NGG-63: 5’-AGGAAGATTATAGCGCCCAG. sgRNAs (10ng/ul) and Cas9 mRNA (30ng/ul) were introduced into FVB mouse 1-cell embryos by pronuclear micro-injection and transferred into pseudo pregnant CD-1 females by the Stanford Transgenic Knockout and Tumor Model Center. Genotyping primers for CRISPR-Cas9 mice carrying deletions of Enhancers 2 and 3 are Crispr Enh23 #2F:5’-GCTGCAGTTCCATTCACACGTTG-3’ and Crispr Enh23 #2R:5’-TGATCTCCCTCCCTCAACTTCCT-3’

### qRT-PCR

Tissue homogenization was performed in TRIzol reagent (Invitrogen) using a glass bead homogenizer (BeadBug, Benchmark Scientific, D1030, speed at 4000x, 3 cycles of 1 minute each). Linear Acrylamide (ThermoFisher Scientific, AM9520) was added and then RNA was purified using the RNeasy Mini Isolation Kit (Qiagen). Genomic DNA contamination was digested using RNAse-Free DNAse (Qiagen, 75254). cDNA was synthesized using a High-Capacity cDNA Reverse Transcription Kit (Life Technologies) according to the manufacturer’s protocol. cDNA was quantified on a NanoDrop 2000 Spectrophotometer (Thermo Scientific). qRT-PCR reactions which were performed with a TaqMan Gene Expression Master Mix (Applied Biosystems) using a StepOnePlus Real-Time PCR Instrument (Applied Biosystems). The relative expression of target genes was calculated using the DDCT method and fold changes were calculated relative to SRP14 as a normalization control as it was most stable between injured and uninjured muscles (Welc et al., 2020). Other genes tested were GAPDH, Actb, Hagh, Rps2; Hagh exhibited similar stability as SRP14, but in our hands, the other reference genes were not stable between uninjured and injured conditions. Probes utilized for assays were: Srp14 (Mm00726104_s1), Pax7(Mm01354484_m1), PDGFRa(Mm00440701_m1), Cebpb (Mm00843434_s1), Wnt1(Mm01300555_g1), Wnt10b (Mm00442104_m1), Wnt5a (Mm00437347_m1), CEBPa (Mm00514283_s1), PPARg (Mm00440940_m1), Axin2 (Mm_00443610). For ease of visualization, the graphs that display the 2^(-DDCT values) show statistical significance values that were calculated on the DDCT values. Data were analyzed using Microsoft Excel and GraphPad (Prism) software.

### Quantification of myofiber size

To measure myofiber cross sectional area, we utilized Alexa-647 conjugated Wheat Germ Agglutinin (WGA) (Molecular Probes #W32466) to stain myofibers because it allowed staining on paraffin sections (Aishwarya et al., 2022). 5 um muscle sections were deparaffinized and then rehydrated into PBS-0.1% Tween-20 (PBST) and incubated in WGA overnight (Molecular probes, Thermo Fisher Scientific, W32466, 1:500 dilution). The next day, slides were washed in PBST and mounted in ProlongGold with DAPI (Invitrogen, P36931). Slides were imaged on a Zeiss Axio Imager.Z2 with an Apotome and images were saved as .czi files and fed into Cellpose (Stringer et al., 2021) to segment the myofibers in WGA-stained sections. Cellpose has been used successfully to quantify CSAs of muscle tissue (Waisman et al., 2021). Sections were analyzed using the Labels to ROIs plugin in Fiji to generate area measurements. GraphPad (Prism) was used to generate histograms and violin plots of the cross-sectional areas. Histograms are displayed as mean ± s.e.m.

### Picrosirius red staining and quantification of collagen deposition

To measure collagen deposition in tissue sections, 5 um paraffin sections were deparaffinized and rehydrated into PBS. Then they were stained using a Picrosirius Red Stain Kit (Connective Tissue Stain) (Abcam, ab150681) following the manufacturer’s instructions. Slides were then imaged on a Zeiss Axio Imager.Z2. To quantify the staining, images were processed in NIH Image (Fiji). The total number of pixels was calculated per field. Then, using the Threshold function the darkly stained collagen areas were selected. Images were changed to binary and the number of pixels marking the collagen-stained region was quantified. The number of collagen pixels was divided by total pixels in each field to give a measurement for picrosirius red staining. Data was entered into GraphPad (Prism) for analysis. For each mouse examined, at least 3-5 sections per mouse were generated and multiple fields were analyzed to generate an average for each animal. Graphs were plotted with the mean ± s. d. Statistical analysis was performed using GraphPad (Prism).

### Pyrosequencing

To measure allele specific expression of Wnt1 and Wnt10b using Pyrosequencing (Wittkopp, 2011), SNPs between CAST/EIJ mice and FVB/NJ mice were identified using the MGI SNP query tool (https://www.informatics.jax.org/snp, GRCm38). Primers were selected to amplify a region containing at least one SNP with products less than 300bp. All primers were generated by (Integrated DNA Technologies, Coralville, IA).

Forward and Reverse primers flanking the SNP for Wnt10b:

SNP_A-F: 5’-biotin-GAAAGGGCCTCCAAGAGTTAT-3’

SNP_A-R: 5’-TGTGGAGTCAATAAGACCCGTATA-3’

Sequencing primer for SNP_A: 5’-GAAAGGGTCTCTCCAA-3’

PCRs were run at 56°C for 45 cycles, giving a 266bp product.

Forward and Reverse primers flanking the SNP for Wnt1:

SNP_2-3-F: 5’-TTGCGCTGTGACCTCTTTGG-3’

SNP_2-3-R: 5’-biotin-AGCTTTCCGTGCCCTTTCAAC-3’

Sequencing primer for SNP_2-3: 5’-ACCTGTAGCTGAAGAGTT-3’

PCRs were run at 58°C for 45 cycles, giving a 154bp product.

Note that these primers did not span intron-exon boundaries. Primer sets were initially tested on purified FVB/NJ and CAST/EIJ genomic DNA that was extracted using Trizol (Invitrogen), precipitated, and quantified on a NanoDrop 2000 Spectrophotometer (Thermo Scientific). For Pyrosequencing experiments, TA muscles belonging to FVB^wt^, CAST/EIJ^wt^, FVB^λ−.1^ /CAST/EIJ^wt^, and FVB^λ−.2^ /CAST/EIJ^wt^ were injected with BaCl2, and tissues were collected at 3 days post-injury.

Tissue homogenization was performed in TRIzol reagent (Invitrogen) using a glass bead homogenizer (BeadBug, Benchmark Scientific, D1030, speed at 4000x, 3 cycles of 1 minute each). Linear Acrylamide (ThermoFisher Scientific, AM9520) was added and then RNA was purified using the RNeasy Mini Isolation Kit (Qiagen). Genomic DNA contamination was digested using an RNAse-Free DNAse (Qiagen, 75254). cDNA was synthesized using a High-Capacity cDNA Reverse Transcription Kit (Life Technologies) according to the manufacturer’s instructions. cDNA was quantified on a NanoDrop 2000 Spectrophotometer (Thermo Scientific). cDNA was used as input for PCR reactions. Products were run on an agarose gel to check for amplification of correctly sized products. Pyrosequencing runs were performed by the Stanford Protein and Nucleic Acid Facility (PAN) on a Pyromark Q24, with results reported as percentages of each nucleotide that was observed at the SNP site.

Statistics were performed on percent values. Allelic percentages were baseline-corrected using the observed minor-allele percentages in the 0% and 100% controls which defined the lower and upper bounds. Sample values were then linearly re-scaled before performing statistical analysis. Corrected proportions were then acrsine square-root transformed for ANOVA (Hsiao et al., 2012). Plots display the corrected % FVB allele percentages for interpretability, but significance values are from the statistical analysis using ANOVA on arcsine transformed values. Data were analyzed using Microsoft Excel and GraphPad (Prism) software.

### Perilipin Staining

Wt and Δ1/Δ2 mice were injured with BaCl2, and at 14 days post injury, TA muscles were isolated and fixed in NBF overnight at 4°C. Muscles were then washed in PBS and processed for Paraffin embedding and sectioning. To sample a large region of the muscle, serial sections with 10 sections per slide consisting of sections spaced 50 um apart for a total of ∼2.5 mm was generated on 5 slides. Sections were deparaffinized and processed into PBS by standard methods. Slides were placed in 200 mls Tris Antigen Unmasking Solution (Vector Laboratories, H-3301) in an Instant Pot Duo Plus Mini (3qt) and pressure cooked for 20 minutes. Pressure was released manually, and slides were left to cool for 30-45 minutes on the benchtop. Slides were removed and a hydrophobic barrier was drawn around each section. Slides were blocked for 1 hour at room temperature (RT) in 10% NDS PBS-0.1% Tween (PBS-T) before incubation overnight in Perilipin 1 antibody (Sigma, P-1998, 1:200 dilution). The next day, slides were washed in PBS-T and then incubated in secondary antibody (Donkey anti-Rabbit Cy5, Jackson Immuno, #711-175-152, 1:1000 dilution). After washing in PBS-T, slides were mounted in Prolong Gold Antifade Reagent with DAPI (Invitrogen, P36931) and imaged on a BZ-X800 microscope (Keyence, Osaka, Japan) and montages were generated to make images of entire cross sections through the muscle. Images were re-oriented in Photoshop and placed on dark backgrounds to generate the figures, and brightness levels were globally adjusted to make staining easier to visualize, but no changes were made to the sections themselves, unless they were cropped by the software during image acquisition.

### Quantification of Adipogenesis by Perilipin Staining

To quantify the extent of adipogenesis in wt vs. del muscle, we first asked how uniformly adipocytes were distributed throughout the muscle at 14 days post-injury. We reasoned that the focal nature of the BaCl2 injury model might affect adipocyte distribution in the muscle post-regeneration. We cut 40-50 serial sections per muscle with each 5 um section spaced 50 ums apart, to survey a total of 2250-2750 um along the length of the muscle. We identified adipocytes using Perilipin antibody, evaluated section area using DAPI staining, and generated a % adipogenesis value for each section (see paragraph below). Both wild-type and Δ1/Δ2 muscle sections show that the presence of adipocytes can fluctuate over the length of the muscle, suggesting that surveying a larger number of sections distributed along the muscle would allow for a more accurate assessment of the overall extent of adipogenesis within the muscle tissue. Sections that displayed significant folds or tears were eliminated from the data set but overall, 30-50 sections were captured and quantified per muscle.

For quantification of Perilipin staining and generation of % adipogenesis value, montaged images of entire sections acquired on the BZ-X800 microscope (Keyence, Osaka, Japan) were imported into Fiji. If images contained extra debris, the EDL muscle, or portions of the epimysium, they were manually erased with the paintbrush tool to avoid quantifying non-TA muscle tissue. No portions of the TA muscle were eliminated. We were unable to develop a good automated sequence for image processing due to differences in DAPI and perilipin staining intensity between slides and specimens, and other features such as folds or wrinkles in some sections that made it difficult to uniformly apply the same parameters across all images. Therefore, processing was done manually but the following steps were applied: The Split Channels function was used to generate separate images for DAPI and Perilipin staining. To quantify the total section area in the DAPI image, the slider of the Threshold function was used to fill the entire section area. The image was then made binary, and the number of pixels in the total section was quantified using the Histogram function. For Perilipin staining, the original image was opened, and the brightness was adjusted using the Auto function to clearly show all perilipin stained adipocytes. The perilipin-only image was then placed next to the original image, and the Threshold function was used to highlight the perilipin adipocytes to make them match the original image. Care was taken not to erase or diminish the Perilipin signals in the single-channel image or to make the Perilipin contours of the adipocytes thicker or brighter than the original image. Speckled background signals in the Perilipin channel that were not part of the adipocyte staining pattern were then eliminated using the Despeckle function. Any folds or speckles that could not be eliminated that added non-adipocyte signals in the Perilipin channel were erased by hand to avoid including them in the quantifications. Binary images were generated, and the number of Adipocyte pixels were counted using the Histogram function. Values for the number of pixels calculated in the Perilipin channel was divided by the number of pixels calculated in the DAPI channel x 100, and this generated the % adipogenesis value. All data were analyzed using GraphPad (Prism) software.

### Quantification of adipogenesis by H&E staining

Quantification of adipogenesis by H&E staining was performed similarly to Perilipin measurements. Automated segmentation was difficult due to the inability of software to distinguish between myofiber and adipocyte shapes in the tissue sections, so analysis was performed manually. H&E stained sections were opened in Photoshop as RGB images, and contrast was enhanced to sharpen boundaries between the unstained adipocyte ‘holes’ and the surrounding stained myofibers. Adipocytes were selected and filled with black using the paint bucket tool. Brightness was decreased to intensify the black adipocyte shapes compared to the background and then identified with the magic wand tool and pasted into a separate image that was imported into FIJI. Data were converted into binary image and the number of pixels was quantified using the histogram tool. Overall section area was quantified by importing the same RGB file into FIJI, splitting it into separate channels. Using the green channel file which displayed the most contrast, the Threshold tool was used identify the contour of the section. Similar threshold values were applied across all files although small adjustments were made due to slight differences between images. Images were converted into binary and then the histogram tool was used to measure pixel values. % adipogenesis was calculated as # of adipocyte pixels/total # pixels in the section *100. All data were analyzed using GraphPad (Prism) software.

### Quantification of extent of injury

To quantify the extent of injury, 3 well-spaced sections were quantified for each muscle. In general, the values across all 3 sections were similar. If values diverged significantly, more sections were counted (up to 5). Total section area was quantified based on DAPI staining. Perilipin stained montaged images of whole muscle sections were imported into Photoshop and channels were split into single colors. The DAPI image was then imported into FIJI. The Thresholding function was used to highlight the entire section area, a binary image was generated and using the Histogram function, the total number of pixels comprising the section area was quantified. In Photoshop, injured area was then marked in the same DAPI-stained TIFF file by using the pencil tool to draw outlines around all myofibers containing centrally located nuclei. Often injured myofibers were found in large contiguous patches which could be outlined. These injured areas were then filled with the bucket tool to mask all portions of the section that were injured. This file was imported into FIJI, converted into a binary image and the pixels representing the injure regions were quantified using the Histogram function. Values for the number of injured area pixels was divided by the number of total pixels calculated for the section area, and this generated the % injury value. Additionally, an “Adipogenesis Index” was calculated to normalize % adipogenesis to the different % injury values which was reported as (% adipogenesis/% injury x 100). All data were analyzed using GraphPad (Prism) software.

### Assessment of dorsal punch biopsy wound closure over time

To follow skin wound healing over time, mice were anaesthetized with isofluorane and placed briefly in a cardboard chamber with two holes cut out on the top, to allow for illumination with a fiber-optic light source and placement of an iPhone SE for daily photography of wound closure. Animals were placed next to a ruler, dorsal side up. Images were collected every 24 hours. Quantification of wound size over time was measured by importing images into Fiji. Images were split into different channels using the Color function, and then the Threshold function was applied to the green channel to demarcate the wound area. The Make Binary function was applied to the image and the number of pixels was counted to quantify the size of the wound. Closure of the wound was compared to the initial size to generate a value for % of starting area over time. Animals were followed for 9 days. All data were analyzed using GraphPad (Prism) software.

## Acknowledgements

The authors thank the Stanford Core facilities, particularly the Stanford Pan Facility for Pyrosequencing, and the Stanford Transgenic Knockout and Tumor Model Center. We thank Dr. Hong Zeng (Stanford Transgenic Knockout and Tumor Model Center) for discussions on experimental design during generation of transgenic and CRISPR-Cas9 animals. We thank Pauline Chu, BS, HT, Dept. Comparative Medicine, Stanford University for tissue processing and embedding during early stages of this work. Additional contributors to early phases of this project include Dr. Barbara Brott, Johanna Kirby, Kia Fathi and Dr. Si Hui Tan. We thank Drs. Ross Metzger, Hernan Espinoza, Catherine Gunther, Ian Heller for many stimulating scientific discussions. We thank Drs. Mark Krasnow and David Kingsley for support and encouragement over many years. We also thank Drs. Aaron Wenger and Jim Notwell from Dr. Gil Bejerano’s laboratory for exploring possible transcription factor binding sites within our enhancers. Thank you to Drs. Thomas Rando and Mike Woczyna for training us on BaCl2 muscle injury, and Dr. Joseph Wu for teaching us the myocardial infarction model. Thank you to Dr. Zhibo Zhang for critical comments on this manuscript. This work is dedicated to the memory of Johanna Kirby and also Y. L. and D. H. M. L.

## Resource availability

Enhancer 2-LacZ, Enhancer 3-LacZ, Enhancer 3-eGFP, Enhancer 2+3-LacZ and Δ1/Δ2 animals will be made available at JAX.

