## Supplementary Figures, Legends, Tables for "A regulatory region that controls Wnt gene expression following tissue injury is required for proper muscle regeneration"

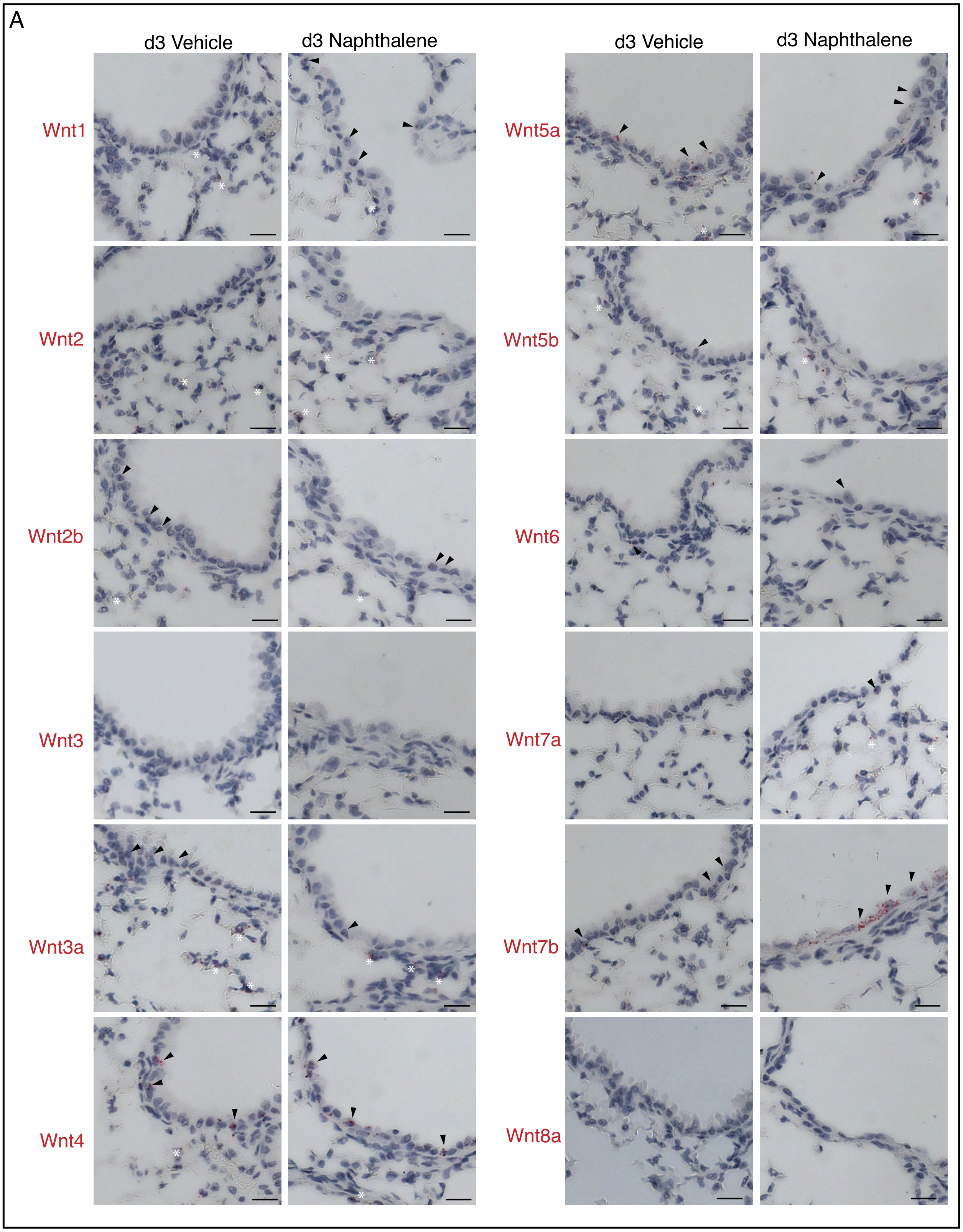
A
d3 Vehicle
d3 Naphthalene
d3 Vehicle
d3 Naphthalene
Wnt1
Wnt2
Wnt2b
Wnt3
Wnt3a
Wnt4
Wnt5a
Wnt5b
Wnt6
Wnt7a
Wnt7b
Wnt8a

Fig. S1

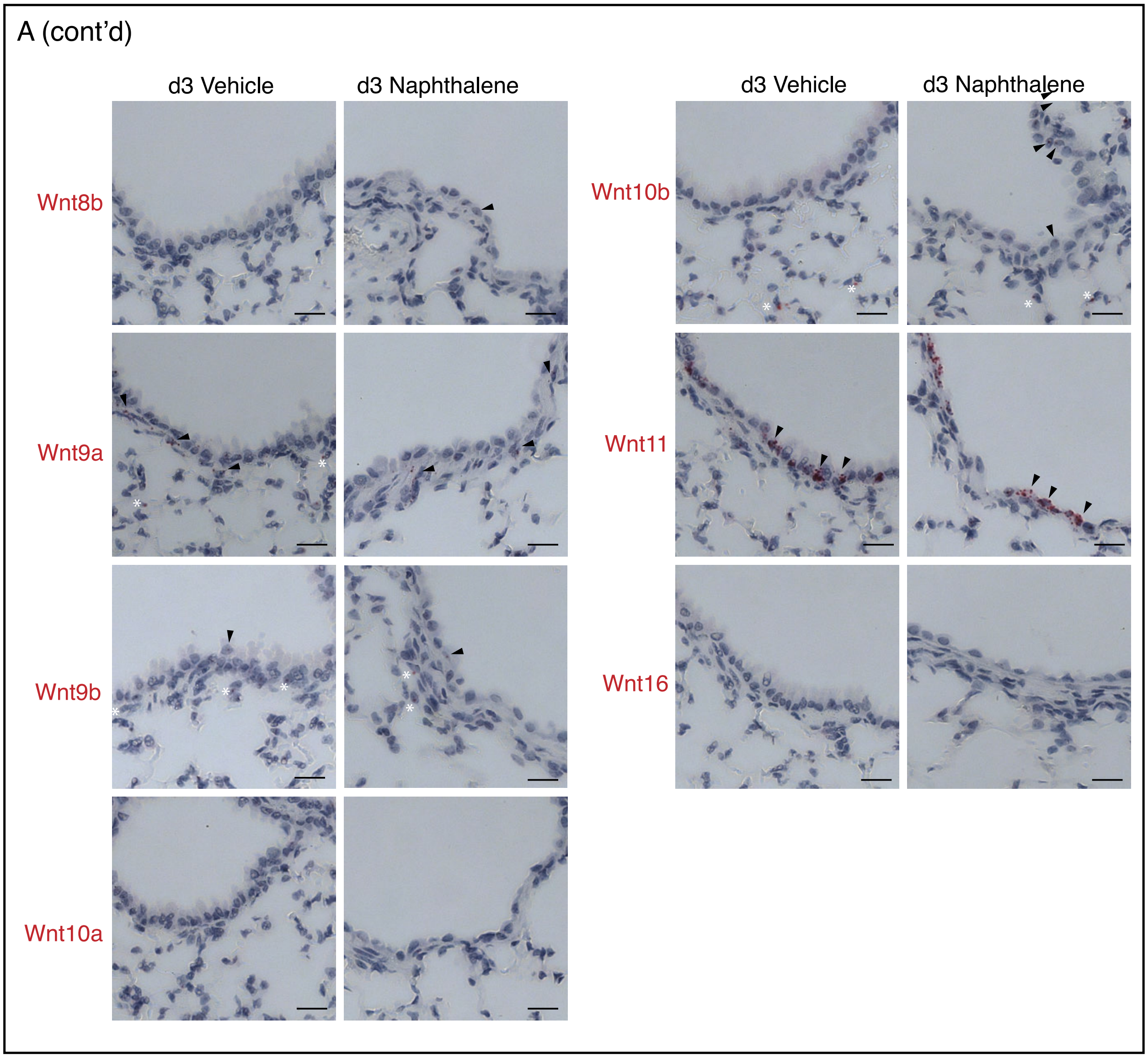
A (cont'd)
d3 Vehicle
d3 Naphthalene
d3 Vehicle
d3 Naphthalene
Wnt8b
Wnt9a
Wnt9b
Wnt10a
Wnt10b
Wnt11
Wnt16

Fig. S1 cont'd

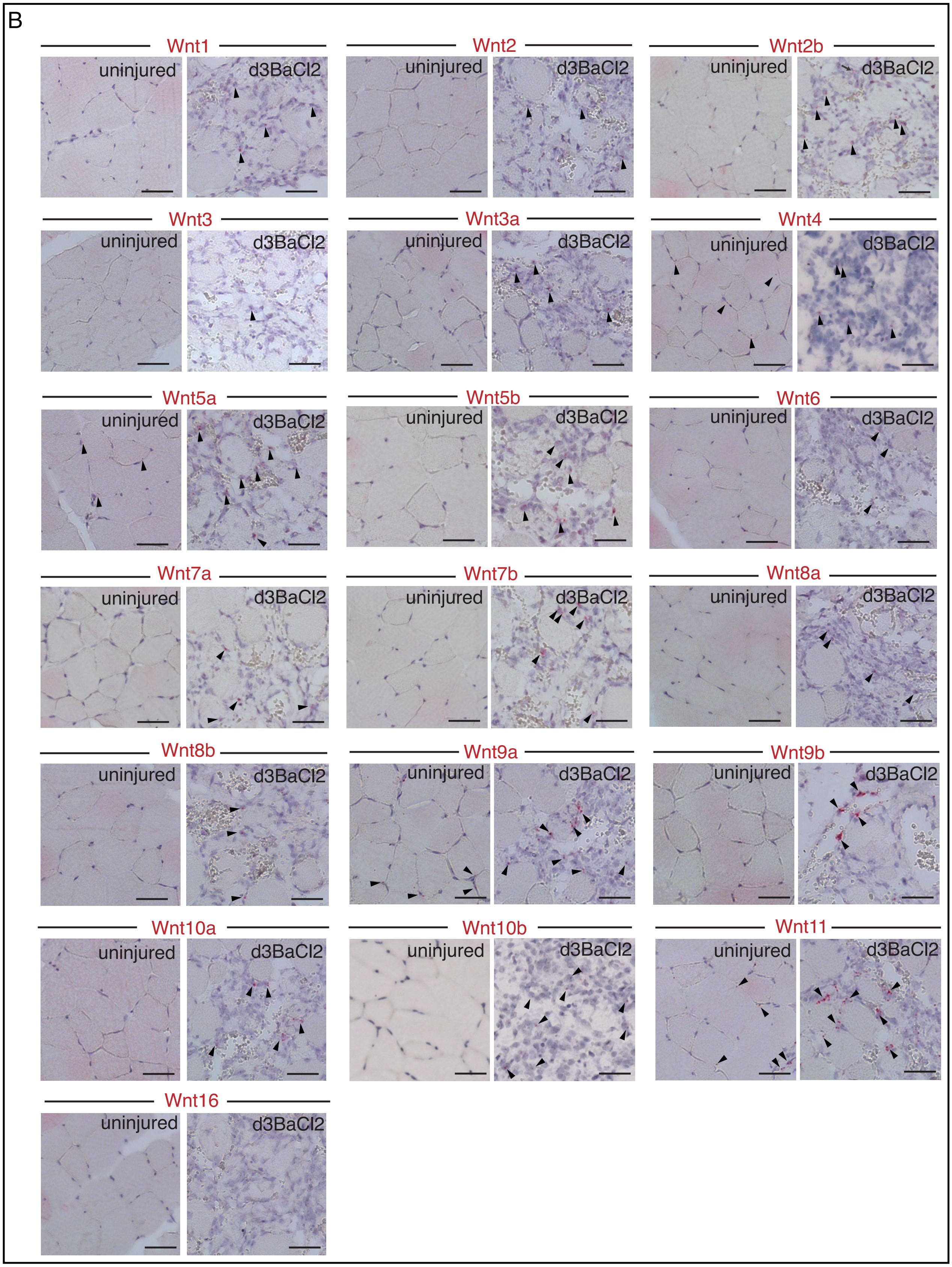
B
Wnt1
uninjured
d3BaCl2
Wnt2
uninjured
d3BaCl2
Wnt2b
uninjured
d3BaCl2
Wnt3
uninjured
d3BaCl2
Wnt3a
uninjured
d3BaCl2
Wnt4
uninjured
d3BaCl2
Wnt5a
uninjured
d3BaCl2
Wnt5b
uninjured
d3BaCl2
Wnt6
uninjured
d3BaCl2
Wnt7a
uninjured
d3BaCl2
Wnt7b
uninjured
d3BaCl2
Wnt8a
uninjured
d3BaCl2
Wnt8b
uninjured
d3BaCl2
Wnt9a
uninjured
d3BaCl2
Wnt9b
uninjured
d3BaCl2
Wnt10a
uninjured
d3BaCl2
Wnt10b
uninjured
d3BaCl2
Wnt11
uninjured
d3BaCl2
Wnt16
uninjured
d3BaCl2

Fig. S1 cont'd

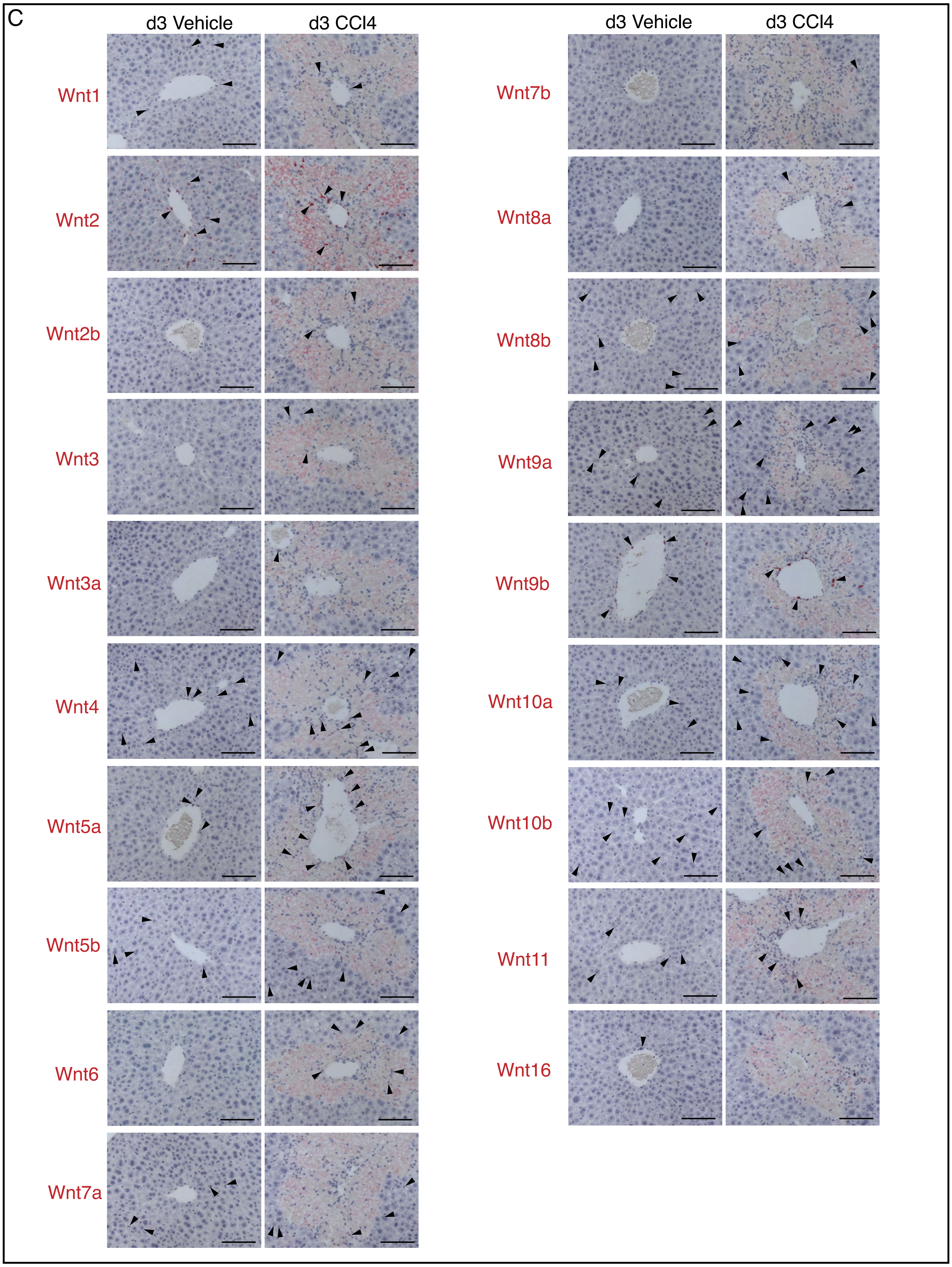
C
d3 Vehicle
d3 CCl4
d3 Vehicle
d3 CCl4
Wnt1
Wnt2
Wnt2b
Wnt3
Wnt3a
Wnt4
Wnt5a
Wnt5b
Wnt6
Wnt7a
Wnt7b
Wnt8a
Wnt8b
Wnt9a
Wnt9b
Wnt10a
Wnt10b
Wnt11
Wnt16

Fig. S1 cont'd

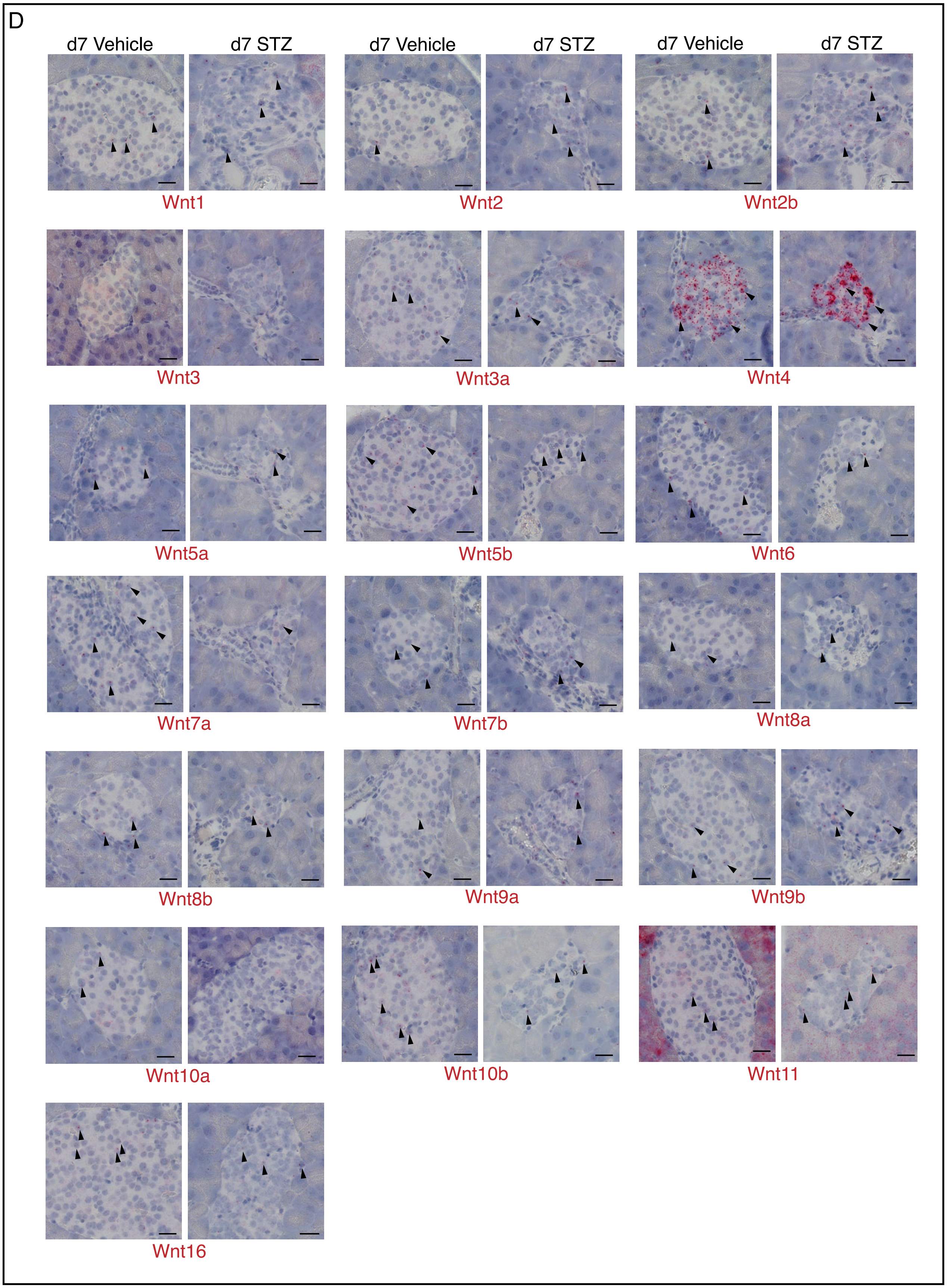
D
d7 Vehicle
d7 STZ
d7 Vehicle
d7 STZ
d7 Vehicle
d7 STZ
Wnt1
Wnt2
Wnt2b
Wnt3
Wnt3a
Wnt4
Wnt5a
Wnt5b
Wnt6
Wnt7a
Wnt7b
Wnt8a
Wnt8b
Wnt9a
Wnt9b
Wnt10a
Wnt10b
Wnt11
Wnt16

Fig. S1 cont'd

**Fig. S1: mRNA in situ hybridization screen for all 19 Wnt genes in 4 different injury models**

(A-D) Four different tissues were screened for Wnt expression before and after injury at 3 days for lung, muscle, and liver, and at 7 days for pancreas by mRNA in situ hybridization. A single representative region is shown for each of the 19 Wnts. Examples of mRNA signals, which appear as red dots, are marked by arrowheads. (A) Lungs were injured with Naphthalene and the airway epithelium or regions immediately below them were analyzed for changes in gene expression. Gene expression changes in the alveolar regions was not scored, but examples of signals are marked by asterisks. (B) Muscles were analyzed at 3 days post-BaCl2 injury. (C) Liver was injured with CCl4 and analyzed at 3 days post-injury. Regions surrounding the central vein were examined for changes in gene expression. The faint pink color surrounding the central vein marks necrotic areas due to the injury, and this represents background staining. (D) Pancreas injured by Streptozotocin was analyzed at 7 days post-treatment. Only signals within islets were analyzed. n=2 mice per injury model. (Scale Bars: 20um (A, B, D), 100um (C))

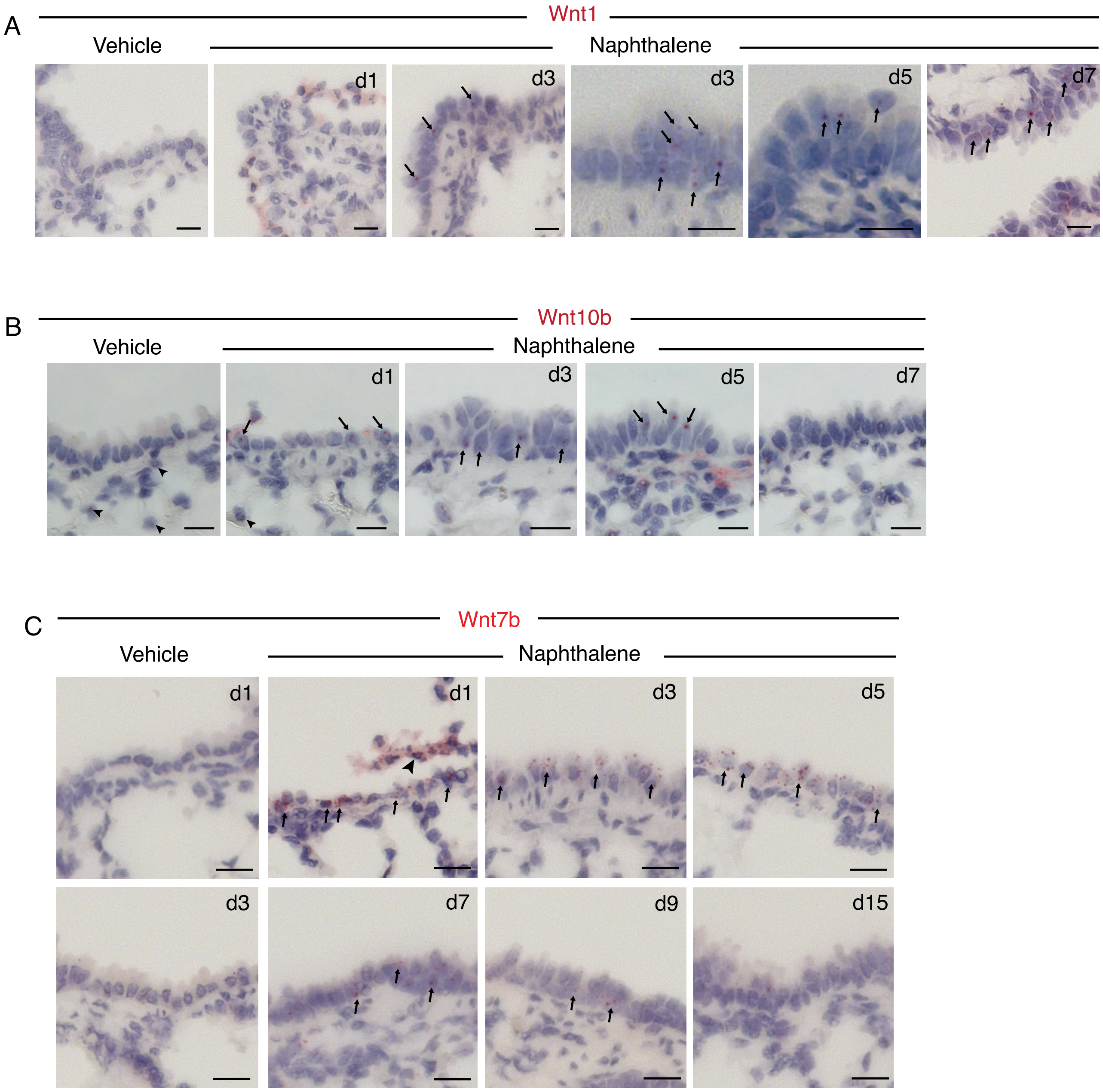
A
Wnt1
Vehicle
Naphthalene
d1
d3
d3
d5
d7
B
Wnt10b
Vehicle
Naphthalene
d1
d3
d5
d7
C
Wnt7b
Vehicle
Naphthalene
d1
d1
d3
d5
d3
d7
d9
d15

Fig. S2

**Fig. S2: mRNA in situ hybridization for Wnts induced following lung injury**

(A-C) Wnt mRNA in situ hybridization for three Wnts that are induced following Naphthalene injury in the lung at different timepoints compared to vehicle injury. (A) *Wnt1* mRNA expression in the lung at 1-7 days post injury shows that Wnt transcripts become visible starting at 3 days post-Naphthalene injection. Expression continues through 7 days. The representative vehicle injury shown is from a d1 vehicle-injected lung. Arrows point to red *Wnt1* transcript signals. (B) *Wnt10b* mRNA expression in the lung at 1-7 days post injury shows that *Wnt10b* transcripts become visible starting at 1 day post-Naphthalene injection. Expression continues through 7 days. Representative vehicle injury shown is from a d1 vehicle-injected lung. Arrows point to red *Wnt10b* transcript signals. In vehicle injected lungs, *Wnt10b* positive expression is found in regions underlying the airways (arrowheads) but is not found in the conducting airway epithelium. (C) *Wnt7b* mRNA expression in the lung at 1-15 days post injury shows that *Wnt7b* is highly expressed in the airway epithelium with transcripts visible at 1-day post-injury and continuing beyond 7 days. Examples of *Wnt7b* expression in vehicle-injected lungs are shown for day 1 and day 3. *Wnt7b* transcripts are not observed in uninjured airways. Arrowhead points to airway epithelial cells that have sloughed off due to Naphthalene injury. Arrows point to *Wnt7b* expression in epithelial cells. *Wnt7b* induction is consistent with previous studies (Volckaert et. al., 2011). Images are representative examples from n=3 mice. (Scale bars: 20um (A, B); 25um (C))

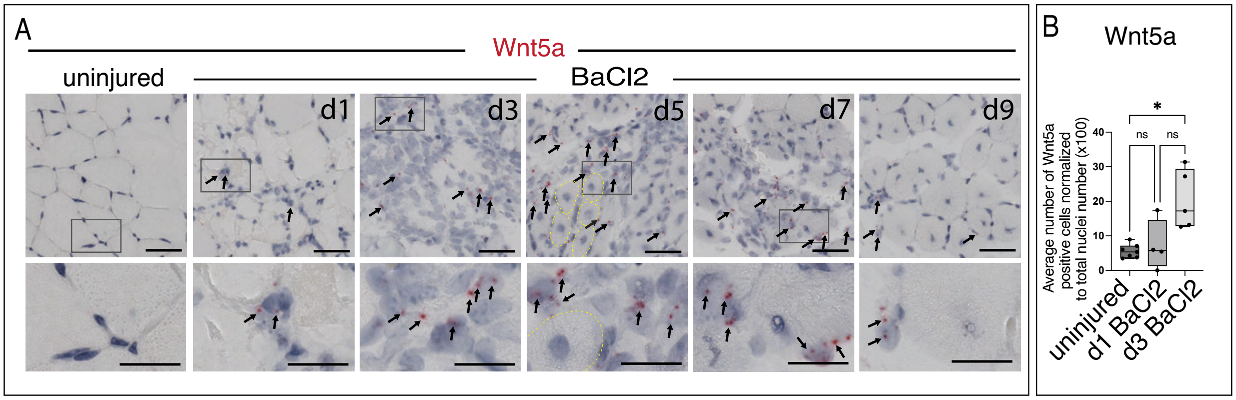

A
Wnt5a
uninjured
BaCl2
d1
d3
d5
d7
d9
B
Wnt5a
*
ns
ns
40
30
20
10
0
Average number of Wnt5a positive cells normalized to total nuclei number (x100)
uninjured
d1 BaCl2
d3 BaCl2


Fig. S3

**Fig. S3: mRNA in situ hybridization for *Wnt5a* following muscle injury**

(A) mRNA in situ hybridization for *Wnt5a* expression post-BaCl2 injury between days 1-9. Uninjured muscle is a representative contralateral muscle from BaCl2 injected mouse at 1-day post-injection. Grey boxes outline magnified areas shown below each panel. *Wnt5a* is rarely observed in uninjured muscles but is induced as early as 1-day post-injury and expression continues through 9 days. *Wnt5a* is predominantly found in cells that reside between regenerating myofibers and not in the nascent myofibers, which become evident by 5 days post-injury and can be identified by their centrally located nuclei. Yellow dotted lines mark examples of nascent myofibers. Arrows point to red mRNA transcript signals. (B) Quantification of *Wnt5a* transcripts shows that Wnt5a is gradually induced following injury between 1-3 days. Each dot represents the average expression of transcripts from 5-10 40x fields that were quantified for each mouse. Uninjured muscles represent combined data from 1 day (n=4) and 3 days post-injury (n=3). Day 1 injured muscles (n=4). Day 3 injured muscles (n=5). Two d3 uninjured muscles were lost during processing and are not included. (*P<0.05 (p=0.0471), Brown-Forsythe and Welch one-tailed ANOVA tests) Graphs show mean + s.d.

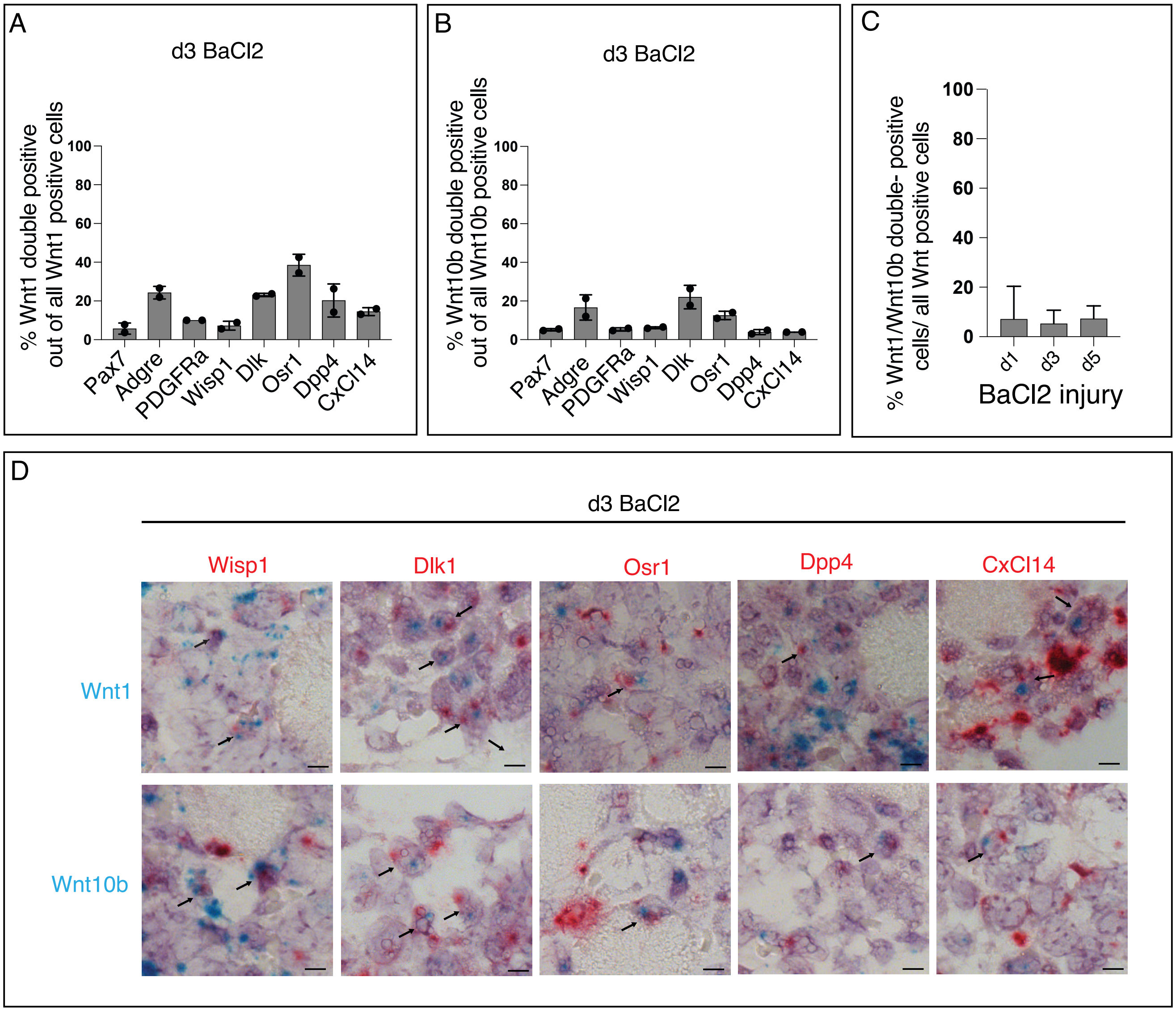
A
d3 BaCl2
% Wnt1 double positive out of all Wnt1 positive cells
Pax7
Adgre
PDGFRa
Wisp1
Dlk
Osr1
Dpp4
CxCl14
B
d3 BaCl2
% Wnt10b double positive out of all Wnt10b positive cells
C
% Wnt1/Wnt10b double- positive cells/ all Wnt positive cells
d1
d3
d5
BaCl2 injury
D
d3 BaCl2
Wisp1
Dlk1
Osr1
Dpp4
CxCl14
Wnt1
Wnt10b

Fig. S4

**Fig. S4: *Wnt1* and *Wnt10b* expression overlaps with multiple cell types including several sub-populations of FAPs**

(A, B) Bar graphs showing the percentage of *Wnt1* and *Wnt10b* positive nuclei that also express other markers of muscle cell types. In addition to *Pax7* and the *Adgre* gene, several FAP markers were examined. Y-axis values represent double-stained nuclei out of all *Wnt1* or *Wnt10b* positive cells that were scored expressed as a percentage. (A) *Wnt1* co-expression with multiple markers at 3 days post-injury. (B) *Wnt10b* co-expression with multiple markers at 3 days post-injury. (C) Bar graph showing the percentage of *Wnt1*-*Wnt10b* double-positive nuclei at 1-, 3-, and 5-days post-injury out of all Wnt-positive cells counted in each field. (D) Examples of nuclei co-expressing *Wnt1* or *Wnt10b* and various FAP markers. Arrows point to double-positive nuclei. (A-D, n=2 animals, A, B, n=6-8 fields for Pax7, Adgre, n=12-17 fields counted for FAP markers; C, n= 8 fields counted at d1 and d3 each, n=12 fields counted at d5.) Error bars are mean + s.d. (Scale bars: 5 um)

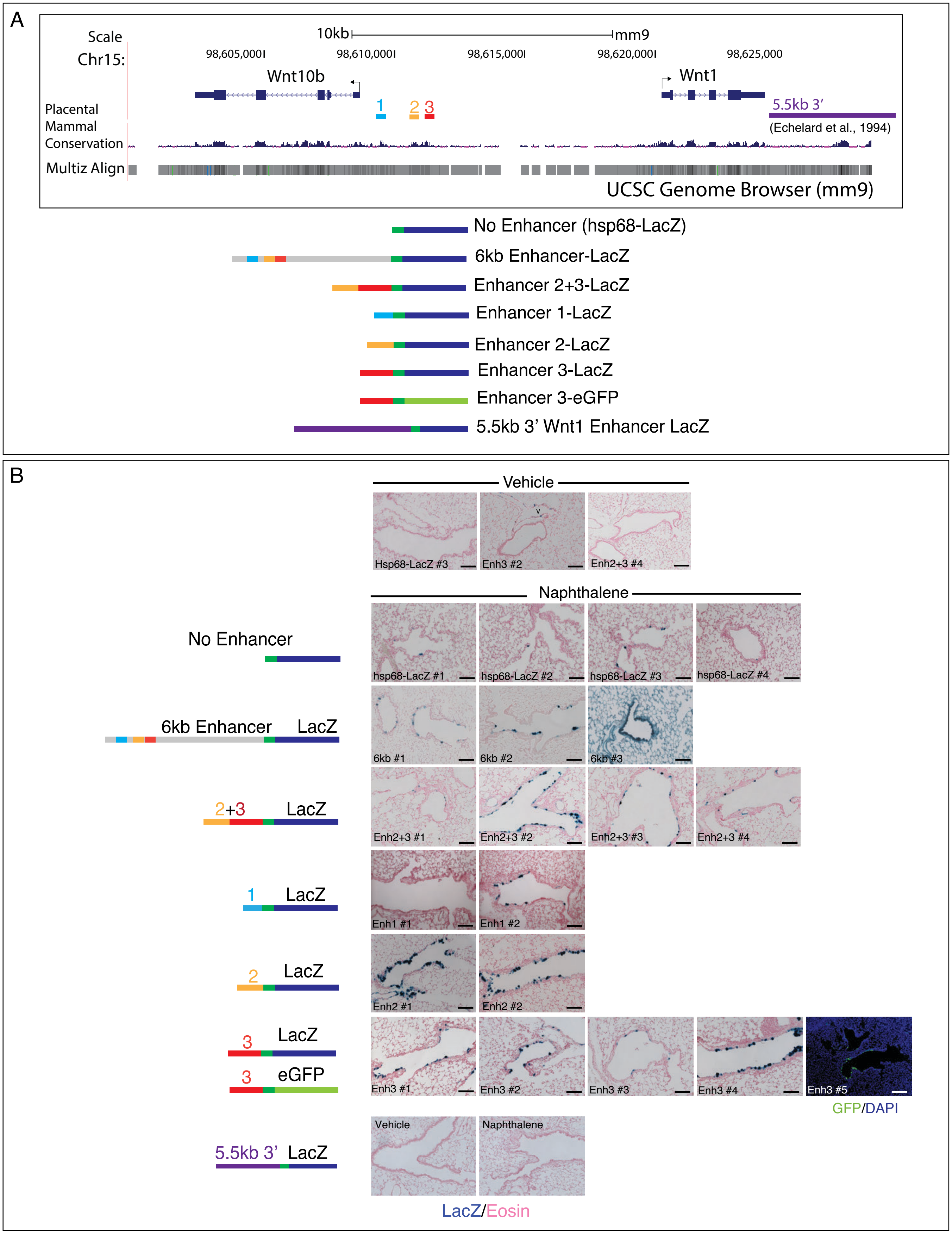
A
Scale
Chr15:
10kb
mm9
98,605,000
98,610,000
98,615,000
98,620,000
98,625,000
Wnt10b
Wnt1
1
2
3
5.5kb 3'
(Echelard et al., 1994)
Placental Mammal Conservation
Multiz Align
UCSC Genome Browser (mm9)
No Enhancer (hsp68-LacZ)
6kb Enhancer-LacZ
Enhancer 2+3-LacZ
Enhancer 1-LacZ
Enhancer 2-LacZ
Enhancer 3-LacZ
Enhancer 3-eGFP
5.5kb 3' Wnt1 Enhancer LacZ
B
Vehicle
Hsp68-LacZ #3
Enh3 #2
Enh2+3 #4
Naphthalene
No Enhancer
hsp68-LacZ #1
hsp68-LacZ #2
hsp68-LacZ #3
hsp68-LacZ #4
6kb Enhancer LacZ
6kb #1
6kb #2
6kb #3
2+3 LacZ
Enh2+3 #1
Enh2+3 #2
Enh2+3 #3
Enh2+3 #4
1 LacZ
Enh1 #1
Enh1 #2
2 LacZ
Enh2 #1
Enh2 #2
3 LacZ
3 eGFP
Enh3 #1
Enh3 #2
Enh3 #3
Enh3 #4
Enh3 #5
GFP/DAPI
5.5kb 3' LacZ
Vehicle
Naphthalene
LacZ/Eosin

Fig. S5

**Fig. S5: Generation of reporter mice to test the injury responsiveness of conserved regulatory sequences that reside between Wnt1 and Wnt10b**

(A)Schematic showing the genomic region between Wnt1 and Wnt10b genes in which 3 peaks of high conservation from the UCSC Genome browser are marked and numbered (blue box = putative enhancer region 1, yellow box = putative enhancer region 2, red box = putative enhancer region 3). The schematics below show these putative regulatory regions fused to an hsp68-LacZ construct (hsp promoter = dark green, LacZ = dark blue). As a control, a construct with only the hsp68-LacZ sequence without any enhancer sequence was generated. Additionally, a 6kb construct containing all three conserved enhancers (6kb Enhancer) plus a construct containing only enhancers 2 and 3 (2+3) were made. All other constructs carried one of the 3 putative enhancers fused to the reporter. An eGFP version of an Enhancer 3 reporter was generated as well. A previously identified regulatory region, located at the 3' end of the Wnt1 gene that drives expression in the embryonic CNS (Echelard et al., 1994) is also included. (B) X-gal-stained lung tissues to assess injury-responsiveness of the reporter constructs are shown. Each panel contains a representative example of lung tissue staining from an independently generated adult mouse line at the G2 generation that was injured with Naphthalene and analyzed at 5 days. Schematics of the constructs are shown next to the images and each independent line is numbered. Vehicle injected lungs displayed virtually no X-gal staining in any of the lines. Three different examples from different mouse lines are shown. Occasionally some LacZ expression in vessels could be observed (eg. Enh3 #2) but this was not consistent across all lines. Constructs with no enhancer gave only basal X-gal staining in rare cells within the airways but the presence of the 6kb enhancer and the 2+3 enhancer produced patches of X-gal stained cells within airways. Enhancer 1 also appeared to display an injury response, but this was only observed in 1 out of 2 lines. All lines generated from constructs containing Enhancer 2 or Enhancer 3 produced staining in airways post-injury. The Enhancer 3 eGFP- construct also displayed expression post-injury. In addition to our candidate enhancers, a Wnt1-LacZ mouse carrying a 3' Enhancer that drives in the embryonic CNS was also tested for injury-responsiveness, but no X-gal staining was observed in the vehicle or Naphthalene treated animals. n = $\geq$ 2 G2 generation mice for all constructs (Scale bars = 100 um)

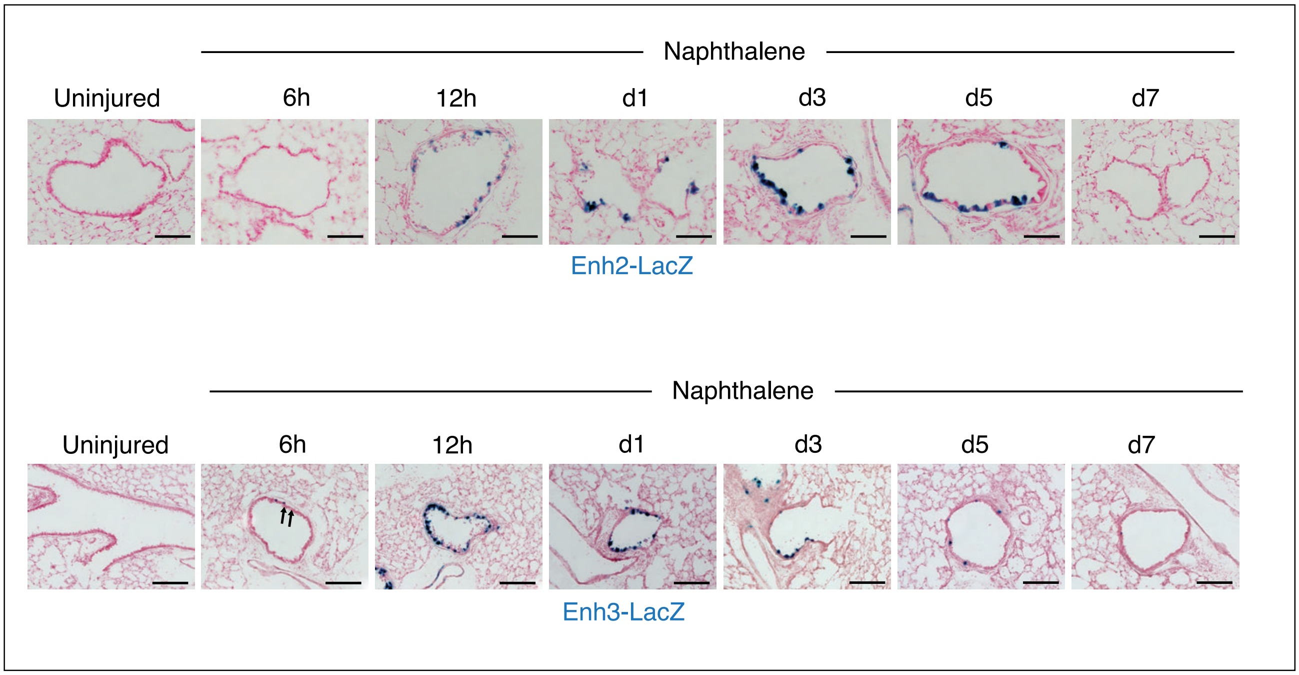
Naphthalene
Uninjured
6h
12h
d1
d3
d5
d7
Enh2-LacZ
Naphthalene
Uninjured
6h
12h
d1
d3
d5
d7
Enh3-LacZ

Fig. S6

**Fig. S6: Timing of Enhancer 2 and Enhancer 3 reporter expression in injured lungs**

Lungs from Naphthalene injured Enhancer 2 and Enhancer 3 mice were stained for X-gal on different days to assess the temporal dynamics of reporter activity. Staining time was adjusted to ensure that X-gal staining in airways did not reach saturation. Enhancer 3 reporter mice displayed detectable X-gal staining by 6 hours post injury (arrows) which peaked between 12h to 1 day and declined thereafter. Enhancer 2 reporter mice display detectable X-gal staining by 12 hours with levels peaking between 3-5 days. Representative examples of uninjured lungs are shown for both Enhancer 2 and 3. n=3 mice for all timepoints with consistent results between animals. (Scale bars: 100 um)

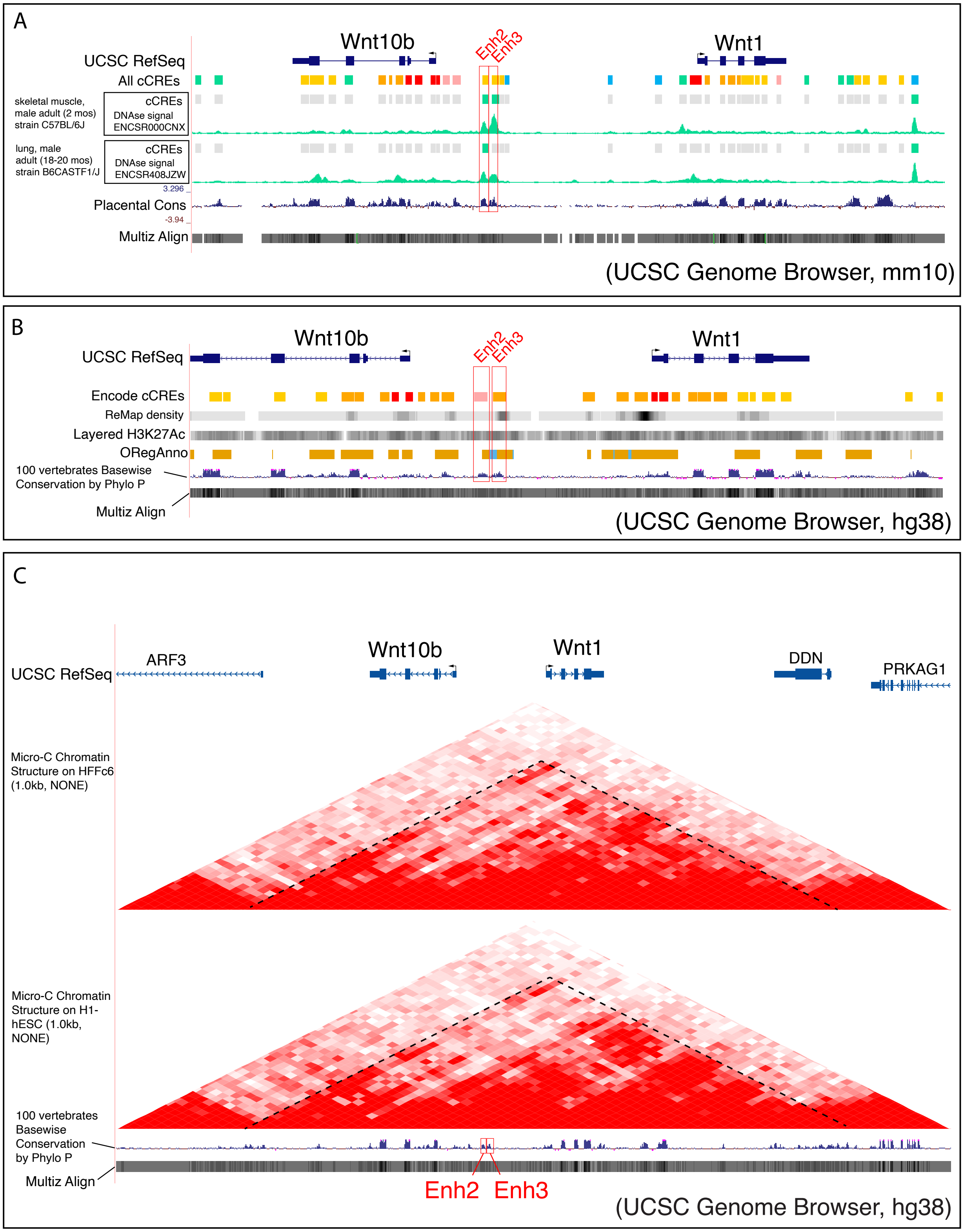
A
Wnt10b
Wnt1
Enh2
Enh3
UCSC RefSeq
All cCREs
skeletal muscle, male adult (2 mos) strain C57BL/6J
cCREs
DNAse signal ENCSR000CNX
lung, male adult (18-20 mos) strain B6CASTF1/J
cCREs
DNAse signal ENCSR408JZW
3.296
Placental Cons
-3.94
Multiz Align
(UCSC Genome Browser, mm10)
B
Wnt10b
Wnt1
Enh2
Enh3
UCSC RefSeq
Encode cCREs
ReMap density
Layered H3K27Ac
ORegAnno
100 vertebrates Basewise Conservation by Phylo P
Multiz Align
(UCSC Genome Browser, hg38)
C
UCSC RefSeq
ARF3
Wnt10b
Wnt1
DDN
PRKAG1
Micro-C Chromatin Structure on HFFc6 (1.0kb, NONE)
Micro-C Chromatin Structure on H1-hESC (1.0kb, NONE)
100 vertebrates Basewise Conservation by Phylo P
Multiz Align
Enh2
Enh3
(UCSC Genome Browser, hg38)

Fig. S7

**Fig. S7: Examples of publicly available data that suggest that Enhancers 2 and 3 can function as regulatory regions.**

(A) Output from the UCSC Genome Browser (mm10) to look for candidate cis-regulatory sequences (cCREs) and signatures of open chromatin shows that Enhancers 2 and 3 could be cis-regulatory regions in both lung and muscle. Both enhancer sites display peaks of DNAse1 hypersensitivity (red boxes marking green peaks). The Enhancer 2 and 3 regions, which can be identified by their conserved peaks (blue peaks, Placental Cons.) on multi-species alignments are also marked (red boxes). (B) Output from the UCSC Genome Browser (hg38) summarizes data from ENCODE, ReMap and ORegAnno outputs and H3K27Ac marks to suggest that Enhancers 2 and 3 could be functional enhancers. The Enhancer 2 and 3 regions, which can be identified by their conserved peaks (blue peaks) on multi-species alignments (100 vertebrates Basewise Conservation by Phylo P and Multiz Align) are outlined by red boxes. The ENCODE data suggests that Enhancer 2 and Enhancer 3 display DNase-H3K4me3 activity (pink box), and a distal enhancer-like signature (orange box) respectively. ReMap, which integrates multiple CHIP-seq and other datasets to output possible regulatory elements shows higher density signals in Enhancer 3. H3K27Ac marks are also seen within Enhancers 2 and 3. The ORegAnno output indicates that Enhancer 2 and 3 regions harbor transcription factor binding sites (orange boxes), and that Enhancer 3 is a possible regulatory region (light blue box). (C) Outputs from the UCSC Genome Browser (hg38) showing Micro-C Chromatin assays to identify possible contact points in and around *Wnt1* and *Wnt10b* suggests that this region could form a topologically associated domain. The overall boundaries spanning *Wnt1* and *Wnt10b* are marked with a dotted line. The locations of Enhancers 2 and 3 are outlined with red boxes where peaks of high conservation are observed (100 vertebrates Basewise Conservation by Phylo P and Multiz Align). Data are from human foreskin fibroblasts (HFFc6 cells) and human embryonic stem cells (H1-hESCs).

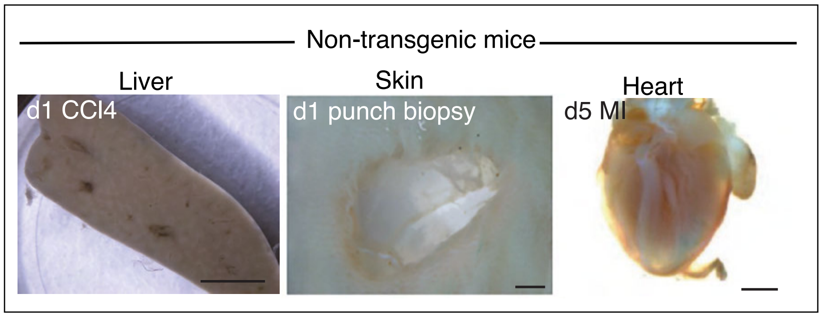
Non-transgenic mice
Liver
Skin
Heart
d1 CCl4
d1 punch biopsy
d5 MI

Fig. S8

**Fig. S8: Non-transgenic tissues do not express LacZ staining**

Examples of injured liver ($CCl_4$ at d1), skin (punch biopsy at d1), and heart (myocardial infarction (MI) at d5) show that non-transgenic animals do not stain for LacZ. (Scale bars: 2 mm liver, skin; 1 mm heart)

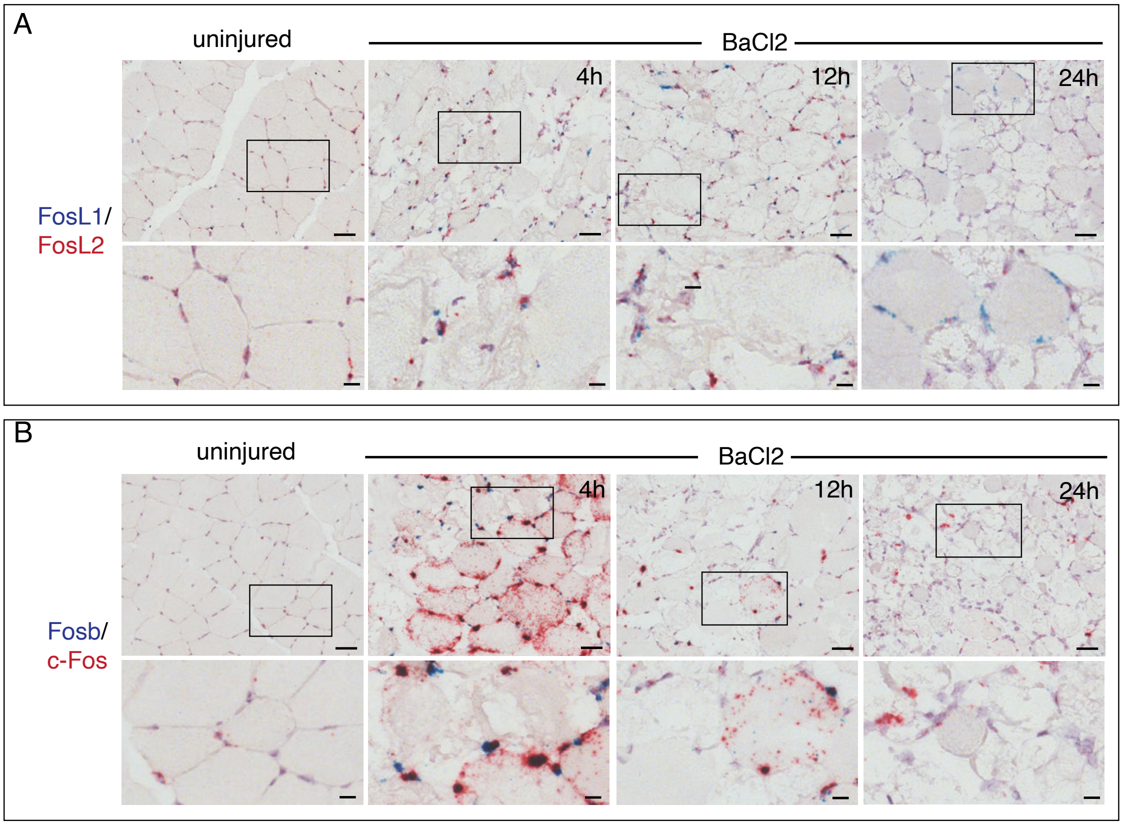
A
uninjured
BaCl2
4h
12h
24h
FosL1/
FosL2
B
uninjured
BaCl2
4h
12h
24h
Fosb/
c-Fos

Fig. S9

**Fig. S9: Members of the AP-1 family of genes are expressed in muscles following BaCl2 injury**

(A, B) Duplex mRNA in situ hybridization to examine the expression of several Fos-family members at 4, 12, and 24 hours post-BaCl2 injury. Grey boxes mark regions that are enlarged and shown below. (A) *FosL1* and *FosL2* staining shows that *FosL2* is present in uninjured muscles, but injury induces the expression of *FosL1* which is detectable at 4 hours and continues for at least 24 hours. *FosL2* is present at all timepoints but may decline at 24 hours, during the time when the myofibers are clearly degenerating. (B) *Fosb* and *c-Fos* staining shows *c-fos* is expressed in uninjured muscles but dramatically increases levels following injury at 4 hours and declines rapidly. *Fosb* is also transiently up regulated at 4-12 hours and then declines. n=2 mice for each timepoint with consistent results between animals. (Scale Bars: 20 um for top row of images, 5 um for enlarged insets)

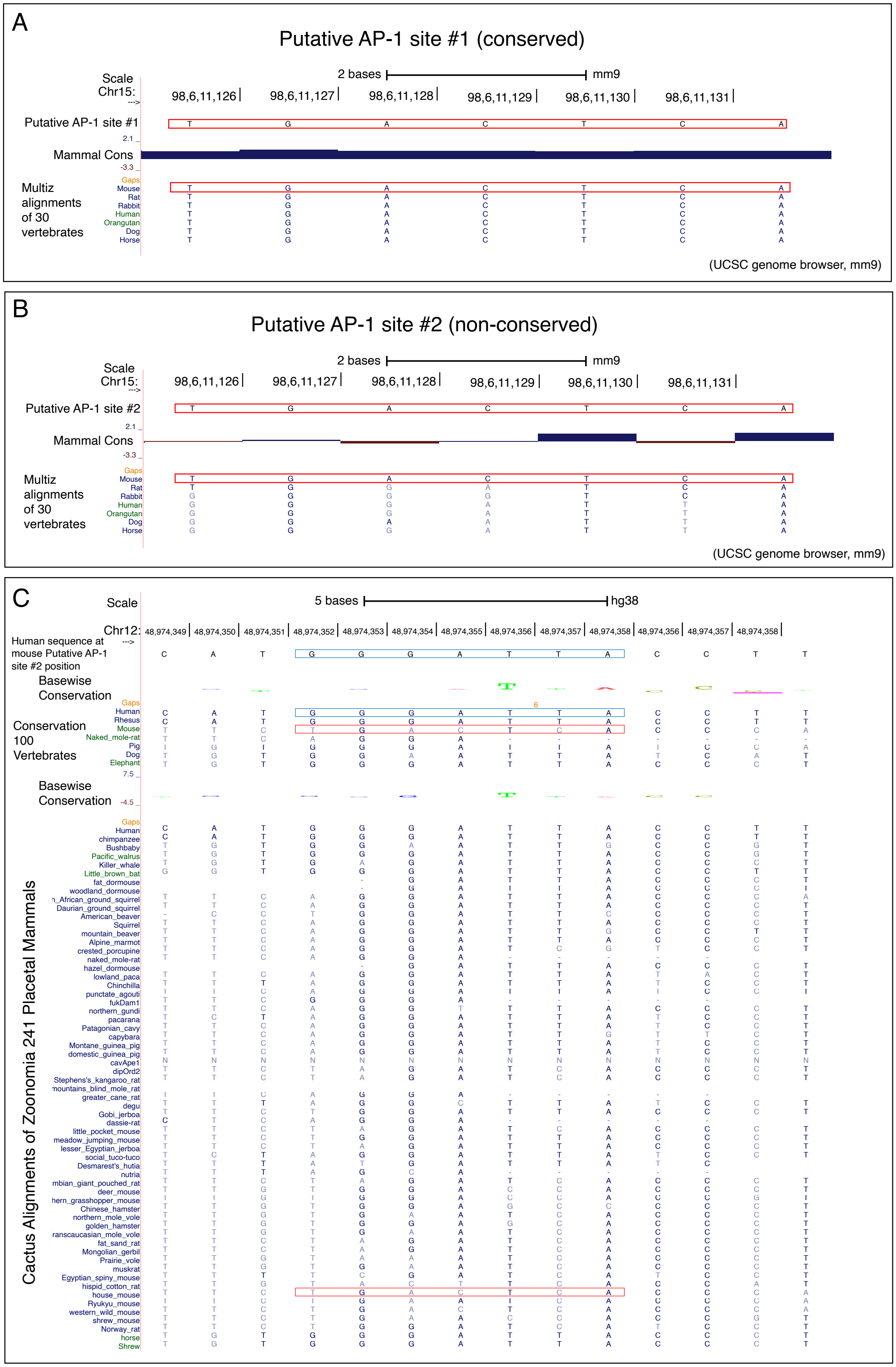
A
Putative AP-1 site #1 (conserved)
Scale
Chr15:
2 bases
mm9
98,6,11,126
98,6,11,127
98,6,11,128
98,6,11,129
98,6,11,130
98,6,11,131
Putative AP-1 site #1
Mammal Cons
Multiz alignments of 30 vertebrates
Gaps
Mouse
Rat
Rabbit
Human
Orangutan
Dog
Horse
(UCSC genome browser, mm9)
B
Putative AP-1 site #2 (non-conserved)
Scale
Chr15:
2 bases
mm9
98,6,11,126
98,6,11,127
98,6,11,128
98,6,11,129
98,6,11,130
98,6,11,131
Putative AP-1 site #2
Mammal Cons
Multiz alignments of 30 vertebrates
(UCSC genome browser, mm9)
C
Scale
5 bases
hg38
Chr12:
Human sequence at mouse Putative AP-1 site #2 position
Basewise Conservation
Conservation 100 Vertebrates
Human
Rhesus
Mouse
Naked_mole-rat
Pig
Dog
Elephant
Cactus Alignments of Zoonomia 241 Placetal Mammals

Fig. S10

**Fig. S10: Two Putative AP-1 binding sites are present in mouse Enhancer 3, but one of them is not conserved**

(A, B) Alignments using the UCSC Genome Browser (mm9) to examine putative AP-1 binding sites identified in Enhancer 3 show that only one of the two sites is highly conserved. The consensus binding sequence TGAG/CTCA and alignments with other species are shown. Rat, rabbit, human, orangutan, dog and horse are listed for comparison. The mouse sequence is outlined by a red box. (A) Putative AP-1 binding site (referred to as site #1) is highly conserved. (B) Putative AP-1 binding site (referred to as site #2) is not highly conserved and in other species does not match the consensus sequence. (C) A more detailed multi-species alignment using the UCSC Genome Growser (hg38) that includes selections from the 100 vertebrates alignment and the 241 placental mammals alignment shows that the putative AP-1 binding site #2 sequence is unique to the house mouse and is not conserved. Alignment of mainly rodent sequences is shown here. Human sequence is outlined by a blue box. The mouse sequence is outlined by a red box.

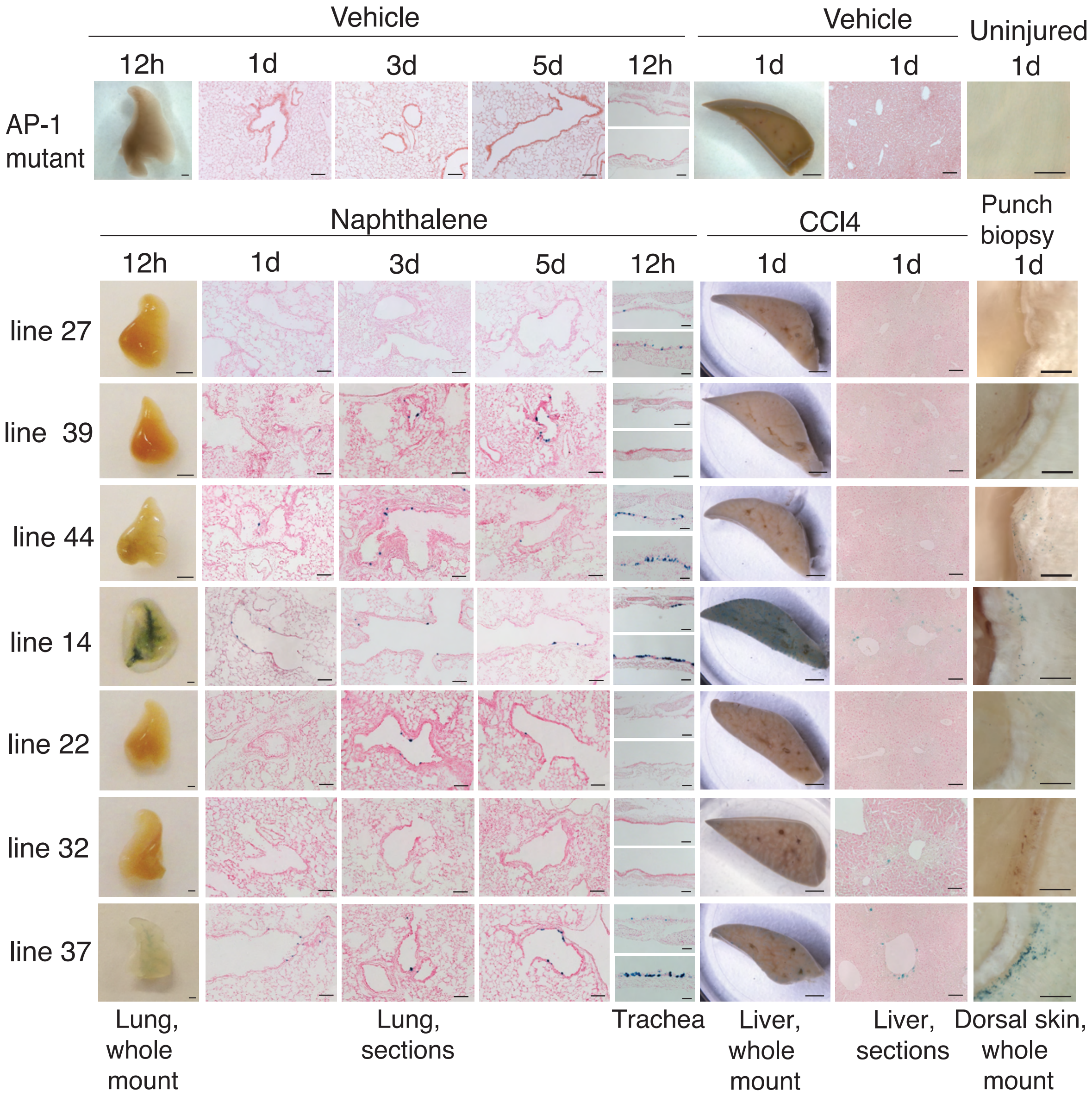
Vehicle
12h
1d
3d
5d
12h
Vehicle
1d
1d
Uninjured
1d
AP-1
mutant
Naphthalene
CCl4
Punch
biopsy
1d
12h
1d
3d
5d
12h
1d
1d
line 27
line 39
line 44
line 14
line 22
line 32
line 37
Lung,
whole
mount
Lung,
sections
Trachea
Liver,
whole
mount
Liver,
sections
Dorsal skin,
whole
mount

Fig. S11

**Fig. S11: Enhancer 3 reporter lines carrying mutated putative AP-1 binding sites fail to display reporter activity in multiple tissues following injury**

Mutating the putative AP-1 binding sites found in Enhancer 3 causes loss of reporter activity post-injury. 7/8 lines that were not shown in Figure 3 are shown here (lines 27, 39, 44, 14, 22, 32, 37). These lines display lack of or reduced LacZ expression following injury in the lung, liver and skin. Representative examples of staining in uninjured mutant AP-1 tissues are also included (from Line 21 and 44). All uninjured tissues from AP-1 mutant reporter mice failed to stain with X-Gal. Tissues shown here are whole mounts (lung, 12h, Naphthalene; liver, 1d, CCl4; skin, d1, punch biopsy) and tissue sections (lung, d1, 3, 5, Naphthalene; trachea 12h, Naphthalene; liver, d1, CCl4). Pink counterstain is Eosin. Scale bars: 100 um lung, trachea, liver sections, 2 mm lung, liver, skin whole mounts)

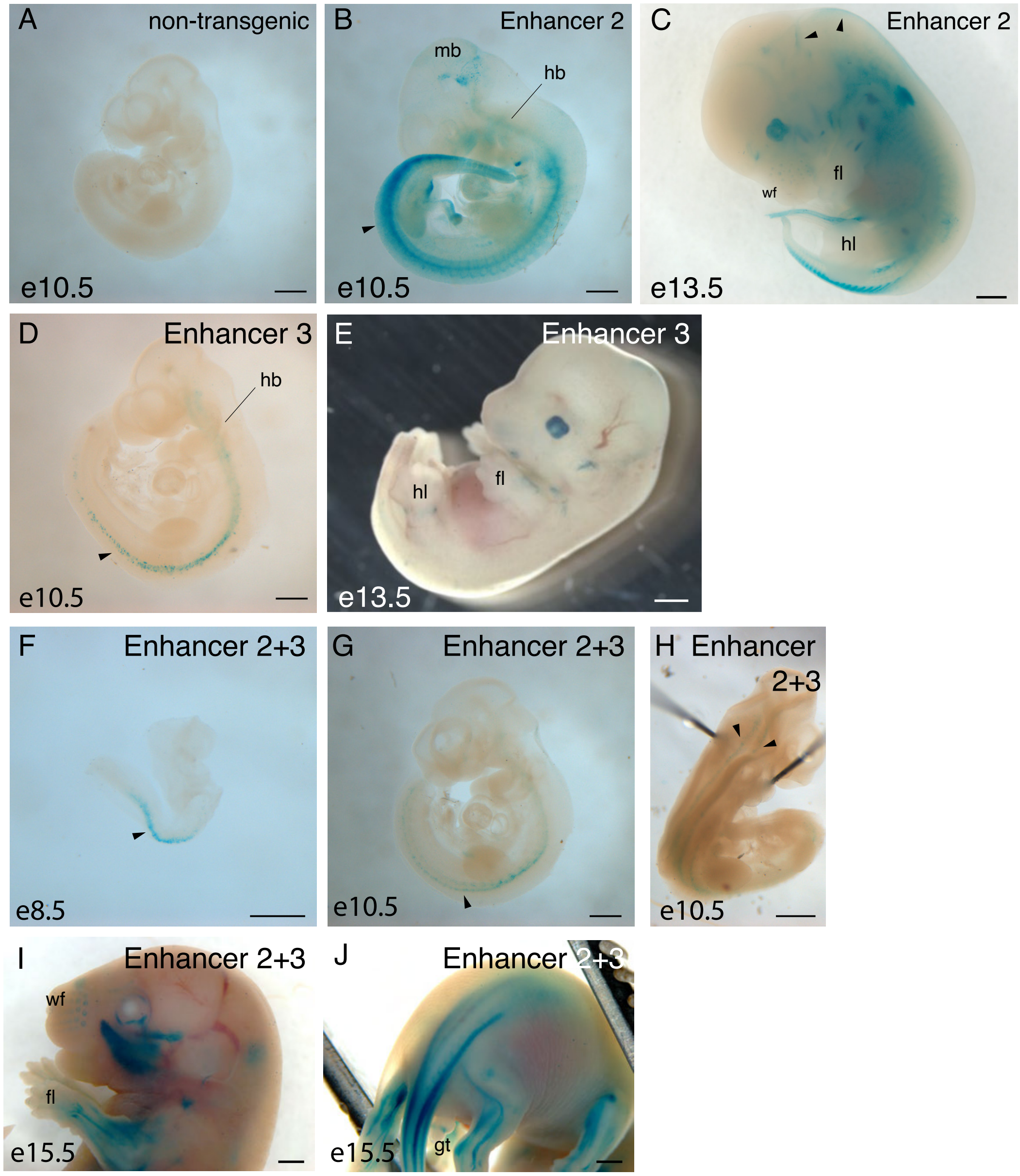

A
non-transgenic
e10.5
B
Enhancer 2
mb
hb
e10.5
C
Enhancer 2
wf
fl
hl
e13.5
D
Enhancer 3
hb
e10.5
E
Enhancer 3
hl
fl
e13.5
F
Enhancer 2+3
e8.5
G
Enhancer 2+3
e10.5
H
Enhancer 2+3
e10.5
I
Enhancer 2+3
wf
fl
e15.5
J
Enhancer 2+3
gt
e15.5


Fig. S12

**Fig. S12 : Enhancer 2, 3, and 2+3 LacZ reporters are expressed in embryos**

(A) A non-transgenic e10.5 embryo showing that animals that lack reporter constructs do not stain for LacZ. (B) Enhancer 2-LacZ embryos at e10.5 display LacZ staining in multiple locations including ventral regions along the anterior-posterior axis (arrowhead) as well as in the brain. (C) At e13.5, Enhancer 2-LacZ embryos display staining in multiple locations that include the tail, forelimbs, whisker follicles, and multiple sites in the head (arrowheads). (D) Enhancer 3-LacZ embryos at e10.5 display scattered cells in two stripes along the body axis towards the ventral side of the CNS (arrowhead). (E) By e13.5, staining is mostly observed on the surface of the skin in patches on the forelimbs and hindlimbs. (F) Enhancer 2+3 embryos at e8.5 display a ventral stripe of LacZ staining (arrowhead). (G) At e10.5 Enhancer 2+3 embryos continue to display staining along the ventral CNS. (H) By viewing LacZ stained Enhancer 2+3 embryos at e10.5 before neural tube closure, it is evident that the ventral reporter expression is comprised of two stripes of cells (arrowheads) that lie along the ventral neural tube. (I) By e15.5 Enhancer 2+3-LacZ embryos display multiple sites of LacZ staining throughout the face, head and forelimbs. (J) A view of the posterior region of an e15.5 Enhancer 2+3-LacZ reporter mouse to show staining in limbs, tail and genital tubercle. (mb, midbrain, hb, hindbrain; fl, forelimb, hl, hindlimb, wf, whisker follicles, gt, genital tubercle) (Scale bars: 0.5 mm (A-E), 1 mm (F-J))

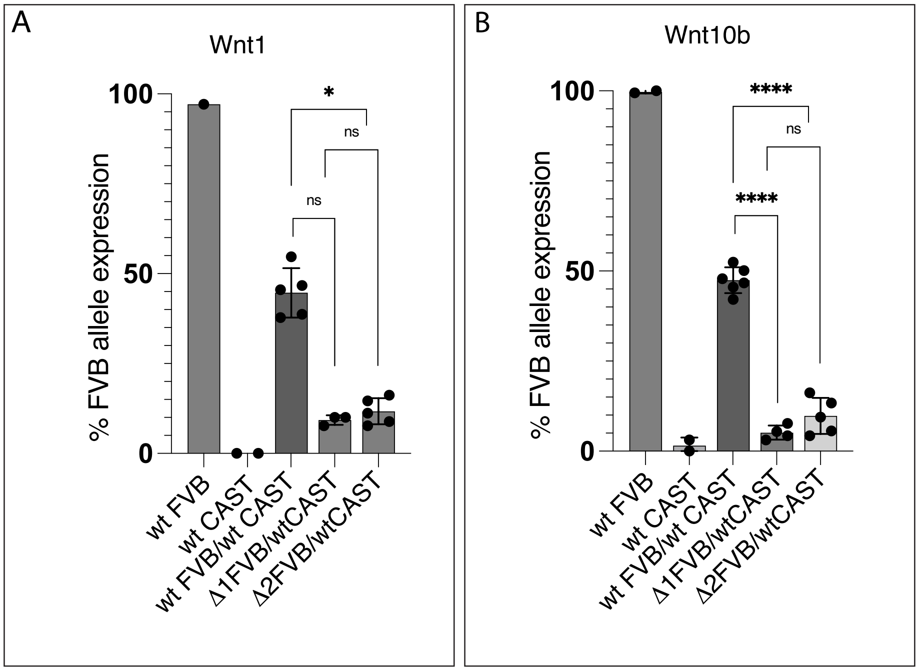
A
Wnt1
100
50
0
% FVB allele expression
*
ns
ns
wt FVB
wt CAST
wt FVB/wt CAST
Δ1FVB/wtCAST
Δ2FVB/wtCAST
B
Wnt10b
100
50
0
% FVB allele expression
****
ns
****
wt FVB
wt CAST
wt FVB/wt CAST
Δ1FVB/wtCAST
Δ2FVB/wtCAST

Fig. S13

**Fig. S13: Pyrosequencing data at 3 days post-injury show that the alleles carrying deletions of Enhancers 2 and 3 are not efficiently expressed**

(A, B) Bar graphs showing *Wnt1* and *Wnt10b* expression in which expression of wildtype vs. mutant (Δ1 or Δ2) alleles can be distinguished by specific SNPs found only in either FVB or Castaneous (CAST/EIJ) mice. Percent expression of the allele specific to the FVB strain is shown. Animals were analyzed at 3 days post-BaCl2 injury. Uninjured muscles are not included. Graphs show % FVB allele expression but the statistical analysis was performed on arcsine transformed values. (A) Percent expression of *Wnt1* from the FVB-specific SNP is shown in mice that are either 100% wildtype FVB (n=2), 100% wild-type CAST (n=2), 50%/50% wild-type FVB/wild-type CAST (n=5), or 50% Δ1 (n=3) or Δ2 (n=5) FVB/wild-type CAST. As expected, SNP expression specific for the FVB allele is detected at nearly 100% in wild-type FVB mice, whereas it is almost undetectable in 100% CAST/EIJ muscles. When both FVB and CAST alleles are wild-type (50%/50%), expression of the FVB-specific *Wnt1* SNP is expressed at close to 50%. Δ1 and Δ2 alleles display much lower expression. Because we had insufficient numbers of 100% FVB and 100% CAST samples, statistical analysis was performed only on the wild-type FVB/wild-type CAST, Δ1 FVB/wild-type CAST, and Δ2 FVB/wild-type CAST samples. In Δ2 muscles, expression of the FVB allele is significantly lower than of the wild type allele. Δ1 muscles also display a similarly low expression but analysis did not show statistical significance likely due to the smaller sample size. (Kruskal Wallis Test, *$p<0.005$ ($p=0.0242$)) Graphs show mean ± s.d. (B) Percent expression of *Wnt10b* from the FVB-specific SNP is shown in mice that are either 100% wildtype FVB (n=1), 100%wild-type CAST (n=2), 50%/50% wild-type FVB/wild-type CAST (n=6), or 50% Δ1 (n=4) or Δ2 (n=5) FVB/wild-type CAST. As expected, SNP expression specific for the FVB allele is detected at nearly 100% in wild-type FVB mice, whereas it is almost undetectable in 100% CAST muscles. When both FVB and CAST alleles are wild-type (50%/50%), expression of the FVB-specific Wnt1 SNP is expressed at close to 50%. Δ1 and Δ2 alleles display much lower expression. Statistical analysis was performed only on the wild-type FVB/wild-type CAST, Δ1 FVB/wild-type CAST, and Δ2 FVB/wild-type CAST samples. In muscles carrying Δ1 or Δ2 alleles, reduced Wnt expression from the mutant FVB allele compared to wild-type is highly significant. (Brown-Forsythe and Welch one-tailed ANOVA tests, ****P<0.0001) Graphs show mean ± s.d.

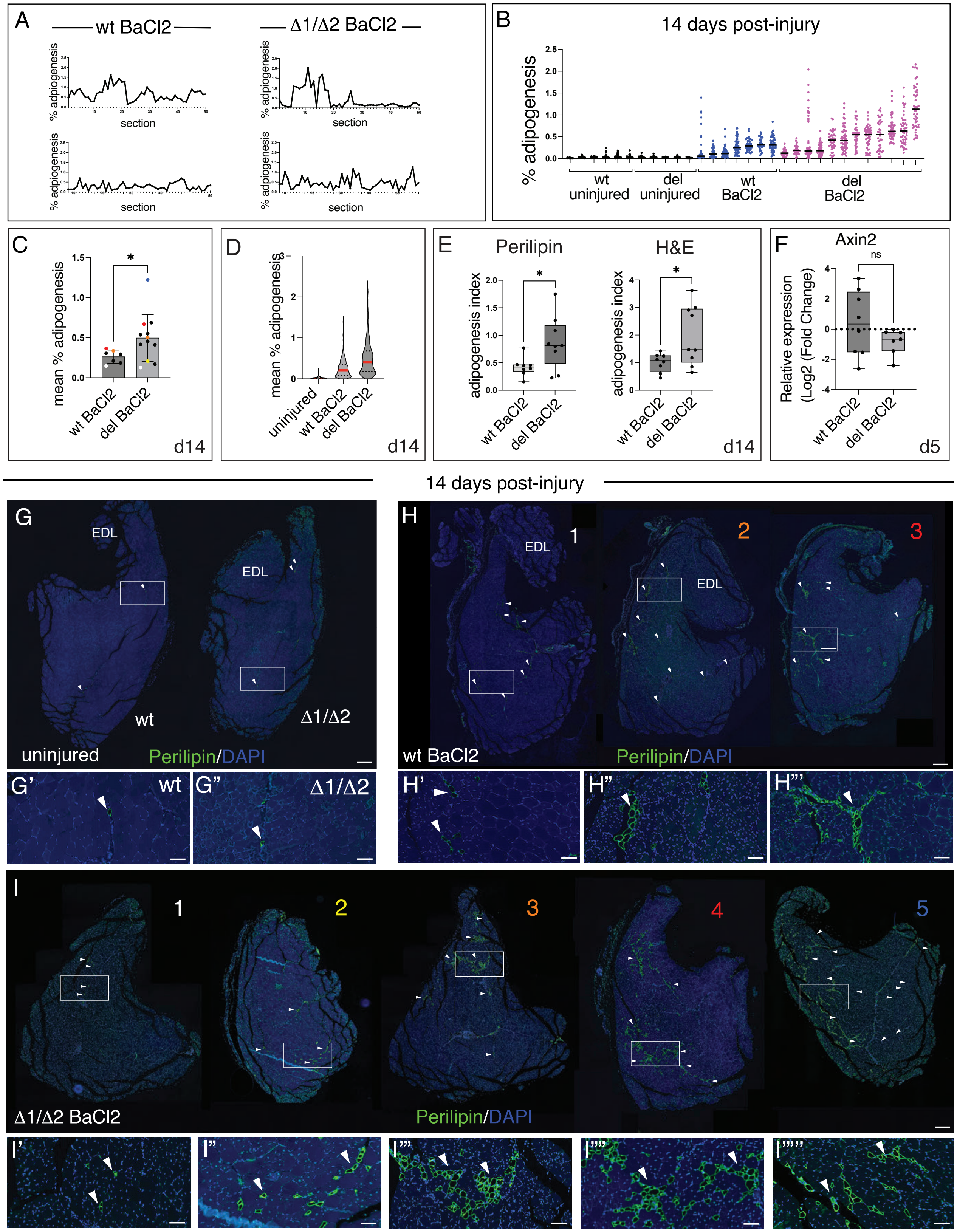
A
wt BaCl2
Δ1/Δ2 BaCl2
% adipogenesis
section
B
14 days post-injury
% adipogenesis
wt uninjured
del uninjured
wt BaCl2
del BaCl2
C
mean % adipogenesis
wt BaCl2
del BaCl2
d14
D
mean % adipogenesis
uninjured
wt BaCl2
del BaCl2
d14
E
Perilipin
H&E
adipogenesis index
wt BaCl2
del BaCl2
d14
F
Axin2
ns
Relative expression (Log2 (Fold Change)
wt BaCl2
del BaCl2
d5
14 days post-injury
G
EDL
wt
Δ1/Δ2
uninjured
Perilipin/DAPI
H
1
2
3
wt BaCl2
Perilipin/DAPI
G'
G"
H'
H"
H'''
I
1
2
3
4
5
Δ1/Δ2 BaCl2
Perilipin/DAPI
I'
I"
I'''
I''''
I'''''

Fig. S14

**Fig. S14 Adipogenesis in BaCl2 injured muscles at 14 days post-injury**

(A)Line graphs showing % adipogenesis values obtained from BaCl2 injected TA muscles at 14 days post-injury over 50 serial sections to show that there are fluctuations in the level of fat observed across sections. Two examples from wt mice and two examples of Δ1/Δ2 muscles are shown. (B) Scatter plots showing the individual % adipogenesis measurements for each section analyzed in individual animals to show the spread of values that were observed at 14 days post-injury. There is a general upward trend in the distribution of values observed in injured Δ1/Δ2 muscles compared to wild type. Uninjured mice show very low % adipogenesis values regardless of genotype. Lines show the median. n=6 wt uninjured, n=5 Δ1/Δ2 uninjured, n=7 wt BaCl2, n=12 Δ1/Δ2 (C) Bar graph to show the mean % adipogenesis values for each wt vs. Δ1 /Δ2 animal. Only animals in which BaCl2 injury produced >45% damage in the tissue are shown. Colored dots on the graph correspond to examples of numbered mouse tissue sections with the matching color s shown in the panels D-F to correlate the plotted mean % adipogenesis values with the staining levels observed in tissue. n=7 wt, n=12 Δ1/Δ2 (*P<0.05, Welch's t-test) (D) Violin plot of mean % adipogenesis values from all fields quantified for Perilipin staining in uninjured wild-type muscles (n=181 fields, n=4 mice), injured wt muscles (n=375 fields, n=8 mice) and injured Δ1/Δ2 muscles (n=423 fields, n=12 mice) at 14-days post-BaCl2 injury to show the distribution of all samples analyzed. The level of adipogenesis observed is elevated in Δ1 /Δ2 muscles compared to wild type. 35-50 fields were counted per mouse. (E) A comparison of adipogenesis levels that are measured when fatty infiltration is evaluated by Perilipin vs. H&E staining. By both staining methods, a similar increase in adipogenesis can be observed in which injured Δ1 /Δ2 muscles display more adipogenesis than wt muscles (n=8 wt, n=9 Δ1 /Δ2 ; 40-50 fields per mouse were quantified for Perilipin staining; n=10 fields per mouse were quantified for H&E staining)(F) RT-qPCR of *Axin2* expression following injury in wt and Δ1 /Δ2 mice shows a slight decrease in expression level that might indicate reduced Wnt signaling when Enhancers 2 and 3 are deleted, but the difference is not statistically significant. n=8 wt, n=7 Δ1 /Δ2 (Welch's t-test, ns = not significant. (G-I) Representative examples of adipogenesis observed in sections of BaCl2 injected TA muscles at 14 days from wt and Δ1 /Δ2 animals that were stained with Perilipin1 antibody to mark adipocytes. Sections are arrayed on a single dark background separated by genotype (G), and/or treatment (H, I) for ease of viewing, but each section represents tissues from separate animals. Arrowheads point to patches of adipogenesis in the muscle. Grey boxes outline magnified regions that are shown in the row below. (G) Two representative examples from uninjured mice (n=1 wt, n=1 Δ1 /Δ2 ) showing very low levels of adipogenesis. Neither wt (G’) nor Δ1 /Δ2 (G’’) show large patches of Perilipin staining. (H) Representative examples from 3 injured wt muscles showing the range of adipogenesis that was observed. Colors of the numbers correspond to the mice shown in the dots on the wt BaCl2 bar of the graph in panel C to show how mean % adipogenesis values obtained for the different experimental animals correlate with the appearance of staining in tissues. The section numbered 1 and magnified panel (H’) shows the lowest levels of adipogenesis, with section 2 (H’’) showing higher levels. Section 3 shows the most adipogenesis throughout the section with large patches of adipocytes (H’’’). (I)Representative examples from 5 injured Δ1/Δ2 muscles showing the range of adipogenesis that was observed. Colors of the numbers correspond to the colored dots that represent different mice shown in the Δ1/Δ2 (del) BaCl2 bar of the graph in panel C to show how the different mean % adipogenesis values obtained for the different experimental animals correlate with the appearance of staining in tissues. Section number 1 and magnified panel (I’) shows the lowest levels of adipogenesis, Section 2 (I’’) shows higher levels but sections 3-5 display the highest Perilipin1 staining, with increasingly large areas of the section showing large regions of fat accumulation. Magnified insets show that large patches of adipocytes are found within the tissues (I’’’,I’’’’, I’’’’’). Error bars represent mean + s.d., EDL = Extensor Digitalis Longus (Scale Bars: 200 um (whole muscle sections), 50 um (insets))

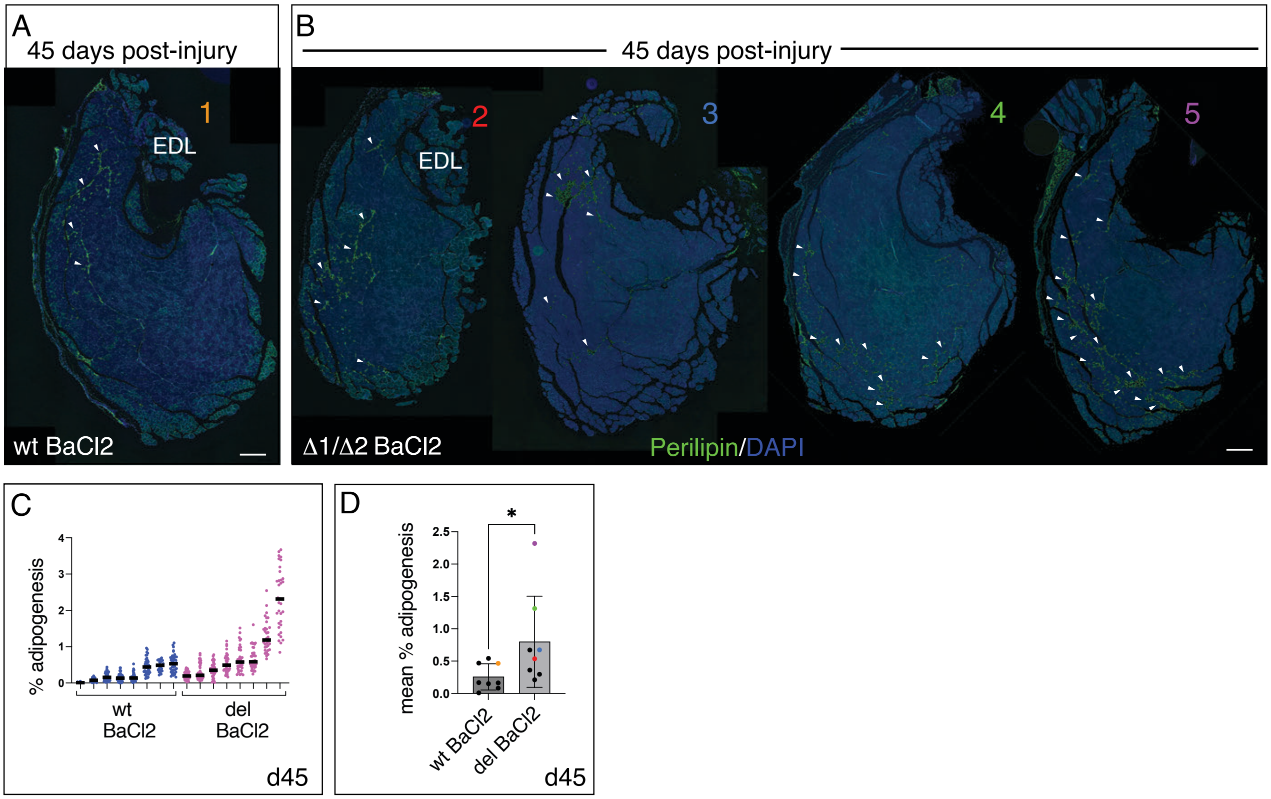
A
45 days post-injury
1
EDL
wt BaCl2
B
45 days post-injury
2
EDL
3
4
5
Δ1/Δ2 BaCl2
Perilipin/DAPI
C
% adipogenesis
wt
BaCl2
del
BaCl2
d45
D
mean % adipogenesis
*
wt BaCl2
del BaCl2
d45

Fig. S15

**Fig. S15: Adipogenesis in BaCl2 injured muscles at 45 days post-injury**

(A) A representative example of a section through the TA muscle of a BaCl2-injected wt muscle after 45 days. Section is stained with Perilipin 1 antibody to visualize adipocytes and DAPI to stain nuclei and provide a contour of the whole section. Arrowheads point to Perilipin stained regions containing adipocytes. Color of the number corresponds to a mouse shown in the graph in panel D to correlate the staining levels observed in this section with mean % adipogenesis values shown on the graph. (B) Representative examples of sections through the TA muscle of BaCl2-injected Δ1/Δ2 muscles after 45 days. Sections are arrayed on a single dark background separated by genotype and treatment for ease of viewing, but each section represents tissues from separate animals; muscles from four different animals are shown. The sections are stained with Perilipin 1 antibody to visualize adipocytes, and DAPI to stain nuclei and provide a contour of the whole section. Arrowheads point to Perilipin stained regions containing adipocytes. Colors of the numbers correspond to the mouse shown in the graph in panel D to correlate the staining levels observed in sections with mean % adipogenesis values obtained for the different experimental animals. (C) Scatter plots showing the individual % adipogenesis measurements for each section analyzed per animal to show the spread of values that was observed. There is a general upward trend in the measurements observed in Δ1/Δ2 muscles compared to wild type. Lines show the median. n= 8 wt BaCl2, n=8 Δ1/Δ2 (D) Bar graph to show the mean % adipogenesis values for individual wt vs. Δ1/Δ2 animals. Colored dots on the graph correspond to examples of numbered mouse tissue sections with the matching color shown in panels A and B to correlate the plotted mean % adipogenesis values with the staining levels observed in tissue. n=8 wt, n=8 Δ1 /Δ2 (*$P<0.05$, Mann-Whitney test) Error bars represent mean + s.d. EDL = Extensor Digitalis Longus muscle (Scale Bar: 200 um)

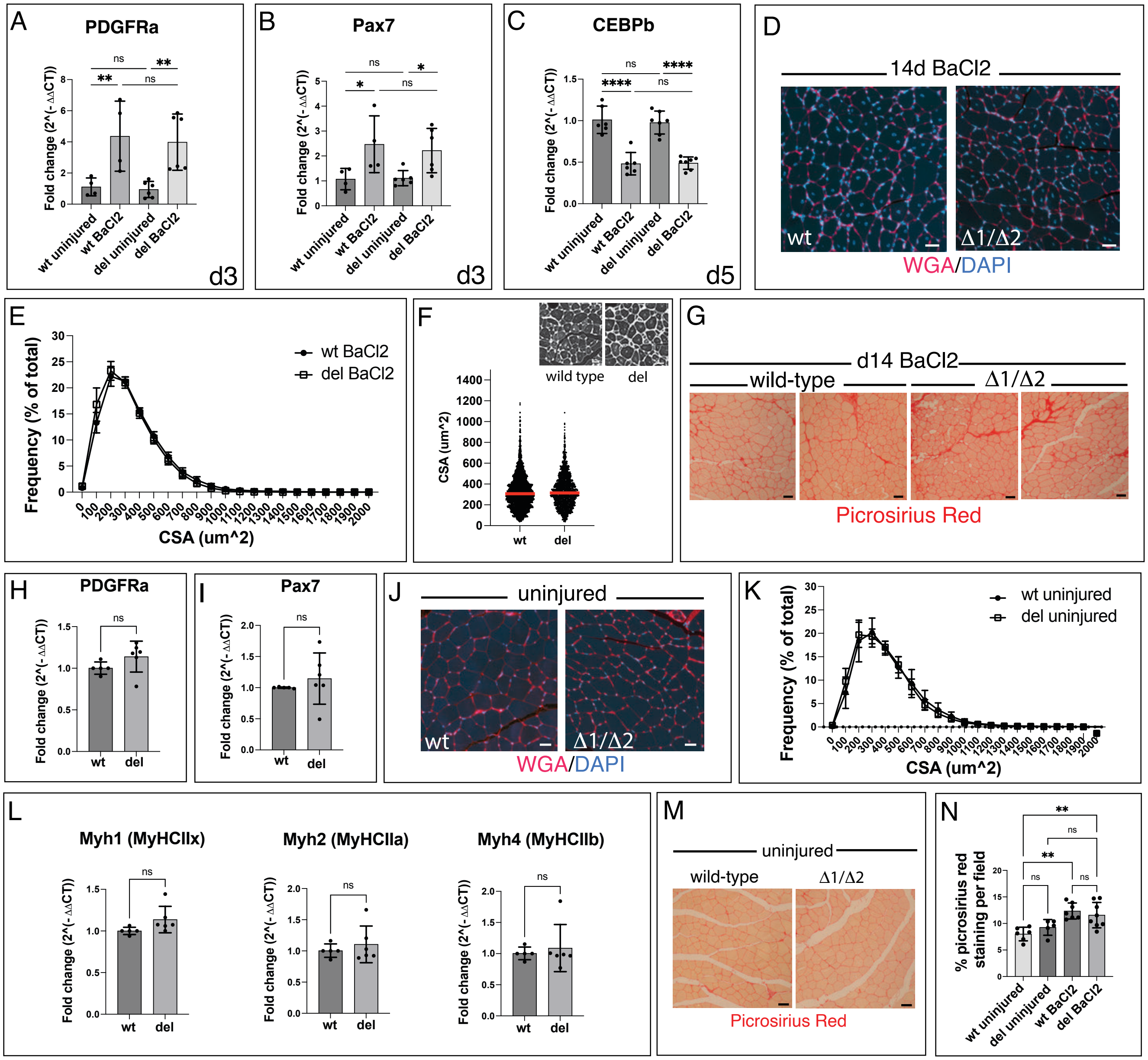
A
PDGFRa
Fold change (2^(-ΔΔCT))
wt uninjured
wt BaCl2
del uninjured
del BaCl2
d3
B
Pax7
d3
C
CEBPb
d5
D
14d BaCl2
wt
Δ1/Δ2
WGA/DAPI
E
wt BaCl2
del BaCl2
Frequency (% of total)
CSA (um^2)
F
wild type
del
CSA (um^2)
wt
del
G
d14 BaCl2
wild-type
Δ1/Δ2
Picrosirius Red
H
PDGFRa
I
Pax7
J
uninjured
wt
Δ1/Δ2
WGA/DAPI
K
wt uninjured
del uninjured
Frequency (% of total)
CSA (um^2)
L
Myh1 (MyHCIIx)
Myh2 (MyHCIIa)
Myh4 (MyHCIIb)
M
uninjured
wild-type
Δ1/Δ2
Picrosirius Red
N
% picrosirius red staining per field
wt uninjured
del uninjured
wt BaCl2
del BaCl2

Fig. S16

**Fig. S16: Comparison of wt and Δ1Δ/2 in injured and uninjured muscles**

(A-H, N) Assessment of responses of wt and Δ1 /Δ2 muscles to BaCl2 injury in injured mice. (A) RT-qPCR of *PDGFRa* expression following injury in wt and Δ1 /Δ2 mice showed significant up-regulation in injured muscles post-BaCl2 injection in both genotypes, but the differences between genotypes are not significant. Uninjured: n=4 wt, n=4 Δ1 /Δ2 , BaCl2 injured: n=6 wt, n=6 Δ1 /Δ2 . (** $p<0.01$, One-way ANOVA with Tukey's Multiple Comparison Test) (B) RT-qPCR of *Pax7* expression following injury in wt and Δ1 /Δ2 mice showed significant up-regulation in injured muscles post-BaCl2 injection in both genotypes, but the differences between genotypes are not significant. Uninjured: n=4 (wt), n=4 Δ1 /Δ2 , BaCl2 injured: n=6 wt, n=6 Δ1 /Δ2 . (* $p<0.05$, One-way ANOVA with Tukey's Multiple Comparison Test) (C) RT-qPCR of *CEBPb* expression following injury in wt and Δ1 /Δ2 mice showed significant down-regulation in injured muscles post-BaCl2 injection in both genotypes, but the differences between genotypes are not significant. Uninjured: n=6 wt, n=6 Δ1 /Δ2 , BaCl2 injured: n=7 wt, n= 7 Δ1 /Δ2 (**** $p<0.0001$, One-way ANOVA with Tukey's Multiple Comparison Test) (D) Representative examples of WGA stained muscle sections comparing BaCl2-injured wt vs Δ1 /Δ2 muscles at 14 days show no obvious differences in myofiber shape or size. n=8 wt, n=8 Δ1 /Δ2 (E) Cross sectional area measurements of injured wt vs Δ1 /Δ2 muscles show no difference between genotypes. n=8 wt, n=8 Δ1 /Δ2 (F) Violin plots showing the range of CSAs displayed by myofibers when only those that display centrally located nuclei are quantified. The range of measured CSA values and median (red line) are very similar between wt and Δ1 /Δ2 animals. wt = 2827 myofibers from n=5 animals, Δ1 /Δ2 = 1759 myofibers from n=4 animals (G) Picrosirius Red staining of wt vs Δ1 /Δ2 muscles at 14 days post-injury shows no obvious differences in the intensity of staining and collagen deposition in the muscle. Two representative examples of each genotype are shown. n=7 wt n=8 Δ1 /Δ2 (H-N) Responses of wt and Δ1 /Δ2 muscles to BaCl2 injury in uninjured mice. (H, I) RT-qPCR of *PDGFRa* (H) and *Pax7* (I) expression in uninjured muscles to compare basal levels between wt and

Δ1 /Δ2 mice show no significant difference between the genotypes. n=5 wt, n=6 Δ1 / Δ2 (*PDGFRa*, Unpaired t-test; *Pax7* Welch's t-test) (J) Representative examples of WGA-stained muscle sections of uninjured wt vs Δ1Δ/2 tissues show no obvious differences in myofiber shape or size. n=4 wt, n=5 Δ1 /Δ2 (K) Cross sectional area measurements of uninjured wt vs Δ1 /Δ2 muscles show no difference between genotypes. n= wt, n=8 Δ1 /Δ2 (L) RT-qPCR of *Myh1, 2* and *4* expression in uninjured muscles to compare basal levels between wt and Δ1 /Δ2 mice showed no significant difference between the genotypes. n=5 wt, n=6 Δ1 /Δ2 (*Myh1, Myh2* Unpaired t-test, *Myh4* Mann-Whitney test) (M) Picrosirius Red staining of wt vs Δ1 /Δ2 muscles show no obvious differences in the intensity of staining and collagen deposition in the muscle prior to injury. (N) Quantification of Picrosirius Red staining uninjured and injured wt vs Δ1 /Δ2 muscles show no difference in the deposition of collagen between genotypes in the uninjured state, or at 14 days post-injury, although collagen deposition is increased in injured muscles compared to uninjured samples. Uninjured: n=6 wt, n=5 Δ1 /Δ2 , BaCl2 injured: n=7 wt, n=8 Δ1 /Δ2 (One-way ANOVA with Sidak's Multiple Comparison's Test) Error bars represent mean + s.d. (RT-qPCR); mean + s.e.m (histograms) (Scale bars: 20 um)

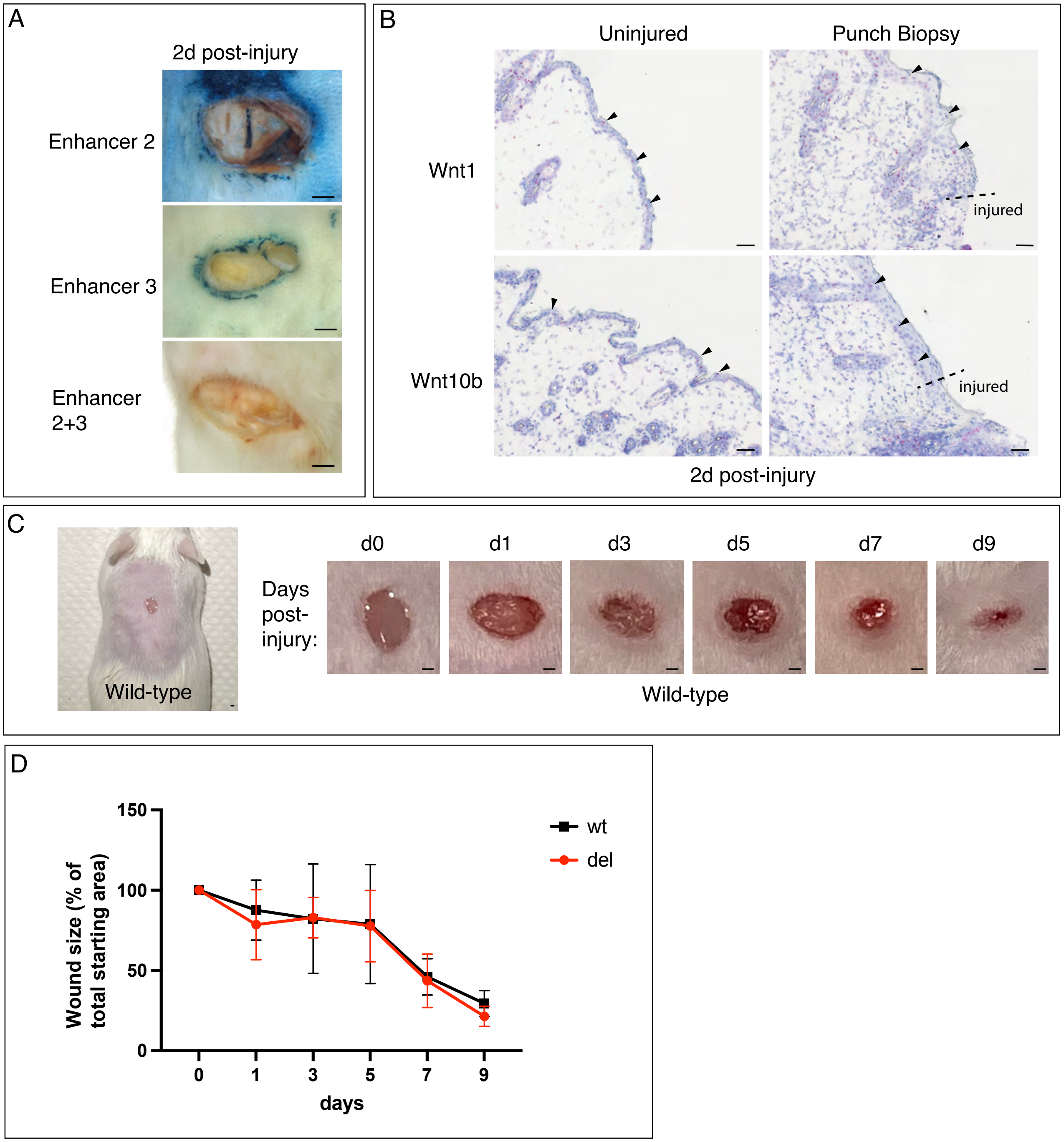
A
2d post-injury
Enhancer 2
Enhancer 3
Enhancer 2+3
B
Uninjured
Punch Biopsy
Wnt1
Wnt10b
injured
injured
2d post-injury
C
Wild-type
Days post-injury:
d0
d1
d3
d5
d7
d9
Wild-type
D
Wound size (% of total starting area)
150
100
50
0
0
1
3
5
7
9
days
wt
del

Fig. S17

**Fig. S17: Mice lacking Enhancers 2 and 3 display no defects in wound healing of the skin**

(A) Expression of Enhancer 2, 3, and 2+3-LacZ in skin following punch biopsy at 2 days post-injury. Both Enhancers 2 and 3 reporters stain with X-gal, but Enhancer 2+3 does not express the reporter. (B) mRNA in situ hybridization showing that expression of *Wnt1* and *Wnt10b* is not observed near the injury site following punch biopsy. The location of the punch biopsy is labeled as ‘injured’ and the boundary of the wound edge is marked with a dotted line. The skin exhibits staining of both *Wnt1* and *Wnt10b* (arrowheads) in the basal layer, and some thickening of the skin appears post-injury in the region flanking the wound, but no obvious elevation in mRNA signals at the wound edge is detected. (C) An example of a dorsal punch biopsy wound in a wild-type mouse and the changes in wound size that are observed over 9 days of recovery. (D) Wound closure measured over 9 days shows that there is no significant difference in healing between wt and $\Delta1/\Delta2$ animals. (n=4 wt, n=4 $\Delta1/\Delta2$; wt vs $\Delta1/\Delta2$ was compared at each time point: Unpaired t-test (1 and d3), Welch’s t-test (d3, d5), Mann-Whitney t-test (d9). Error bars represent mean + s.d. Scale bar: 1mm (A, C), 50um (B)

### Table S1: Summary of Wnt expression in 4 injury models

| Tissue | Lung (Naphthalene, d3) | | Muscle (BaCl2, d3) | | Liver (CCl4, d3) | | Pancreas (STZ, d7) | |
|---|---|---|---|---|---|---|---|---|
| | Uninjured | Injured | Uninjured | Injured | Uninjured | Injured | Uninjured | Injured |
| Wnt1 | - | + | - | + | +/- | +/- | + | + |
| Wnt2 | - | - | - | + | ++ | +++ | +/- | + |
| Wnt2b | +/- | +/- | - | + | - | +/- | + | + |
| Wnt3 | - | - | - | + | - | +/- | - | - |
| Wnt3a | + | +/- | - | +/- | - | +/- | + | + |
| Wnt4 | ++ | ++ | +/- | + | + | ++ | ++ | +++ |
| Wnt5a | ++ | ++ | +/- | + | ++ | ++ | + | + |
| Wnt5b | +/- | - | - | + | + | + | + | + |
| Wnt6 | +/- | +/- | - | +/- | - | + | + | + |
| Wnt7a | +/- | + | - | + | + | + | + | +/- |
| Wnt7b | + | ++ | - | + | - | +/- | + | + |
| Wnt8a | - | - | - | + | - | + | +/- | +/- |
| Wnt8b | - | -/+ | - | + | + | + | + | + |
| Wnt9a | ++ | ++ | + | ++ | + | + | + | + |
| Wnt9b | +/- | +/- | - | ++ vessels | ++ | +++ | + | + |
| Wnt10a | - | - | - | + | +/- | + | +/- | - |
| Wnt10b | -/+ | + | - | + | +/- | + | + | + |
| Wnt11 | ++ | ++ | + | ++ | +/- | ++ | + | + |
| Wnt16 | - | - | - | - | - | - | + | +/- |

Naphthalene (N= >2 mice per treatment condition); BaCl2 = Barium Chloride (N=2 mice per treatment condition), CCl4 = Carbon Tetrachloride (N=2 mice per treatment condition); STZ = Streptozotocin (N=1 mouse per treatment condition)

Scoring:
Lung: (only staining within airways or directly underlying airways was scored, cells in the alveolar compartment are marked by white asterisks) - = no signal; +/- = rare or weak signals; + = single nuclear spots observed; ++ = nuclei with multiple spots of staining

Muscle: - = no signal; +/- = signals are rare or weak, + = single nuclear spots observed, ++ = multiple spots per nucleus observed

Liver: - = no signal; +/- = rare nuclei with single spots; + = readily observable nuclei with single spots; ++ = nuclei with multiple spots of staining; +++ = many cells with nuclei with multiple spots of staining, or dense staining.

Pancreas (only signals in islets were scored): - = no signal; +/- = signals are rare or weak, + = single nuclear spots found in islets, ++ = multiple spots per nucleus seen in islets, +++ = dense staining

**Table S2: Expression of Injury responsive reporters in muscle**

| Reporter | Timepoint | Expression Pattern | Expression |
|---|---|---|---|
| Enh 2, uninjured | 6h, 12h, d1, d3, d5 | None * | - |
| Enh 2 | 6h | None | - |
| Enh 2 | 12h | Speckled distribution (whole mount) | + |
| Enh 2 | d1 | Speckled distribution (whole mount), individual cells residing between myofibers (section) | ++ |
| Enh 2 | d3 | Uniform staining throughout tissue (whole mount), individual cells residing between myofibers, staining in regenerating myofibers of varying intensity (section) | ++++ |
| Enh 2 | d5 | Individual cells residing between myofibers, staining in regenerating myofibers of varying intensity (section) | +++++ |
| Enh 3, uninjured | 6h, 12h, d1, d3, d5 | None | - |
| Enh 3 | 6h | Speckled distribution (whole mount), individual cells residing between myofibers (section) | +++ |
| Enh 3 | 12h | Speckled distribution (whole mount), individual cells residing between myofibers (section) | +++ |
| Enh 3 | d1 | Speckled distribution (whole mount), individual cells residing between myofibers (section) | +++ |
| Enh 3 | d3 | Speckled distribution (whole mount), individual cells residing between myofibers (section) | +++ |
| Enh 3 | d5 | Individual cells residing between myofibers, staining in uninjured myofibers starts to appear (considered background) (section) | + |
| Enh 2+3, uninjured | 6h, d1, d3, d5 | None | - |
| Enh 2+3 | 6h | None | - |
| Enh 2+3 | 12h | NA | NA |
| Enh 2+3 | d1 | Speckled distribution (whole mount) | ++ |

| | | | |
|---|---|---|---|
| Enh 2+3 | d3 | Speckled distribution (whole mount), individual cells residing between myofibers (section) | ++ |
| Enh 2+3 | d5 | Staining in cells residing between myofibers is extremely rare, no staining in regenerating myofibers. | - |

(*Note: Occasional uninjured myofibers show staining but this is considered non-specific background and is can appear in lines carrying the hsp68-LacZ construct; number and intensity varies between tissues, this pattern can be seen in all specimens, both uninjured and injured muscles and is therefore not included in the assessment we present here)

**Table S3: Reporter activity of AP-1 mutant lines compared to wild-type Enhancer 3 -LacZ**

| Reporter Line | Naphthalene injury | | | | | CCl4 injury | | Punch Biopsy |
|---|---|---|---|---|---|---|---|---|
| | lung | lung | lung | lung | trachea | liver | liver | skin |
| | 12h | 1d | 3d | 5d | 12h | 1d | 1d | 1d |
| | whole mount | section | section | section | section | whole mount | section | Whole mount |
| Enhancer 3 (Wild type) Uninjured | - | - | - | - | - | - | - | - |
| Enhancer 3 (Wild type) | ++++ | +++ | +++ | ++ | + | ++++ | +++ | ++++ |
| AP-1 mut. Uninjured* | - | - | - | - | - | - | - | - |
| AP-1 mut. #21 | + | ++ | + | +++ | - | - | - | - |
| AP-1 mut. #27 | - | - | - | - | + | - | - | - |
| AP-1 mut. #39 | - | - | + | + | - | - | - | + |
| AP-1 mut. #44 | + | + | + | + | ++ | - | - | + |
| AP-1 mut. #14 | ++ | + | + | + | ++ | + | + | + |
| AP-1 mut. #22 | - | - | + | + | - | - | - | - |
| AP-1 mut. #32 | - | - | - | - | - | - | + | - |
| AP-1 mut. #37 | + | + | + | + | ++ | - | + | ++ |

*AP-1 mut. Uninjured: A single entry summarizes the expression in these controls.  In the absence of injury, none of the lines showed lacZ staining in lung, liver, and skin.
